# Enhancing CAR T cell therapy using Fab-Based Constitutively Heterodimeric Cytokine Receptors

**DOI:** 10.1101/2023.04.24.538039

**Authors:** Matteo Righi, Isaac Gannon, Matthew Robson, Saket Srivastava, Evangelia Kokalaki, Thomas Grothier, Francesco Nannini, Christopher Allen, Bai Yuchen, James Sillibourne, Shaun Cordoba, Simon Thomas, Martin Pule

## Abstract

Adoptive T cell therapy aims to achieve lasting tumour clearance, requiring enhanced engraftment and survival of the immune cells. Cytokines are paramount modulators of T cell survival and proliferation. Cytokine receptors signal via ligand-induced dimerization, and this principle has been hijacked utilising non-native dimerization domains. A major limitation of current technologies resides in the absence of a module that recapitulates the natural cytokine receptor heterodimeric pairing. To circumvent this, we created a new engineered cytokine receptor able to constitutively recreate receptor-heterodimer utilising the heterodimerization domain derived from the IgG1 antibody (dFab_CCR). We found that the signal delivered by the dFab_CCR-IL2 proficiently mimics the cytokine receptor heterodimerization, with transcriptomic signatures similar to that obtained by the activation of IL2 native receptor. Importantly, we found that this dimerization structure is agnostic, efficiently activating signaling through four cytokine receptor families.

Using a combination of *in vivo* and *in vitro* screening approaches, we characterized a library of 18 dFab_CCRs co-expressed with a clinically relevant solid tumor-specific GD2 CAR. Based on this characterization we suggest that the co-expression of either the common β-chain GMCSF or the IL18 dFab_CCRs is optimal to improve CAR T cell expansion, engraftment, and efficacy.

Our results demonstrate how the Fab dimerization is efficient and versatile in recapitulating a cytokine receptor heterodimerization signal. This module could be applied for the enhancement of adoptive T cell therapies, and therapies based on other immune cell types. Furthermore, these results provide a choice of cytokine signal to incorporate with adoptive T cells therapies.

## INTRODUCTION

The full activation and proliferation of adoptively transferred immune cells (e.g. TILs, TCR T cells and CAR T cells) require multiple immunologic signals (1). In addition to antigen receptor signaling, cytokine receptor activation directs and amplifies T cell differentiation and expansion (2). However, within the tumor microenvironment, immunostimulatory cytokines are limited (3). Furthermore, in conditions of minimal residual disease, or prolonged trafficking to sites of disease, the scarcity of antigen receptor activation limits T cell auto- and paracrine cytokine production, additionally restricting T cell cytokine activation (4).

Several approaches have been explored in adoptive T cell therapy to overcome this lack of endogenous immunostimulatory cytokines. Cytokines such as IL2 can be given systemically, however side-effects are common and prolonged administration is impractical (5,6). T cells can be engineered to secrete cytokines, however this approach is also associated with systemic adverse effects (7,8).

Alternative approaches have focused on engineering cytokine receptors to deliver cytokine signals in the absence of cytokines. Some engineered cytokine receptors signal in response to a pharmaceutical agent (9), or ubiquitous and inhibitory microenvironmental factors (10,11). In other approaches, cytokine receptors are engineered to be constitutively active without the requirement for exogenous stimuli (12,13).

We add to the literature of constitutively active cytokine receptors with a novel format which forces cytokine receptor heterodimerization by replacing cytokine receptor ectodomains with antibody Kappa Constant Light Chain (KCL) and CH1 domain (dFab_CCR). Expressed in human T cells, we show that the dFab_CCR design is highly versatile, being able to transmit any major cytokine receptor family signal. Such flexibility allowed for the characterization of several dFab_CCRs that enhanced CAR T cell effector function and persistence both *in vitro* and *in vivo*. We identified IL18 and GMCSF dFab_CCR as multifaceted modulators of CAR T cells effector function by improving persistence, enhancing cytokine secretion and augmenting aerobic metabolism. This translated into enhanced anti-tumor activity in several animal models.

## STAR Methods

### Retroviral and Plasmid Constructs

Molecular cloning was performed by Golden Gate assembly using *Bsa I*, *BsmB I*, *Bbs I* restriction enzymes depending on the DNA sequence. G-blocks encoding for cytokine receptors sequences, constant heavy chain 1, constant light kappa chain and constant light lambda chain were purchased from IDT. Codon optimization was performed using GeneArt Codon Optimization Tool that strove to keep GC content at 60% and eliminate cryptic splicing, hairpins, literal repeats, and any possible *cis*-acting sequences. Each open reading frame was cloned into the splicing oncoretroviral vector SFG, as polycistronic nucleotide sequences separated by interposed foot-and-mouth disease (FMD)-2A TaV motifs, codon wobbled to prevent retroviral recombination. The human GD2 CAR was composed of the Huk666 scFv (14), a CD8 alpha stalk and trans-membrane domain along with a 41bb-CD3ζ endodomain. The human CD19 CAR comprised the fmc63 scFv (15), a CD8 alpha stalk and a trans-membrane domain along with a 41bb-CD3ζ endodomain. The PSMA CAR was composed of the 10C11 scFv (in house), a CD8 alpha stalk and a trans-membrane domain along with a 41bb-CD3ζ endodomain.

The dFab_CCRs were cloned utilizing the Constant Heavy 1 region and the Constant Light region of the human IgG antibody. The transmembrane and endodomain were derived from the indicated cytokine receptor. The murine dFab_CCRs were generated utilizing the indicated murine cytokine receptor transmembrane and endodomain. All human constructs were co-expressed with RQR8 (16) as a surrogate of CAR gene expression.

The murine GD2 CAR comprises the muk666 scFv, a murine CD8 alpha stalk and trans-membrane domain along with a murine CD28 and CD3ζ endodomains (17). The murine constructs were co-expressed with THY1.1 as surrogate of CAR expression.

### Cell lines

HEK-293T, Phoenix ECO, SKOV-3 GD2^+^, SKOV-3 mKATE/CD19^+^ and CHLA-255 cells were cultured in I10 medium consisting of Iscove’s modified Dulbecco’s medium (IMDM, Gibco) supplemented with 10% fetal bovine serum (FBS, HyClone, Thermo Scientific) and 2 mM GlutaMAX (Sigma). SupT1 NT, SupT1 GD2^+^, CT26 NT, CT26 GD2^+^, B16F10 NT, B16F10 GD2^+^, NALM6 and NALM6-CD19KO cell lines were cultured in R10 medium consisting of RPMI (RPMI-1640, Gibco) supplemented with 10% FBS (HyClone, Thermo Scientific) and 2 mM GlutaMAX. HEK-293 T, Phoenix Eco, SupT1, CT26, SKOV3 and B16F10 lines were obtained from the American Type Culture Collection.

CHLA-255 cells were kindly donated by from COG Cell Culture Core + Xenograft Repository, Texas Tech Uni HSC Cancer. SupT1, SKOV3, CT26 and B16F10 were transduced with an SFG vector to express human GD2. SKOV3 were transduced with an SFG vector to express human CD19 and red fluorescent protein mKATE. CHLA-255 and NALM6 were transduced with an SFG vector to express firefly luciferase.

### Retroviral transduction of primary human T cells

RD114-pseudotyped γ-retroviral supernatants were generated by transient transfection of HEK-293T cells with an SFG transfer vector plasmid (4.7 µg), an RD114 envelope expression plasmid (RDF, a gift from M. Collins and Y. Takeuchi, University College London, 3.2 µg), and a packaging plasmid expressing MoMLV Gag-pol (a gift from E. Vanin, Baylor College of Medicine, 4.7 µg). Transfection was facilitated using GeneJuice (Merck Millipore), following manufacturer’s instructions.

Leucocyte cones of healthy donors were purchased from National Health Service Blood and Transplant (NHSBT, UK). Whole blood was extracted from each cone and diluted to 50mL with sterile PBS. PBMCs were isolated by Ficoll gradient centrifugation using SepMate 50 (StemCell) layering 25 mL of whole blood mixture to each SepMate 50. The cells were centrifuged at 1200 *g* for 20 minutes. The buffy coat was extracted and washed twice with sterile PBS. Isolated PBMCs cells were resuspended at 1x10^6^ cells/mL in R10 and stimulated with 50 ng anti-CD3 (Miltenyi) and 50 ng anti-CD28 (Miltenyi) per 1 x 10^6^ cells. 100 iU/mL IL2 (GenScript) was added following overnight stimulation. 24 hours after IL2 supplementation, cells were collected, plated at a density of 1 x 10^6^ cells per well (1 mL) on retronectin-coated (Takara) 6-well plates with 3 mL of retroviral supernatant in the presence of 100 iU/mL of IL2, and centrifuged at 1,000 *g* for 40 min. 24 hours post spinoculation, the T cells were harvested and re-plated in complete R10 media supplemented with 100 iU IL2. Transduction efficiency was determined on day 5 after transduction, and further experiments were commenced on days 5–9 after transduction.

### Retroviral transduction of murine T cells

Ecotropic Virus was prepared by transiently transfecting Phoenix-Eco (PhEco, RRID:Addgene_12371) adherent packaging cells with an SFG vector (4.7 µg) and pseudotyped with ecotropic envelop vector pCL-ECO (2.68 µg). Transfection was facilitated using GeneJuice (Merck Millipore), following manufacturer’s instructions.

Splenocytes were extracted as follows: Spleens were pulped and passed through a 70 μm cell strainer. Red blood cells were lysed using ACK red blood cell lysis buffer (Gibco) for 5 min at room temperature (RT). The reaction was stopped with 15mL of Hank’s Balanced Salt Solution (HBSS, Gibco). After the cells were pelleted, the splenocytes were passed through a 40 μm cells strainer to obtain a single cell suspension. Isolated splenocytes were activated with 2 µg Concanavalin A (Sigma) and 1 ng murine IL7 (Miltenyi) per 1.5 x 10^6^ cells for 24 hours. 5 x 10^6^ cells were plated on retronectin-coated (Takara) 6-well plates in 2 mL of retroviral supernatant, spun 800 *g* for 90 minutes. The wells were supplemented with 4 mL of complete media R10 incubated for 72 hours at 37°C in 5% CO_2_, in the presence of human IL2 (100 iU/mL; GenScript).

### Transduction efficiency evaluation

Staining steps were performed at RT for 12 minutes, with PBS washes between steps. Cells were co-stained with either eFluor 780 or eFluor450 fixable viability dye (eBioscience) depending on the experimental condition. Transduced human T cells were detected by staining for the RQR8 marker, utilizing anti-CD34 antibody. Murine CAR T cell transduction was evaluated by staining for THY1.1 marker. The samples were stained in 96 well plates and resuspended in 100 μL of PBS. Flow cytometry was performed using the MacsQuantX flow cytometer (Miltenyi), running 50 μL of cell suspension. Data analysis was conducted using FlowJo v10 (Treestar, RRID:SCR_008520).

### T cell Immunophenotyping

Staining steps were performed at RT for 12 min, with PBS washes between steps. Transduced cells were identified by RQR8 positivity utilising a CD34-specific antibody, and live cells selected via Sytox live dead cell dye exclusion. Memory, exhaustion, and activation phenotypes in CD4^+^ and CD8^+^ CAR T cells were based on the expression of CD45RA, CCR7, PD-1, LAG-3, TIM-3, KLRG-1, CD25, CD69 and CD95.

The samples were stained in 96 well plate and resuspended in 100 μL of PBS. Flow cytometry was performed using the MacsQuantX flow cytometer (Miltenyi), running 50 μL of cell suspension. Data analysis was conducted using FlowJo v10 (Treestar, RRID:SCR_008520).

### Animal sample phenotyping

Animals were sacrificed by Isoflurane intoxication followed by cervical dislocation. Spleens were processed as described for the murine T cell generation. Tumor infiltrating lymphocytes were enriched from tumour biopsies by using Tumor Dissociation Kit, murine (Miltenyi) and processed utilising gentleMACS™ Dissociator (Miltenyi). Erythrocytes were lysed using 500 μL of ACK lysis buffer (Sigma) for 5 minutes at RT. Cells were strained twice, first utilising a 70 μm cell strainer then a 40 μm cell strainer. 2 x 10^6^ cells were aliquoted for phenotyping by flow cytometry. First, the Fc receptor was blocked using anti-CD32/CD16 (BioLegend) to avoid non-specific binding. Transduced CAR T cell populations were identified based on the expression of CD45, CD3, CD4, THY1.1 and the absence of CD11b. The identification of murine T cell memory subpopulation were performed by the detection of CD62L and CD44. Myeloid population was defined by the expression of CD45 and CD11b. Monocytes were identified by the expression of F4/80. Dedritic Cells (DC) were identified by the expression of CD11b..

The samples were stained in 96 well plates and resuspended in 100 μL of PBS. Flow cytometry was performed using the MacsQuantX flow cytometer (Miltenyi), running 50 μL of cell suspension. Data analysis was conducted using FlowJo v10 (Treestar, RRID:SCR_008520).

### Intracellular staining

Between 1 x 10^5^ to 5 x 10^4^ cells/well were pelleted by centrifugation at 1000 *g* for 2 minutes at room temperature. The cells were stained with eFluor450 fixable viability dye (Biolegend), CD3 (Biolegend) and CD34 (Biolegend) for 10 minutes at RT. The cells were washed once with PBS and pelleted 1000 *g* for 2 minutes. The cells were fixed and permeabilized according to the manufacturer’s protocol using BD Cytofix/Cytoperm™ Fixation/Permeabilization Solution Kit (Fisher). The cells were then stained with anti-Human Kappa Light Chain Biotin (Thermo) in permeabilization/washing solution for 12 minutes at RT, washed once with permeabilization/washing solution and pelleted by centrifugation at 1000 *g* for 2 minutes. The cells washed once more with 200 μL of permeabilization/washing solution and pelleted as before.

The cells were then stained with PE-conjugated Streptavidin (Biolegend) in permeabilization/washing solution for 12 minutes at RT The samples were stained in 96 well plates and resuspended in 100 μL of PBS. Flow cytometry was performed using the MacsQuantX flow cytometer (Miltenyi), running 50 μL of cell suspension. Data analysis was conducted using FlowJo v10 (Treestar, RRID:SCR_008520).

### Human CAR T cells FACS based cytotoxicity assay co-culture

CAR transduced T cells were normalized based on RQR8-expression by dilution with non-transduced T cells. Target and effector cells were resuspended in R10 media and plated in a 96-well plate flat bottom at the desired volume to achieve the desired effector:target ratio. Target cell number was kept constant within each experiment, at 5 × 10^4^ cells. After 48 hours, the plate was centrifuged at 1000 *g* for 2 min, 100 μL of supernatant was removed for cytokine analysis and CountBright beads (Thermo Fisher) were added to allow normalization of cell numbers acquired. T cells were distinguished from target cells by positivity for CD2 and CD3. Gating on single live target cells was performed according to exclusion of fixable viability dye eFluor780. Flow cytometry was performed using the MacsQuantX flow cytometer (Milteny). Data analysis was conducted using FlowJo v10 (Treestar, RRID:SCR_008520). Percentage of live cells was calculated relative to the number of live target cells after co-culture with non-transduced T cells.

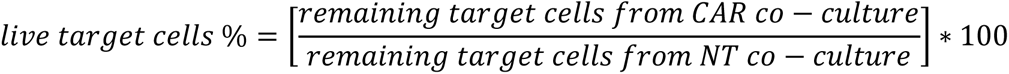

### Murine CAR T cells FACS based cytotoxicity assay co-culture

Target cells were plated 24 hours prior to the assay at a density of 2.5 x 10^4^ cells / well in a 96-well plate flat bottom. Murine CAR transduced T cells were normalized based on THY-expression by dilution with non-transduced T cells and plated over the target cells to achieve the desired effector:target (E:T) ratio. After 48 hours, the plate was centrifuged at 1000 *g* for 2 minutes, supernatant was removed for cytokine analysis and CountBright beads (Thermo Fisher) were added to allow normalization of cells numbers acquired. Target cells were detached from the plate using 50 μL of Trypsin (Gibco) for 5 minutes at 37°C. The reaction was stopped with 100 μL of R10 and the cells were collected in a new 96-well plate. Following a wash with PBS the cells were ready to stain.

Murine T cells were distinguished from target cells by positivity for CD3 and CD45. Gating on single live target cells was performed according to exclusion of fixable viability dye eFluor780. Flow cytometry was performed using the MacsQuantX flow cytometer (Milteny). Data analysis was conducted using FlowJo v10 (Treestar, RRID:SCR_008520). Percentage of live cells was calculated relative to the number of live target cells after co-culture with non-transduced T cells.

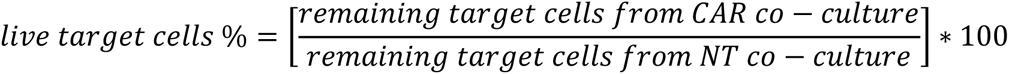

### Cytokine Starvation assay

CAR T cells were pre-labelled with Cell Trace Violet (ThermoFisher) according to manufacturer procedure. Transduced T cells were gated via detection of CD3 and CD34, whereas non-transduced T cells were gated via CD3 only. If ruxolitinib was used, T cells were pre-incubated with either 1 µM or 10 µM of ruxolitinib for 4 hours, then washed to remove excess ruxolitinib. 1 x 10^5^ cells were cultured for either 4 or 7 days in R10 media without the addition of exogenous cytokines in a 96-well plate flat bottom. Murine T cells were plated at 1 x 10^5^ cells in a 96-well plate flat bottom, without CTV pre-labelling. CTV Mean Fluoresced value was calculated, gating on single live target cells was performed according to exclusion of fixable viability dye eFluor780 and positivity for RQR8 (Human CAR T Cells) or THY1.1 (murine CAR T cells). Flow cytometry was performed using the MacsQuantX flow cytometer (Milteny). Data analysis was conducted using FlowJo v10 (Treestar, RRID:SCR_008520). The T fold expansion was calculated as follow:

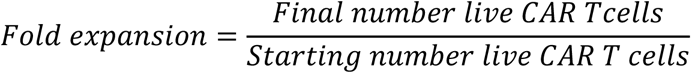

### Serial Tumor re-challenge Assay

1 x 10^6^ CAR T cells and 1 x 10^5^ of either SKOV-3-GD2^+^ cells or SKOV-3-CD19^+^ cells E:T ratio of 10:1, were co-cultured in cytokine free R10 media for 7 days in 24-well plate. After 7 days, the CAR T cells were carefully harvested, and the viable CAR T cells were enumerated. 1 x 10^6^ live CAR T cells were re-plated over fresh 1 x 10^6^ SKOV-3 to reach the E:T ratio of 10:1. The co-culture was incubated for further 7 days. The cycle of stimulations was performed for a total of three re-challenging.

### Serial Starvation Assay

0.5 x 10^6^ CAR T cells were cultured in cytokine free R10 media for 7 days in 24-well plate. After 7 days, the CAR T cells were carefully harvested, and the viable T cells were enumerated. Up to 0.5 x 10^6^ CAR T cells were re-plated and cultured for further 7 days. The cycle of starvation was continued until the CAR T cells number at the day of the harvesting fell below the counter limit of detection (0.05 x 10^6^). The data are plotted as 10^6^ cumulative numbers, calculated as the sum of 10^6^ viable CAR T cells every 7 days.

### Activation of either GD2-Specific or CD19-specific CAR-T Cells with scFv Anti-idiotype antibodies

10 μg of α-huk666 idiotype (for GD2CAR T Cells) or 1 μg of α-fmc63 idiotype (for CD19 CAR T cells) were coated on non-tissue culture treated 24-well plates for 16 hours at 4°C. Plates were washed twice before plating T cells (1 x 10^6^ cells/well). Plates were centrifuged at 500 *g* for 5 minutes and incubated at 37°C for 24 to 48 hours.

### Analysis of cytokine production

IFNγ, IL2 were detected utilising a specific Biolegend ElisaMax kit, following manufacturer instructions. Cytokine bead array (Biolegend, Legend Plex custom-made) was performed according to manufacturer’s protocol. The samples were detected using Fortessa LRS (BD) and analyzed using LEGENDplex™ Data Analysis Software Suite (Biolegend)

### RNA sequencing

The transduced T cells were isolated using CD34 microbeads (Miltenyi) and rested for 24 hours in R10 media. Subsequently, the T cells were seeded at 1 x 10^6^ cells/mL with or without 100000 iU/mL of HuIL2 (GeneScript). After 72 hours, rHuIL12 was replenished. After 7 days the cells were lysate and the mRNA was extracted with the illustra™ RNAspin Mini RNA Isolation Kit (GE Healthcare). The RNA was quantified with NanoDrop (ThermoFisher). The quality of the RNA was assessed with Agilent TapeStation (Agilent, RRID:SCR_019547). Roughly 40 million paired end reads (150 bp) were sequenced on Illumina Hiseq sequencer (GeneWiz) for each individual CAR-T sample with different CCR modules.

Fastq reads were mapped to the human genome (B37.3) and gene model GenCode (V19) with Omicsoft Aligner 4 (18). Gene count data was normalized by logGenometric mean values. Differential gene expression analysis was carried out by using DeSeq2 General Linear Model test (RRID:SCR_015687) (19). Geneset Enrichment Analysis (Subramanian) Pre-ranked tool (GSEAPreranked) was used to conduct the pathway and functional analysis on the differentially expressed genes.

### Nanostring

Total mRNA was collected using the PureLink RNA Mini Kit (ThermoFisher). Gene expression analysis was performed using the CAR-T cell Panel version 1 (NanoString, Seattle, WA) and RNA Profiling Core using the nCounter Analysis System. Data were analyzed using nSolver 4.0 software (NanoString).

### Analysis of Metabolic Parameters

Mitochondrial function was assessed with the extracellular flux analyzer (SeahorseBioscience). Individual wells of a XF96 cell-culture microplates were coated with CellTak following manufacturer’s instructions. The matrix was adsorbed overnight at 37°C. Mitochondrial function was assessed 24 hours post-activation or 24 hours post-starvation. Regarding the GD2 CAR T cells, 1.25 x 10^5^ cells were resuspended in XF assay medium (non-buffered RPMI 1640) containing 5.5 mM glucose, 2 mM L-glutamine, and 1 mM sodium pyruvate. Regarding the CD19 CAR experiments, 5 x 10^4^ cells were resuspended in XF assay medium (non-buffered RPMI 1640) containing 5.5 mM glucose, 2 mM L-glutamine, and 1 mM sodium pyruvate.

The microplate was centrifuged at 1,000 *g* for 5 minutes and incubated in standard culture conditions for 60 minutes. During instrument calibration (30 minutes), the XF96 assay cartridges were calibrated in accordance with the manufacturer’s instructions. Cellular OCRs were measured under basal conditions and, following treatment with 1.5 mM oligomycin, 1.5 mM FCCP, and 40 nM rotenone, with 1 mM antimycin A (XF Cell Mito Stress kit, Seahorse Bioscience).

### Barcode analysis

Genomic DNA was isolated using PureLink™ Genomic DNA Mini Kit (ThermoFisher). Barcode sequence was amplified via PCR using the forward primer (ACACTCTTTCCCTACACGACGCTCTTCCGATCTCCGCCAACGCGGTCGCAC) and The reverse primer(GACTGGAGTTCAGACGTGTGCTCTTCCGATCTGATGAGAACAGTATCGATTAGGGTTGACGGC) at 65°C amplification temperature. PCR products were isolated from a 3% agarose gel. The purified PCR product were sequenced using Amplicon-EZ next generation sequencing service (GeneWiz). Roughly 50,000 reads per sample were fully sequence with 2 x 250 bp sequencing. The contigs of paired-end reads were created and Fastq quality performed to exclude disastrous reads. The pair-end read were aligned together and the barcode sequence reads counted. Values normalized to the total reads obtained in each sample.

### Phosphorylated protein Immunoblot analysis

1 x 10^6^ cells were harvested and pelleted at 5000 *g* for 2 minutes. The samples were washed with of ice-cold PBS and the cells were pelleted at 5000 *g* for 2 minutes. The pellet was lysate in 50 µL of RIPA buffer (Millipore) supplemented with protease and phosphatase inhibitor (Abcam, ab201119). The cells were incubated in ice for 15 minutes. The tubes were then vortexed and incubated for further 15 minutes. The lysate was clarified by centrifuging the cells at max speed for 10 minutes at 4°C. The lysate was then transferred in PCR tube strips with 12.5 μL of NuPAGE™ LDS Sample Buffer (4X) (Invitrogen™, NP0007). The sample was then boiled at 95°C for 5 minutes. 8 μL of each sample was run on a premade 4–20% Mini-PROTEAN® TGX™ Precast Gel (Abcam), at 180V, 500 mA for 37 minutes on constant voltage mode. 5 μL of protein ladder was run as a size marker.

Post-separation, SDS-PAGE gel was transferred to Trans-Blot Turbo Midi 0.2 µm PVDF membrane (Abcam, 1704157) using Trans-Blot® Turbo™ Midi by semi-dry transfer (Abcam). Following transfer, membranes were blocked with 1x TBST (Sigma) supplemented with 5% BSA for 24 hours at 4°C. Membrane staining was undertaken with antibody diluted to appropriate concentration in 1x TBST supplemented with 5% BSA for 1 hour on orbital shaker, 480 rpm, at RT. The sample was then washed 3 times with 1x TBST for 10 minutes each time. The secondary staining was performed as per primary staining. Following the last TBST was, the membrane was washed with distilled water, then the membrane developed by incubation with Pierce ECL Plus Western Blotting Substrate (Abcam), according to the manufacturer’s instructions.

### Study approval

All procedures in this study gained the approval of The Animal Welfare and Ethical Review Body and the United Kingdom Home Office (Autolus PPL No. P244BBE6B). All procedures are performed in accordance with the United Kingdom Home Office Animals (Scientific Procedures) Act 1986 and in adherence to Imperial College London or Autolus SOPs. This study is necessary and justifiable with due consideration to the ‘3Rs’ (the reduction, refinement and replacement of animals in research).

### Establishment of subcutaneous syngeneic colon carcinoma mouse model

6-8 weeks old Balb/c female mice (strain code 028, Charles Rivers) were subcutaneously injected with 1 x 10^6^ (100 μL) of CT26 colon carcinoma cells modified to express GD2. 9 days after tumour engraftment the mice were sub-lethally irritated with 4.5 Gy TBI. 24 hours after TBI, 1 x 10^6^ transduced CAR T cells were injected iv (100 μL). Tumour growth was monitored twice a week via calliper measurements. Peripheral blood via tail cut was taken every 7 days. Mice were euthanized when the tumor reached a maximum length or breadth of 1 cm^3^ or sudden body weight loss ≥ 20%. Animals were sacrificed by Isoflurane intoxication followed by cervical dislocation. Spleen were collected at the time of euthanization for CAR T cells tracking.

### Establishment of subcutaneous syngeneic melanoma mouse model

6-8 weeks old female C57Bl/6J mice (strain code 632, Charles River), were subcutaneously injected with 1 x 10^5^ (100 μL) of B16F10 melanoma cells modified to express GD2. 7 days after tumour engraftment the mice were sub-lethally irritated with 4.5 Gy TBI.

24 hours after TBI, 3 x 10^6^ transduced T cells were injected iv (100 μL). Tumour growth was monitored twice a week via calliper measurements. Peripheral blood via tail cut was taken every 7 days. Mice were euthanized when the tumor reached a maximum length or breadth of 1 cm^3^ or sudden body weight loss ≥ 20%. Animals were sacrificed by Isoflurane intoxication followed by cervical dislocation. Bone marrow and spleen were collected at the time of euthanization for CAR T cells tracking.

### Establishment of metastatic neuroblastoma xenograft mouse model

10–14 weeks old female NOD.Cg-Prkdc scid Il2rg tm1Wjl /SzJ (NSG, Jackson Lab, #005557) were intravenously injected with 1 x 10^6^ Firefly-luciferase expressing GD2^+^ CHLA-255 cells (CHLA-255 FLluc, in house) (100 μL). 15 days later, mice were injected intravenously with 1 x 10^6^ transduced CAR T-cells (100 μL). Tumor growth was indirectly assessed by bi-weekly bioluminescent imaging.

The mice were imaged for bioluminescence signal from tumor cells using the IVIS® system (IVIS, Xenogen Corporation, Alameda, CA) 10–15 minutes after 150 mg/kg D-luciferin (Xenogen) per mouse was injected intraperitoneally. Mice were euthanized when the sudden body weight loss ≥ 20%. Animals were sacrificed by Isoflurane intoxication followed by cervical dislocation. Bone marrow and spleen were collected at the time of euthanization for CAR T cells tracking.

### Establishment of human acute lymphoblastic leukemia (ALL) xenograft mouse model

10–14 weeks old female NOD.Cg-Prkdc scid Il2rg tm1Wjl /SzJ (NSG, Jackson Lab, #005557) were intravenously injected with 1 x 10^6^ Firefly-luciferase expressing NALM6 cells (NALM6 FLuc, in house) (100 μL). 4 days later, mice were injected intravenously with 2.5 x 10^6^ transduced CAR T-cells (100 μL). Tumor growth was indirectly assessed by bi-weekly bioluminescent imaging. The mice were imaged for bioluminescence signal from tumor cells using the IVIS® system (IVIS, Xenogen Corporation, Alameda, CA) 10–15 minutes after 150 mg/kg D-luciferin (Xenogen) per mouse was injected intraperitoneally. Mice were euthanized when body weight loss ≥ 20%. Animals were sacrificed by Isoflurane intoxication followed by cervical dislocation. Bone marrow and spleen were collected at the time of euthanization for CAR T cells tracking.

### Statistical analysis

Data were presented as mean ± SEM unless indicated otherwise. Graphs and statistics were generated using Prism 9.0 software for Windows (Graphpad Software Inc., La Jolla, CA, RRID:SCR_000306). The differences between means were calculated using two-tailed unpaired *t*-test, one-way ANOVA, and two-way ANOVA. Turkey’s correction for multiple comparisons was used to calculate adjusted p-value when appropriate.

Specific statistical test used for each figure was described in the corresponding figure legend. Survival determined from the time of tumor cell injection was analyzed by the Kaplan-Meier method and differences in survival between groups were compared by the log-rank test.

P values: ns P > 0.05, * P ≤ 0.05, ** P ≤ 0.01, *** P ≤ 0.001, ****, P ≤ 0.0001.

### Illustrations

All illustrations were created with BioRender.com (RRID:SCR_018361)

## Data Availability

The data generated in this study are available upon request from the corresponding author. The transcriptomic data generated in this study are publicly available in Gene Expression Omnibus (GEO) at GSE227161 and GSE227312.

## RESULTS

### The IL2 receptor endodomains can be constitutively activated by Ig Fab domain-induced heterodimerization

We sought to engineer a constitutively active IL2 receptor by approximation of the IL2 receptor β chain (IL2Rβ) and the common γ chain (CγC). To achieve this, the transmembrane and intracellular domains of IL2Rβ and CγC were fused with the human Ig constant heavy chain 1 (CH1) and light κ chain constant region (ΚLC) respectively (dFab_CCR-IL2). dFab_CCR-IL2 chains and the marker gene RQR8 were co-expressed in a single retroviral vector (Fig. 1A). Flow-cytometric analysis of dFab_CCR-IL2 transduced T cells via detection of Light Kappa chain (Fig 1B) or CH1 chain (Supplementary Fig S1A) showed that both dFab components were expressed, however mainly detected intracellularly. When cultured under conditions of cytokine starvation, dFab_CCR-IL2-expressing T cells increased in numbers (Fig. 1C, right), and diluted CTV dye (Fig. 1C, left), suggesting that some cytokine receptor signaling was recapitulated.

**Figure 1.**
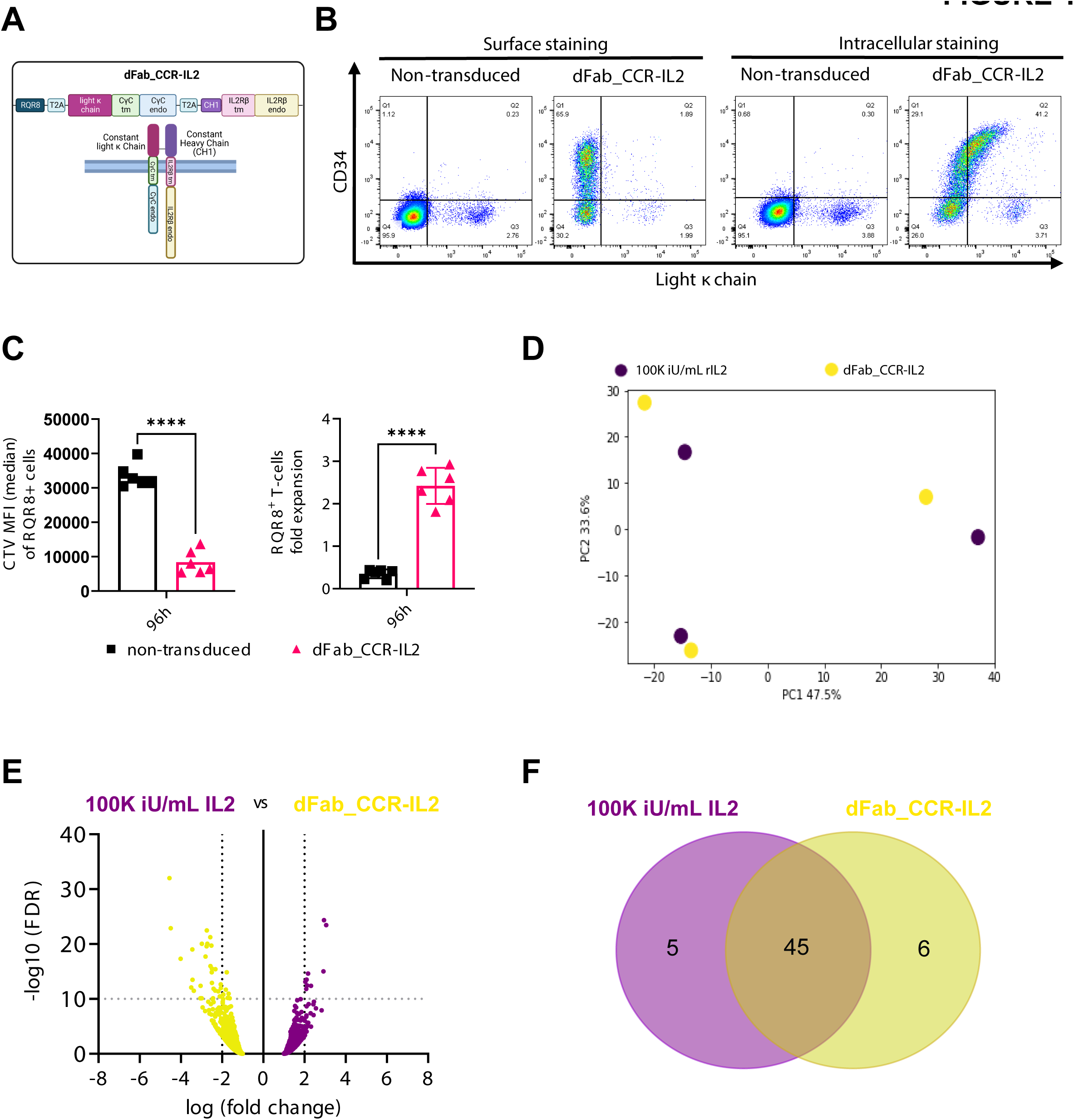
Constitutive heterodimeric dFab_CCR-IL2 recapitulate a proliferative IL2 signal in primary T cells. **(A)** Schematic of the dFab_CCR-IL2 format composed of the Constant heavy chain fused to the IL2 receptor β chain. The IL2 receptor CγC was fused to the Constant light Kappa chain (dFabκ_CCR-IL2). **(B)** Flow-cytometric analysis of the constant Light Kappa chain expression and RQR8 (CD34) in dFab_CCR-IL2 expressing T cells evaluated by surface staining or intracellular staining after fixation/permeabilization. **(C)** Quantitated proliferation of dFab_CCR-IL2 T cells cultured for 96 hours in cytokine starvation. Proliferation expressed as CTV MFI dilution (left), and fold expansion (right) of RQR8^+^ / CD3^+^ T cells (n=6, student t-test). **(D)** PCA plot analysing the variation in gene expression profile driven by the donors and by the two conditions. **(E)** Volcano plot representing the differential gene expression between dFab_CCR-IL2 T cells and non-transduced T cell stimulated with 100000 iU/mL IL2 (n=3). **(F)** Venn Diagram of the top 50 GSEA (Gene Set Enrichment Analysis) between dFab_CCR-IL2 T cells and T cell stimulated with 100000 iU/mL IL2 (n=3).

We next determined that dFab_CCR-IL2 signal can be modulated by truncating the receptor endodomain (20,21) from either the CγC (Supplementary Fig. S2A, B) or IL2Rβ (Supplementary Fig. S2C, D). Alternatively, dFab_CCR-IL2 signal can be controlled by off-the-shelf FDA-approved JAK inhibitor ruxolitinib (22) (Supplementary Fig. S3A).

These data suggested that Fab-based cytokine receptor is a versatile way of achieving constitutive cytokine receptor function.

### dFab_CCR-IL2 induces functional effects similar to those of the native IL2 receptor

To determine whether differences exist in signaling between the dFab_CCR and the native cytokine receptor, we compared T cells transduced with the dFab_CCR-IL2 with non-transduced T cells cultured in the presence of 100000 iU /mL exogenous IL2. Both resulted in T cell proliferation whilst T cells transduced with the dFab_CCR-IL2 expanded more than non-transduced T cells in 100000 iU/ml IL2 (Supplementary Fig. S4A), and both exogenous IL2 stimulation and dFab-CCR_IL2 expression resulted in phosphorylated STAT5 (Supplementary Fig. S4B).

We next performed a transcriptomic analysis following 7 days of cytokine starvation of dFab_CCR-IL2 expression compared with exogenous cytokine (Supplementary Fig. S4C). Principal component analysis (PCA) revealed no influence from the donors tested (n=3), whilst showing a degree of separation between the two conditions (Fig. 1D, Supplementary Fig. S4D). Nonetheless, differential gene expression analysis showed no global transcriptomic shift between the signaling delivered by the dFab_CCR-IL2 and exogenous IL2 receptor (Fig. 1E).

Given the similarity in the overall gene expression, we interrogated differences in specific pathways activated by the dFab_CCR-IL2 and the exogenous IL2 stimulation. Forty-four of the top 50 up regulated pathways (Supplementary Table S1) were shared between the two conditions (Fig. 1F). Furthermore, the FDR q-value of the top 10 shared pathways revealed a more pronounced proliferative profile from the dFab_CCR-IL2 transduced T cells with lower overall values than for the non-transduced T cells stimulated with exogenous IL2 (Supplementary Table S1).

Taken together, these data indicated that dFab_CCR-IL2 provides a functional cytokine-like signal.

### The dFab_CCR format can be used to activate multiple cytokine receptor families

A broad range of cytokine dFab-CCR formats were generated (Fig. 2A,) and grouped by respective cytokine receptor family: CγC receptor, the IL12 family, the IL1 family, IL10 and IL17 family (Supplementary Fig. S5A). Surface and intracellular Fab expression (Light Kappa %) and Fab density levels (Light Kappa MFI) detected by flow-cytometry varied between receptor families (Supplementary Fig. S5C, D), but were comparable within the same receptor family. Functional effects in primary human T cells were then explored.

**Figure 2.**
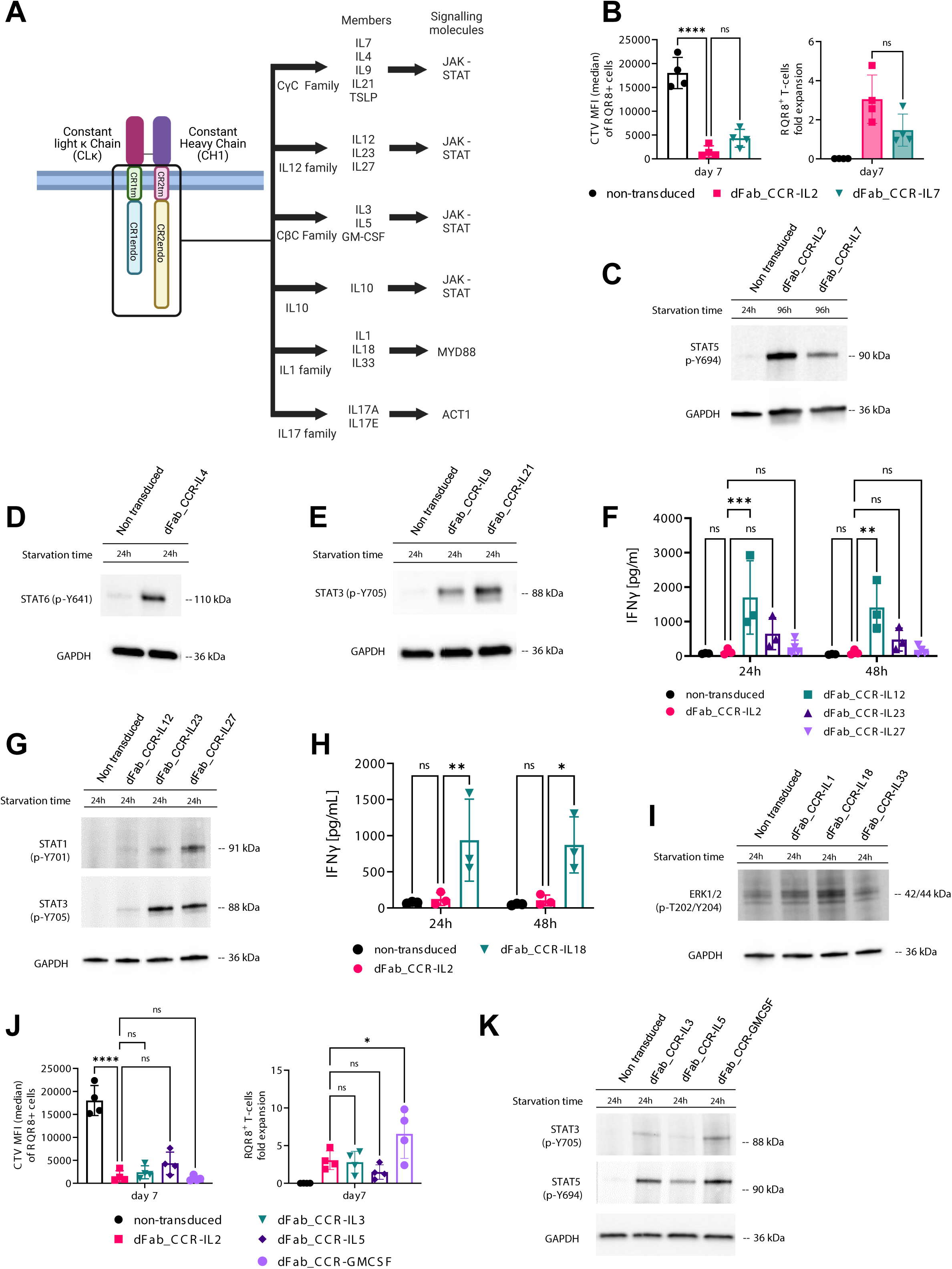
The dFab-CCR heterodimerisation format activates a broad range of cytokine receptors families. **(A)** Schematic of the main cytokine receptors family members and the viral vector design selected for the generation of the dFab_CCR library. **(B)** Quantitated proliferation of T cells engineered with either IL2 or IL7 dFab_CCRs, expressed as CTV MFI dilution (left), and fold expansion (right) of RQR8^+^ / CD3^+^ T cells (n=4, one-way ANOVA). **(C)** Immunoblot analysis of STAT5 phosphorylation, Y694. Non-transduced T cells were cultured in cytokine starvation for 24 hours. dFab_CCR-IL2 or -IL7 were cytokine starved for 96 hours. GAPDH was used as loading control. Data are representative of three independent experiments. **(D)** Immunoblot analysis of in vitro STAT6 phosphorylation, Y641. Non-transduced and dFab_CCR-IL4 T cells were cultured in cytokine starvation for 24 hours. GAPDH was used as loading control. Data are representative of three independent experiments. **(E)** Immunoblot analysis of in vitro STAT3 phosphorylation, Y705. Non-transduced, IL9 and -IL21 dFab_CCR T cells were cultured in cytokine starvation for 24 hours. GAPDH was used as loading control. Data are representative of three independent experiments. **(F)** IFNγ secreted by -IL2, -IL12, -IL23 and -IL27 dFab_CCR transduced T cells cultured for either 24 or 48 hours in cytokine starvation (n=3, one-way ANOVA). **(G)** Immunoblot analysis of in vitro STAT1 and STAT3 phosphorylation. Non-transduced, dFab_CCR-IL12, -IL23 and -IL27 T cells were cytokine starved for 24 hours. GAPDH was used as loading control. Data are representative of three independent experiments. **(H)** IFNγ secreted by IL2, -IL18dFab_CCR T cells coltured for either 24 or 48 hours in cytokine starvation (n=3, one-way ANOVA). **(I)** Immunoblot analysis of in vitro ERK1/2 phosphorylation, T202 and Y204. Non-transduced, IL1, -IL18 and -IL33 dFab_CCR T cells were cytokine starved for 24 hours. GAPDH was used as loading control. Data are representative of three independent experiments. **(J)** Quantitated *in vitro* proliferation of T cells engineered with either -IL3, -IL5 and -GMCSF dFab_CCRs, expressed as CTV MFI dilution (left), and fold expansion (right) of RQR8^+^ / CD3^+^ T cells (n=4, one-way ANOVA). **(K)** Immunoblot analysis of in vitro STAT3 and STAT5 phosphorylation. Non-transduced T cells were cytokine starved for 24 hours. IL3, IL5 and GMCSF dFab_CCRs T cells were cytokine starved for 96 hours. GAPDH was used as loading control. Data are representative of three independent experiments. All data are presented as mean ± SEM.

Western-blot analysis of the dFab_CCR CγC receptor family demonstrated the correct activation of the respective STAT molecules (Fig. 2C-2E), yet only the dFab_CCR-IL7 was able to induce T cell proliferation (Fig. 2B, Supplementary Fig. S6A-S6C).

The IL12 family members (IL12 and IL23) and IL18 from the IL1 receptor family dFab_CCRs showed increased baseline IFNγ release both at 24 and 48 hours (Fig. 2F, H), as expected (23,24). Furthermore, all the IL12 receptor family (Fig. 2G, STAT1 and STAT3), IL1 family (Fig. 2I, ERK1/2) and both IL10 andIL17A (Supplementary Fig. S6D, E) dFab_CCRs, correctly activated their respective signaling molecules.

Finally, we determined whether dFab_CCR could be applied beyond lymphoid cytokine receptors, by testing myeloid cells specific common β-chain: dFab_CCR-IL3, -IL5 and - GMCSF (Supplementary Fig. S5A). T cells engineered with all the CβC dFab_CCRs displayed significant proliferation (Fig. 2J) and resulted in the phosphorylation of either STAT3 or STAT5 proteins (Fig. 2K).

Taken together these results demonstrated that the dFab format could be applied to broad spectrum of cytokine receptor families.

### dFab_CCR-IL2 and CAR can function when co-expressed with a CAR in a single vector

We initially tested whether RQR8/dFab_CCR-IL2 could be co-expressed with a 2nd generation 41bbζ GD2 CAR (Fig. 3A). Expected RQR8 and dFab_CCR-IL2 ratio was maintained (Supplementary Fig. S7A) and co-expressing the dFab_CCR-IL2 provided a proliferative advantage (Supplementary Fig. S7B, S7C), whilst killing and cytokine secretion via the GD2 CAR was largely comparable (Supplementary Fig. S7D -S7F).

**Figure 3.**
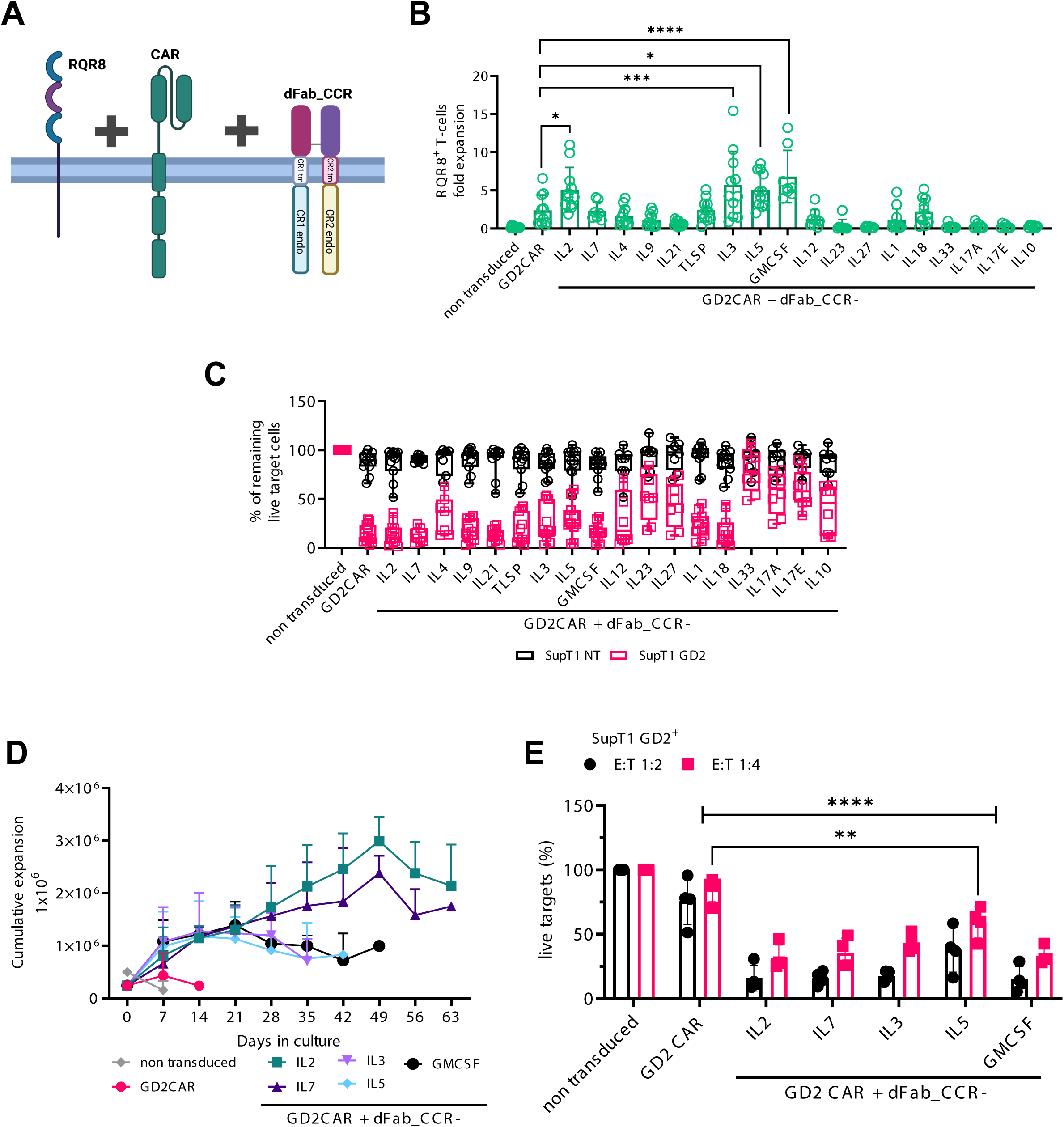
Co-expression of the dFab_CCRs with GD2 CAR reveal differential effects on CAR T cell functionality. **(A)** Schematic of the of the tetra-cistronic γ-RV vectors co-expressing RQR8, 2^nd^ generation 41bbζ GD2 CAR and dFab_CCR. **(B)** Day 7 quantification of *in vitro* proliferation of GD2 CAR T cells engineered with the library of dFab_CCRs, expressed as fold expansion of RQR8^+^ / CD3^+^ GD2 CAR T cells (n=11, GD2CAR + dFab_CCR-IL7 n=4one-way, ANOVA). **(C)** Killing of SupT1-NT (black) or SupT1-GD2^+^ after 48 hours co-culture with CAR-T cells co-expressing the library of dFab_CCRs at a 1:4 effector:target ratio. Data shows mean percentage (± SD) of live cells compared to non-transduced (NT) control (n=11, GD2CAR + dFab_CCR-IL7 n=4, two-way ANOVA). **(D)** Extended *in vitro* persistence of IL2, IL7, IL3, IL5 or GMCSF dFab_CCR co-expressing GD2 CAR T cells cultured in cytokine-free complete cell culture media. Live cells were counted weekly using DAPI/acridine orange (AO)I staining. Data shows mean 10^6^ live cells (± SD) (n=6). **(E)** Killing of and GD2^+^ after 48 hours co-culture with GD2 CAR-T cells recovered after 7 days of cytokine starvation at 1:2 and 1:4 effector:target ratio. Data shows mean percentage (± SD) of live cells compared to non-transduced (NT) control (n=4, two-way ANOVA). Data are representative of three independent experiments. All data are presented as mean ± SEM.

Given that these preliminary data indicated that dFab_CCR-IL2 may have a functional benefit when co-expressed with a CAR, we proceeded to test 18 dFab_CCRs receptors, representing different cytokine receptor families, co-expressed with the same GD2 CAR. Co-expression of dFab_CCR-IL2 or -GMCSF with the GD2 CAR resulted in the strongest antigen independent proliferation, with a fold expansion between 5 and 7 (Fig. 3B). Looking at the CAR effector function, the expression of most of the CγC dFab_CCRs (IL2, IL7), CβC (GMCSF) or the pro-inflammatory cytokine receptors (IL1, IL18) preserved CAR T cell cytotoxicity (Fig. 3C). The remaining dFab_CCRs library impacted GD2 CAR T cell cytotoxicity with different magnitudes (Fig. 3C).

Most of the dFab_CCRs tested did not alter CAR-mediated IFNγ secretion, however, IL18 or IL12 dFab_CCRs increased IFNγ secretion in response to cognate antigen (Supplementary Fig. S8A). Additionally, dFab_CCR-IL18 expression enhanced IL2 secretion (Supplementary Fig. S8B). Similar results with respect to proliferation, cytotoxicity and cytokine secretion were obtained with the dFab_CCR IL2, IL7, GMCSF and IL18 co-expressed with a second-generation CD19 CAR (Supplementary Fig. S9A, S9B). Additionally, dFab_CCR-IL18 demonstrated increased IFNγ and IL2 secretion when co-expressed with the second-generation CD19 CAR (Supplementary Fig. S9C, S9D). Finally, co-expression of dFab_CCRs with an irrelevant second-generation CAR did not mediate nonspecific tumor cell killing or autonomous cytokine secretion despite demonstrating the expected proliferation (Supplementary Fig. S10).

### dFab_CCRs prolonged CAR T cell expansion over time

Constitutive cytokine signaling could cause uncontrolled T cell proliferation. To address this, we challenged a selection of dFab_CCRs to continuous antigen and cytokine starvation. Non-transduced T cells died within the first week in culture, whereas CAR alone sustained T cells for only two weeks (Fig. 3D). All the dFab_CCRs CAR T cells expanded over the first 21 days (Fig. 3D). Interestingly, after day 21 the CβC dFab_CCRs and the CγC dFab_CCRs proliferative potency diverged. The CβC dFab_CCRs exhibited a gradual reduction in live cell number until day 49. Conversely, the CγC dFab_CCRs (IL2 and IL7) maintained a steady cell growth peaking at day 49, and a steady contraction until day 63 (Fig. 3D).

We then assessed whether cytokine withdrawal would affect CAR T cells cytotoxicity. Seven day-starved CAR T cells were co-cultured with GD2^+^ target cells (SupT1 GD2^+^). At both E:T ratios tested, the selected dFab_CCRs tested retained a strong CAR T cell cytolytic capacity (Fig. 3E). Concordantly, dFab_CCRs co-expression sustained elevated IFNγ secretion (Supplementary Fig. S8C). Expression of CAR alone failed to protect both cytotoxicity and IFNγ secretion.

These results demonstrated that the selection of dFab_CCRs showed a prolonged but finite proliferative potential.

### dFab_CCRs influence CAR T cells performance after serial antigen exposure

*In vitro* chronic antigen exposure is a relevant way to evaluate T cell functional exhaustion (25). We therefore subjected the dFab_CCRs CAR T cells to weekly exposure of fresh antigen-positive tumor cells (Fig. 4A). In contrast to dFab_CCR-IL12, all the dFab_CCRs CAR T cells expanded in cell number following 3 rounds of antigen stimulation. with the GMCSF dFab_CCR inducing the strongest expansion, to a greater degree compared to either GD2 CAR (Fig. 4B). Importantly, chronic stimulation did not interfere with dFab_CCR signaling. Co-expression of IL2, and GMCSF dFab_CCRs resulted in 3-to-4-fold CAR T cell expansion compared to CAR T alone, while IL7 dFab_CCRs conferred more modest proliferation (Fig. 4C). The remaining dFab_CCRs showed differential functionality, with the CβC CCRs mediating sustained expansion (Supplementary Fig. S11). Repeated stimulations reduced the cytolytic ability of GD2 CAR T cells (Fig. 4D). No improvement in cytolytic ability was observed with the co-expression of IL7, or IL18 dFab_CCR. In contrast, IL2 and IL12 dFab_CCR showed a trend of improved T cell cytotoxicity (Fig. 4D), with dFab_CCR-GMCSF showing significant enhancement. Co-expression of the dFab_CCRs with GD2 CAR resulted in increased levels of IFNγ (Supplementary Fig. S12), with IL2, IL7, GMCSF and IL18 dFab_CCRs doubling the amount of IFNγ secreted, whilst dFab_CCR-IL12 improved IFNγ secretion to approximately 4-fold compared to GD2 CAR alone.

**Figure 4.**
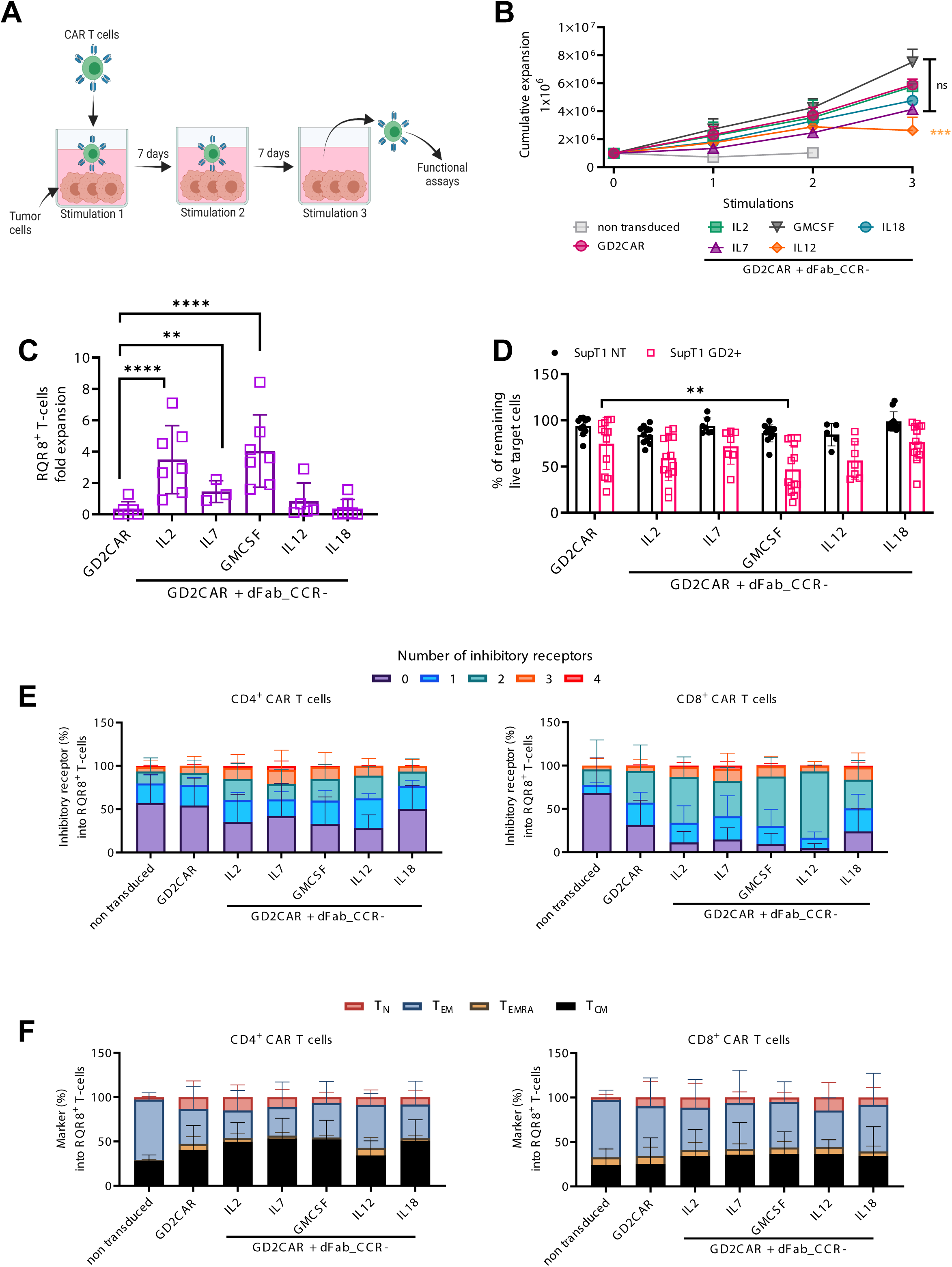
dFab_CCR co-expression sustain GD2 CAR T cell function after chronic antigen exposure. **(A)** Chronic antigen stimulation experimental design. The first co-culture was initiated with 1×10^6^ GD2 CAR T-cells together with 0.1×10^6^ SKOV3 cells for 7 days, in the absence of exogenous cytokines. For the second and third co-cultures, T-cells were harvested from the previous co-culture and then replated in new culture medium with fresh tumour cells at the same 10:1 E:T ratio. **(B)** Cumulative expansion of selected GD2 CAR T cells from (A) after three rounds of co-culture (n=11, n=4 GD2CAR + dFab_CCR-IL7, one-way ANOVA). **(C)** Quantitated *in vitro* proliferation of GD2 CAR T cells recovered after three rounds of co-culture (B) and cultured for 7 days in cytokine starvation condition. Proliferation expressed as fold expansion of RQR8^+^ / CD3^+^ GD2 CAR T cells (n=11, GD2CAR + dFab_CCR-IL7 n=4, one-way ANOVA). All data are presented as mean ± SEM. **(D)** Killing of SupT1 NT (black) and GD2^+^ (red) after 48 hours co-culture with GD2 CAR-T cells recovered after three rounds of co-culture (B). at 1:4 effector:target ratio. Data shows mean percentage (± SD) of live cells compared to non-transduced (NT) control (n=11, GD2CAR + dFab_CCR-IL7 n=4, GD2CAR + dFab_CCR-IL7, two-way ANOVA). **(E)** Exhaustion phenotype of CAR T cells after three rounds of chronic antigen exposure. Exhaustion evaluated by the expression of TIM3, LAG3, PD-1 or KLRG1 in either CD4^+^ (top) or CD8^+^ (bottom) T cells. Stacked bars show percentage of cells expressing 0, 1, 2, 3 or 4 markers per individual donors (n=11, GD2CAR + dFab_CCR-IL7 n=4, one-way ANOVA). **(F)** Memory phenotype of CAR T cells after three rounds of chronic antigen exposure. Memory phenotype evaluated by the expression of CD45RA and/or CCR7 in either CD4^+^ (top) or CD8^+^ (bottom) T cells. Memory phenotype defined as: naïve T cells (CD45RA^+^, CCR7^+^), central memory (CM) T cells (CD45RA^-^, CCR7^+^), effector memory (EM) T cells (CD45RA^-^, CCR7) and terminally differentiated (TEMRA) T cells (CD45RA^+^, CCR7^-^). Stacked bars show percentage of cells in each population markers per individual donors (n=11, GD2CAR + dFab_CCR-IL7 n=4, one-way ANOVA). All data are presented as mean ± SEM.

When co-expressed with a CD19 CAR, repeated stimulation resulted in similar expansion of T cells expressing both the CAR and the selected dFab_CCRs (Supplementary Fig. S13A). However, chronic stimulation did not impair CD19 CAR T cell cytotoxicity, regardless of presence or absence of the dFab_CCRs (Supplementary Fig. S13B). Here, the dFab_CCR-18 was able improve IFNγ secretion 3-fold compared to CD19 CAR alone (Supplementary Fig. S13C).

### dFab_CCR co-expression maintain a less differentiated and a non-exhausted CAR T cell phenotype

We next sought to determine the effect of each of the dFab_CCRs on CAR T cell memory differentiation and exhaustion (Fig 4E, F, Supplementary Fig. S14A, B). After three rounds of antigen exposure IL2, IL7, the GMCSF and the IL12 dFab_CCRs showed increased percentage of cells expressing multiple exhaustion markers compared to the CAR alone (Fig. 4E), with a more prominent effect in the CD8^+^ T cell population.

The IL18 dFab_CCR, despite enhancing IFNγ and IL2 secretion, maintained a similar exhaustion marker expression profile as the CAR alone (Fig. 4E). The majority of the dFab_CCRs expanded the central memory population in both the CD4^+^ and CD8^+^ T cell compartment (Supplementary Fig. S14B). Importantly, IL2, IL7, GMCSF, IL18 dFab_CCR demonstrated a less differentiated phenotype (Fig. 4F). Others, (IL9, IL21 and the IL12) showed an increased percentage of terminally differentiated effector cells and effector memory population, with a more pronounced effect in CD8^+^ compartment (Supplementary Fig. S14B). Similarly, co-expression of the selected dFab_CCRs with the CD19 CAR promoted lower exhaustion marker expression and mediated a less differentiated phenotype despite the enhanced functionality (Supplementary Fig. S15A, B).

### dFab_CCR-IL18 improved human CAR T cell persistence in neuroblastoma xenograft model

Only a small panel of dFab CCRs can be easily screened in parallel *in vivo* due to the low-throughput characteristic of standard models. In order to assess the persistence of the dFab_CCRs *in vivo* we employed a pooled DNA library of barcoded GD2 CAR dFab_CCRs constructs which allows assessment of the proliferation of each individual CCR (Supplementary Fig. S16). We inoculated T cells expressing the library into NSG mice engrafted with GD2-positive CHLA255 tumors (Fig. 5A, B). Flow cytometric analysis showed higher CAR T cell engraftment in CHLA-bearing mice (Fig. 5 C and 5E), compared to tumor free mice (Figure 5D, 5F). The analysis of barcodes frequency from both tumor-free and tumor-bearing NSG mice showed expansion of IL2, IL4 and IL18 dFab_CCRs compared to their relative abundance at the day of injection (Fig. 5E, F). The greatest degree of expansion was shown by the dFab_CCR-IL18 representing 40% the total CAR T cell population recovered (Fig. 5F) *in vivo*.

**Figure 5.**
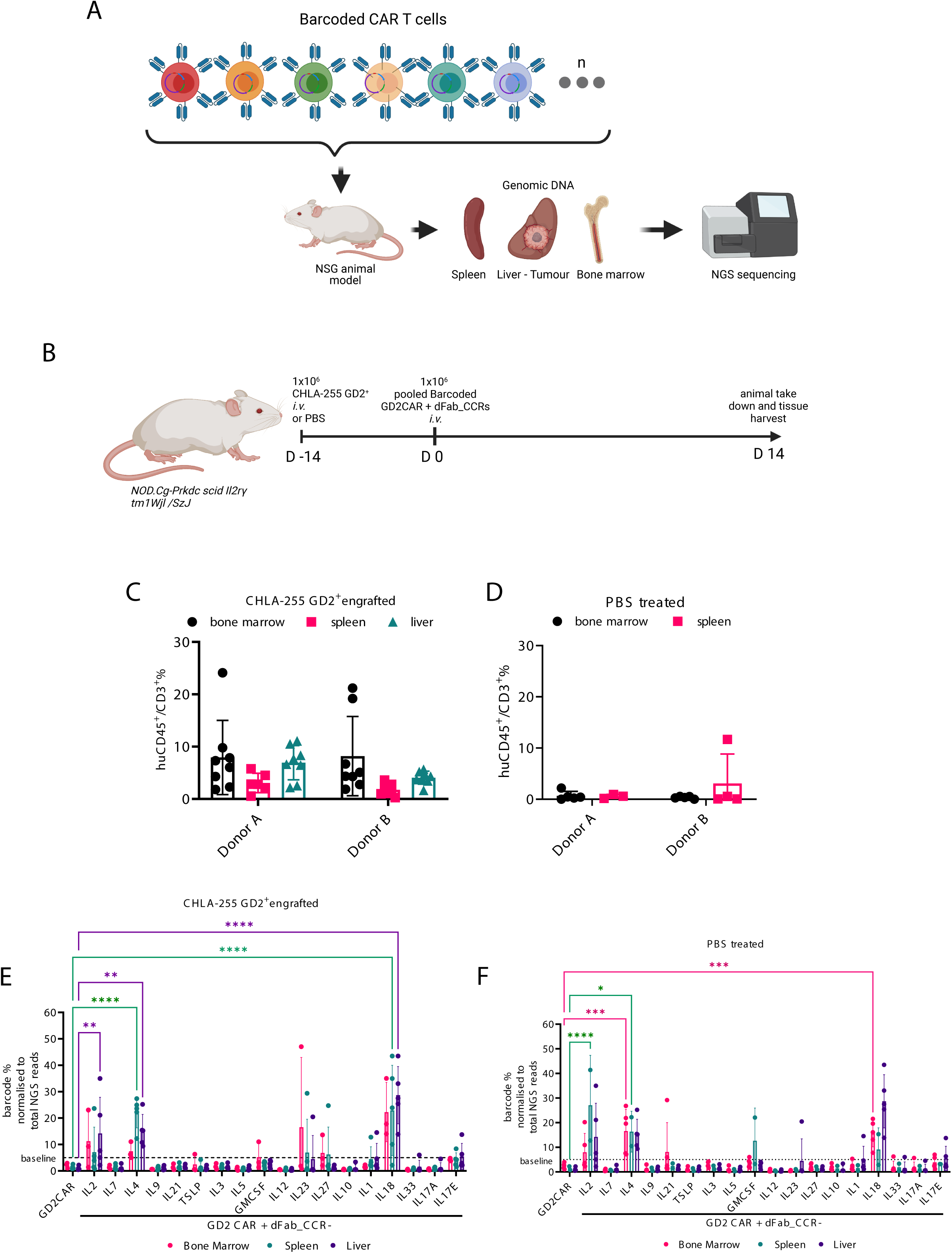
The pooled DNA-barcoded dFab_CCRs library identifies pro-survival dFab_CCRs *in vivo*. **(A)** Experimental set-up the xenograph *in vivo* model of the barcoded GD2CAR co-expressing dFab_CCRs pooled library. **(B)** NOD.Cg-Prkdc scid Il2rg tm1Wjl /SzJ (NSG) mice were either engrafted with the GD2-expressing CHLA neuroblastoma cell line or left tumour free. **(C, D)** Flow cytometry analysis of barcoded human CAR T cells pooled population, 14 days after injection. Graphs represent the percentage of CD45+ / CD3+ human T cells from CHLA255 neuroblastoma xenograft (C) or PBS control (D) NSG mice. All data are presented as mean ± SEM. **(E, F)** Quantitated representation of individual barcoded frequency within the pooled barcode population 14 days post infusion in CHLA255 neuroblastoma xenograft (E) or PBS control (F) NSG mice (n=5 mice per group, one-way ANOVA). All data are presented as mean ± SEM.

### dFab_CCR-IL18 and GMCSF co-expression promoted a distinctive T cell effector profile

To understand better the unexpected *in vivo* effects of either the dFab_CCR-IL18 or dFab_CCR-GMCSF, transcriptomic, cytokine array and metabolomic analysis were performed on a selection of dFab_CCRs in CAR T cells. First, the NanoString was utilized in resting conditions or following GD2 CAR activation (Supplementary Fig. S17A). Principal component analysis (PCA) indicated that the dFab_CCR-IL18 promoted a distinctive transcriptomic profile following activation (Fig. 6A), clustering distinctly from CAR alone or the other dFab_CCRs tested (IL2, IL7, GMCSF). IL2 and IL7 dFab_CCRs co-localized in both activated and non-activated CAR T cells (Fig. 6A). Finally, dFab_CCR-GMCSF segregated more closely with the IL7 and IL2 dFab_CCR, although retained a degree of separation (Fig. 6A).

**Figure 6.**
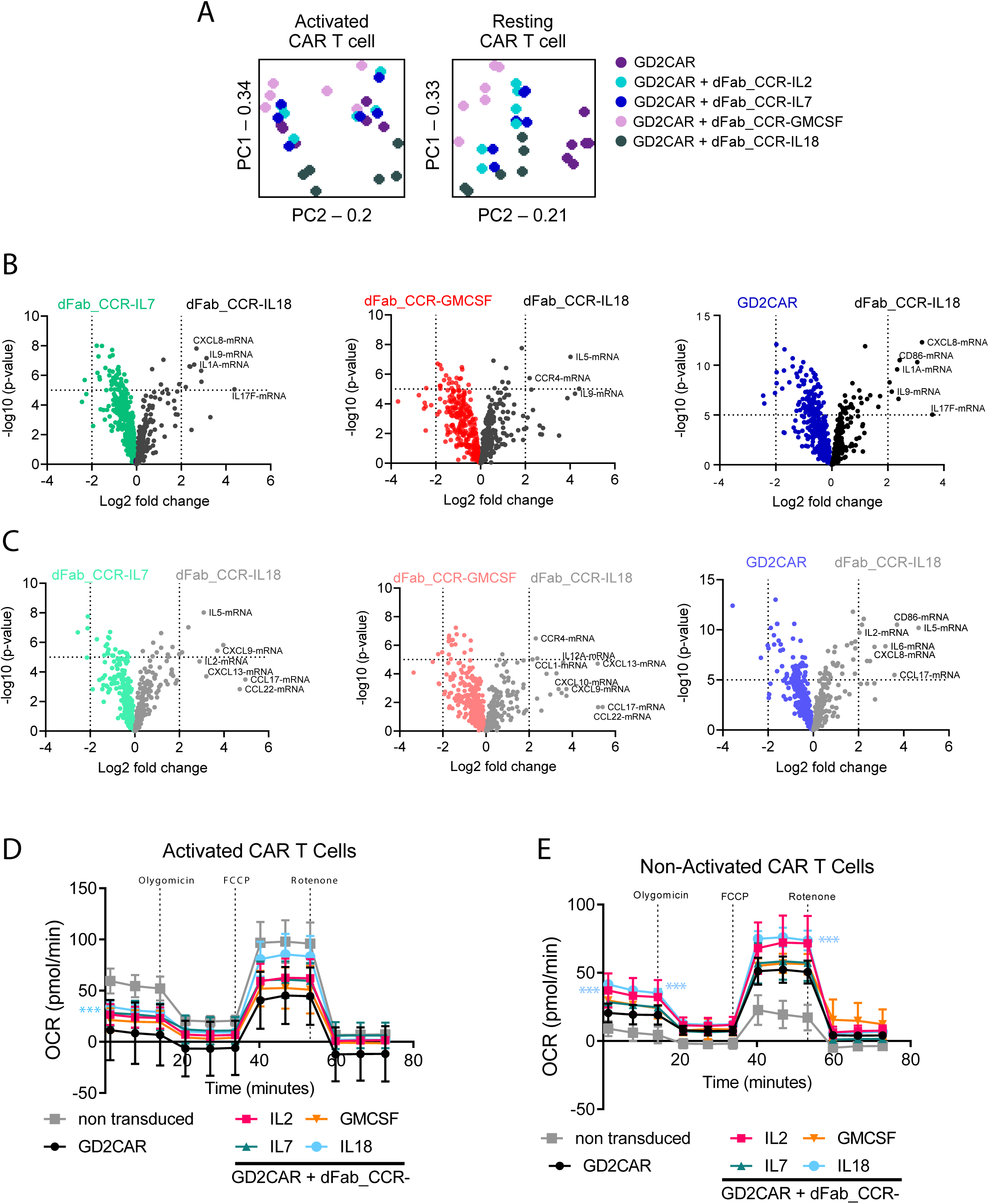
IL18 and GMCSF dFab_CCRs deliver functionally and transcriptomically different out-put to GD2 CAR T cell. **(A)** PCA analysis of each individual dFab_CCR transcriptome in activated CAR T cells (left) or non-activated (resting) CAR T cells (right), **(B)** Volcano plot representing the comparison of dFab_CCR-IL18 CAR T cells against either dFab_CCR-IL7, dFab_CCR-GMCSF or CAR alone in activated CAR T cells. **(C)** Volcano plot representing the comparison of dFab_CCR-IL18 CAR T cells against either dFab_CCR-IL7 or dFab_CCR-GMCSF or CAR alone in non-activated CAR T cells. **(D, E)** The oxygen consumption rates (OCRs) and the individual metabolic parameter were analysed either after 24 hours of antigen starvation (Non activated CAR T cells, D) or after 24 hours of activation (Activated CAR T cells, F) (*n* = 5, one-way ANOVA) All data are presented as mean ± SEM.

Differential gene expression analysis showed that dFab_CCR-IL18 co-expression uniquely influenced the expression of interleukin- and chemokine-related genes such as CXCL9 and CXCL10, cytokines such as the Th2 cytokine IL5 as well as IL17 and IL1 genes suggesting a more profound pro-inflammatory state (Fig. 6B: activated T cells, 6C: resting T cells). In accordance with the PCA data, differential gene expression confirmed the overlapping similarities of IL7 and IL2 dFab_CCRs (Supplementary Fig. S17B activated T cells, S17C resting T cells), whilst the dFab_CCR-GMCSF expression induced more pronounced differences in genes expression (Supplementary Fig. S17B, S17C).

We then tested the secretion of Th1, Th2 cytokines and chemokines. When co-expressed with either the GD2 or CD19 CAR, dFab_CCR-IL18 CAR T cells demonstrated a higher trend for chemokine secretion compared to CAR alone, despite not reaching statistical significance (Supplementary Fig. S18). Concurring with findings from Lange et al. (26), dFab_CCR IL18 expressing CAR T cells demonstrated increased levels of Th2 cytokines IL5 and IL13, particularly when co-expressed with a CD19 CAR (Supplementary Fig S18A, B). Interestingly, dFab_CCR-GMCSF co-expressing CAR T cells exhibited a reduction in the secretion of both chemokines and cytokines (Supplementary Fig. S18).

Finally, regardless of the CAR, dFab_CCR-IL18 influenced CAR T cell metabolic profile with higher respiratory capacity mainly in resting conditions (Fig. 6D: resting T cells, 6E: activated T cells). IL18 and IL2 dFab_CCRs drove a greater maximal respiratory capacity, and ATP production in non-activated T cells (Supplementary Fig. S19).

Furthermore, dFab_CCR-IL18 induced higher basal respiration in both resting and activated condition (Supplementary Fig. S19). Despite not reaching statistical significance dFab_CCR-IL18 showed higher maximal respiratory capacity and ATP production compared to the other dFab_CCRs in activated CAR T cells (Supplementary Fig. S19).

Overall, these data indicated that dFab_CCR-IL18 creates a distinct profile characterized by pro-inflammatory transcriptomic and metabolic state representative of a less differentiated T cell population (27).

### GMCSFmu and IL18mu dFab_CCR co-expression enhances murine GD2 CAR T cells antitumor activity in several tumour models

In order to test the efficacy of the selected dFab_CCRs on human CAR T cell product *in vivo*, we tested dFab_CCR/CD19 CAR T cells in a xenograft model of B-ALL (Supplementary Fig. S20A). dFab_CCR-GMCSF and -IL7 induced peripheral expansion (supp fig S20B) and complete tumor control by day 14 (Supplementary Fig. S20C, S21A). However, early onset of xeno GVHD was observed, hampering the longevity of the experiment. In contrast, dFab_CCR-18 promoted comparable anti-tumor activity, with complete tumor control until the end of the model (day 24), with no xeno GVHD observed. Importantly, none of the dFab_CCRs co-expressed with the irrelevant CAR promoted anti-tumor activity or GVHD toxicity (Supplementary Fig. S20C, S21B).

We then designed murine equivalents of the CCRs and confirmed function in murine T cells (Supplementary Fig. S22) prior to assessing function in a syngeneic, immunocompetent CT26 colon carcinoma *in vivo* model (Supplementary Fig. S23A). *In vitro,* CAR T cells alone were unable to mediate tumor rejection (Fig. 7A, Supplementary Fig S23B), resulting in no mice surviving past day 24 (Fig. 7B). In contrast, co-expression of the dFab_CCR_IL18mu completely controlled the tumor growth in 3 out of 10 mice, up to day 50, and delayed tumor progression in a further 3 mice (Fig. 7A, Supplementary Fig. S23B), with 50% of the mice alive at day 50 post CAR T cell injection (Fig. 7B). Similarly, dFab_CCR-GMCSFmu promoted a delay in tumor growth in 4 out of 8 mice, unpalpable tumor in 1 out of 8 mice by day 31 (Fig. 7A, Supplementary Fig. S23B), and improved mice survival by more than 15 days compared to the CAR alone (Fig. 7B). The enhanced tumor control displayed by dFab_CCR-IL18mu and GMCSFmu was accompanied by increased CAR T engraftment, with a higher peak at day 10 post-CAR T cell infusion.

**Figure 7.**
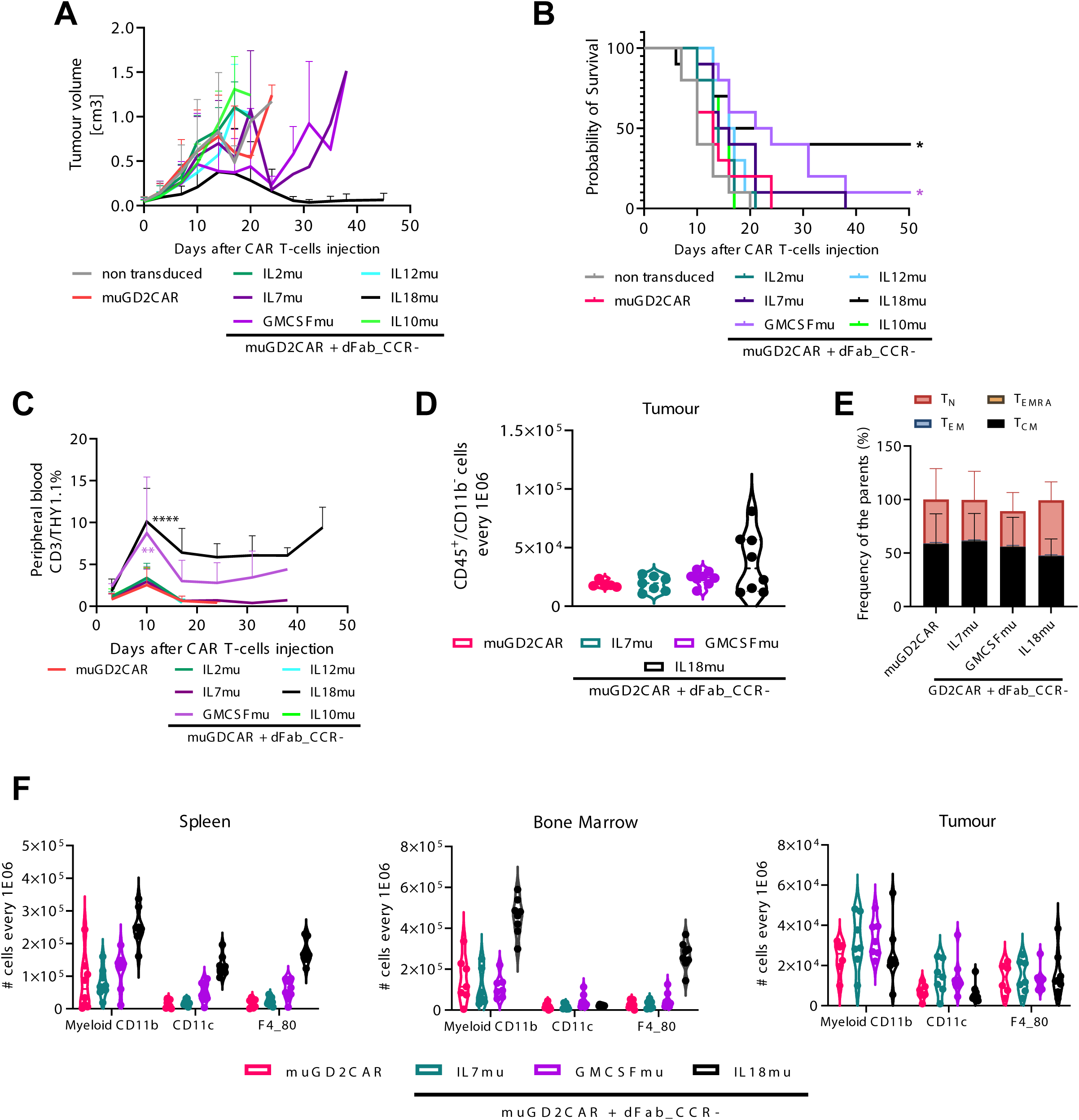
IL18 and GMCSF dFab_CCRs enhances adoptive CAR T cell immunotherapy against colon carcinoma immunocompetent animal models. **(A)** Tumour volume quantification over time (*n* = 8). **(B)** Kaplan Mayer survival curve. **(C)** GD2 CAR T cells peripheral engraftment quantitated via flow cytometry by the expression of murine CD45, murine CD3 and THY1.1 (n=8). **(D)** Day10 GD2 CAR T cells tumour trafficking measured as number of THY1.1^+^ CAR T cells every 1E06 CD3^+^/CD45^+^/CD11b^-^ lymphoid cells (n-8). **(E)** Memory phenotype of murine THY1.1 CAR T cells in the tumour. Memory phenotype evaluated by the expression of CD62L and CD44. Naïve T cells (C44^-^, CD62L^+^), effector memory (EM) T cells (CD44^+^, CD62L+), Central Memory (CM) T cells (CD44+, CD62L^+^) and terminally differentiated (TEMRA) T cells (CD44^-^, CD62L^-^). Stacked bars show percentage of cells in each population markers per individual donor (n=8). **(F)** Myeloid cells quantification in spleen, bone marrow, tumour defines by the expression of total CD11b (myeloid), CD11b/CD11c (dendritic cells), CD11b/F4-80 (Macrophages), every 1E06 CD45+/CD11b-cells. (n=8). All data are presented as mean ± SEM.

Furthermore, both dFab_CCRs sustained prolonged peripheral persistence of the transduced CAR T cells over time (Fig. 7C, Supplementary Fig. S23C). Notably, dFab_CCR-IL18mu was accompanied by weight loss in treated mice, resulting in the culling of 2 out of 9 mice by day 10 post-treatment; however, the remaining mice recovered weight to baseline (Supplementary Fig. S23D).

Enhanced functionality of CAR T cells expressing the dFab_CCRs was also observed in a highly aggressive B16F10 melanoma sub cutaneous model (Supplementary Fig. S24A). The CAR alone failed to control tumor growth with a comparable overall survival to non-transduced T cells (Supplementary Fig. S24B). Conversely, -IL7mu, -IL18mu and -GMCSFmu dFab_CCRs improved anti-tumor efficacy with prolonged survival up to day 31, 35 and 39 respectively (Supplementary Fig. S24B) without apparent toxicity manifesting as weight loss (Supplementary Fig. S24C). Additionally, dFab_CCR-GMCSFmu-treated mice showed significantly greater T cell engraftment (Supplementary Fig. S24D).

### dFab_CCR-IL18 influence the recruitments of immune cell in the tumour microenvironment

To investigate the improved anti-tumor efficacy in the syngeneic colon carcinoma and melanoma models, we evaluated the effect of dFab_CCR expression on trafficking, phenotype of the murine CAR T cells and the interaction with the host immune milieu (Supplementary Fig. S25A). dFab_CCR-IL18 CAR T cells exhibited improved trafficking within the tumor microenvironment (Fig. 7D), maintaining a less differentiated phenotype with a higher proportion of naïve and central memory T cell (Fig. 7E), and a higher proportion of central memory T cells in both bone marrow and spleen (Supplementary Fig. S25D). Additionally, dFab_CCR-18 expression induced prominent recruitment of myeloid cells, especially dendritic cells and macrophage in both bone marrow and spleen (Fig. 7F); whilst within the tumor, IL7 and GMCSF dFab_CCRs promoted higher myeloid cells recruitment (Supplementary Fig. S25C).

Finally, dFab_CCR-18 CAR T cells induced significantly higher IFNγ levels, and displayed a trend to increased levels of CXCL10 chemokine, TNFα and IL6, albeit without reaching statistical significance (Supplementary Fig. S25E).

Altogether, these data indicated that IL18 dFab_CCR promoted the strongest interaction with the host immune system; likely caused by its complex chemokine and cytokine secretion.

## DISCUSSION

In this study, we describe a new format of constitutively active cytokine receptor (dFab_CCR), which is broadly generalizable across different cytokine receptors families (figure 2). We tested the utility of co-expressing the dFab_CCRs in CAR T cells and demonstrated the enhancement of CAR T cell activity. Of note, we found that IL18 and GMCSF dFab_CCR conferred particular and differing functional benefit to CAR T cells.

Given that cytokine receptors usually require endodomain heterodimerization to signal, we explored using the Ig CH1/Light chain constant region interaction to induce cytokine receptor heterodimerization. CH1 and kappa or lambda light chain constant (KLC or LLC) form a densely packed and stable heterodimeric interface (28,29) and have been previously used for heterodimeric assembly of recombinant proteins (30,31). We hypothesized that we could induce constitutive cytokine receptor signaling by replacing cytokine receptor ectodomains with CH1 and the KLC region. Initial exploration of this strategy utilizing the IL2 receptor endodomains in primary human T cells demonstrated that an IL2 signal could be recapitulated by functional and transcriptional analysis.

Adoptive immunotherapy has frequently been combined with systemic administration of cytokines to enhance T cell activity; however, this often causes systemic toxicity (6). Strategies have been developed to engineer therapeutic immune cells to avoid systemic administration of cytokines. The simplest strategy is to engineer T cells to constitutively secrete a particular cytokine (7,8). This approach still risks systemic cytokine toxicity (Zhang et al., 2015) and is susceptible to down-regulation of the cognate cytokine receptor (32). Further strategies were devised which overcame these limitations. Early solutions included chimeric cytokine receptors which could be activated by pharmaceuticals such erythropoietin (33) or small molecules such as rapamycin (9). Alternative approaches were also developed to engineer constitutively active receptors, such as IL7 by cysteine and/or proline insertions in the IL7Rα transmembrane (CR7) (34). Despite being efficacious, this strategy is not applicable to cytokine receptors that function as homodimers, such as IL2 β-chain (20,35) or IL12Rβ2 homodimers (36).

Alternative methods include a membrane-tethered IL15 receptor generated by fusing the IL15 to the IL15Rα by a flexible linker (12). However, this approach might result in unwanted activation of neighboring cells. Furthermore, such formats could be restricted by γ-chain availability due to their interaction with other native common γ-chain receptor partners (37).

We add to this literature with the dFab_CCR approach. Following initial proof-of-concept with IL2, truncations of either the IL2Rβ chain, or common γ-chain endodomains was demonstrated to potentially enable “tuning” of the signal intensity and the degree of T cell proliferation.

Given the similar architecture of the entire class I cytokine receptor family, the versatility of the dFab_CCR was explored. Six cytokine receptor families comprising of a total of 18 different individual dFab_CCRs were investigated spanning from T cell to myeloid cell specific receptors. The dFab_CCR approach resulted in activation of downstream signaling molecules from all the 18-cytokine receptors tested. The IL12 family (IL12, IL23) and IL18 dFab_CCRs were functionally active resulting in high basal level of IFNγ secretion, possibly due to their higher expression density. Interestingly, despite being myeloid cell-specific, common β-chain dFab_CCRs (e.g., GMCSF) promoted strong proliferation in T cell through the activation of both STAT3 and STAT5.

Having established the versatility of dFab_CCR, the utility in the setting of adoptive immunotherapy was then investigated. Given the success of CAR T cells in lymphoid malignancies (38–40) and early promise in solid cancer (14), dFab_CCRs were tested in combination with both a GD2 and a CD19 second generation 41bbζ CARs. We explored whether the 18 individual dFab_CCRs conferred a differential functional advantage to CAR T cells. Using individualized *in vitro* dFab_CCR testing as well as a pooled barcoded *in vitro* and *in vivo* approach, the majority of dFab_CCRs maintained CAR T cell antigen-specific cytotoxicity and cytokine secretion. This remained true when a selection of dFab_CCRs was co-expressed with an irrelevant CAR.

Under conditions of chronic antigen stimulation, IL2 and the common β-chain dFab_CCRs sustained CAR T cell cytolytic activity upon serial re-challenge with fresh target cells, compared to the CAR alone. Importantly, IL2, IL7 and the common β-chain dFab_CCRs maintained enhanced CAR T cell expansion in the absence of exogenous cytokines, before and after repeated antigen stimulation, regardless of the co-expressed CAR partner. Additionally, these dFab_CCR-expressing CAR T cells displayed a higher proportion of the Tcm population suggesting a less differentiated state.

Surprisingly, when co-expressed with the GD2 CAR, dFab_CCR-CAR based upon immunostimulatory cytokines such as IL23 and IL27 (24) reduced cytolytic activity, cytokine secretion and hampered proliferation upon chronic antigen stimulation.

Notably, these experimental approaches highlighted distinct effects of dFab_CCR-IL18 on CAR T function. IL-18 is a pro-inflammatory cytokine belonging to the IL-1 superfamily. In T cells, IL18 facilitates CD4 T helper 1 (Th1) polarization in conjunction with IL12 while Th2 polarization together with IL-2 (41). In preclinical models, intra-tumoral production of IL18 showed preferable safety profile compared to IL12 (42–44). The IL18 receptor signals via MYD88 (45), which plays a critical role in innate immunity (46). MYD88 have used in adoptive cell therapy as an alternative CAR co-stimulatory domain, showing improved CAR T-cell effector function in the setting of chronic antigen stimulation (47–49). Furthermore, several approaches have been characterized by delivering an engineered IL18 cytokine or an IL18 receptor signal within the tumor microenvironment, showing increased chemokine and Th1 cytokine secretion (26,50,51).

In this study, dFab_CCR-IL18 increased CAR T cell persistence and anti-tumor response in two syngeneic animal models, enhanced survival in an NSG neuroblastoma model and enhanced anti-tumor control in a CD19 NSG model. Furthermore, in the syngeneic colon carcinoma model, a less differentiated phenotype of dFab_CCR-IL18 expressing CAR T cells was observed accompanied by an increased recruitment of the host lymphoid and myeloid cells.

Further transcriptomic and metabolomic *in vitro* analysis revealed that dFab_CCR-IL18 promoted a pronounced pro-inflammatory state, and a higher aerobic metabolism characteristic of less differentiated CAR T cells (27). Recently, Gottschalk and colleagues presented an elegant receptor switch that re-directs GMCSF secreted upon CAR T-cell activation, to an IL18 signal in CAR T cells (26). Concordantly, they also reported an increase Th2 cytokine secretion together with improved Th1 effector function and better *in vivo* tumor control (26).

This toolkit of a range of constitutively active CCRs displayed various ways could enhance CAR T cell therapy by: i) improvement of proliferation, ii) maintenance of effector function, iii) enhancement of cytokine secretion influencing the host immune system, iv) improved aerobic metabolism and v) acceptable safety profile. Gene therapies based upon the expression of the common γ-chain have been associated with malignant transformation in the context of HSCs SCID therapy (52).

However, such toxicities have been related to viral construct design (53,54); as subsequent clinical studies using self-inactivating lentivirus reported no malignant transformation (55). Here we demonstrate that the use of common γ-chain dFab_CCRs in mature T cells showed enhanced but time-limited cell expansion, both *in vitro* and *in vivo*. Accordingly, expression of CR7 in CAR T cells exhibited time-limited cell proliferation (34). Additionally, a single incubation with FDA-approved JAK inhibitor Ruxolitinib inhibited dFab_CCR-IL2 signaling.

dFab_CCR is a new architecture for constitutive cytokine receptors. The dFab_CCR architecture is versatile and can be used to induce constitutive signaling from a range of cytokine receptors. Certain dFab_CCRs, for instance those derived from GM-CSF or IL18 receptors, may convey advantages for enhanced CAR T cell function. Future clinical exploration may determine the utility of this and other forms of cytokine signaling, particularly in the setting of adoptive immunotherapy for solid cancer.

## Authors’ contributions

**M. Righi:** Conceptualization, formal analysis, validation, investigation, visualization, methodology, writing–original draft, writing–review and editing. **I. Gannon:** CAR characterisation and Metabolomic Investigation. **M. Robson:** Animal Model Investigation and data analysis. **S. Srivastava:** Animal Model Investigation. **E. Kokalaki:** Animal Model Investigation. **F. Nannini:** RNA sequencing and DNA barcoding investigation. **Y. Bai:** RNA sequencing and data analysis**. C. Allen:** DNA plasmid generation. **J. Sillibourne:** DNA plasmid generation and proofreading. **S. Cordoba:** Conceptualization, Supervision, methodology, project administration,. **S. Thomas:** Visualization, writing–review and editing. **M. Pule:** Conceptualization, resources, data curation, formal analysis, supervision, funding acquisition, validation, visualization, writing–original draft, project administration, writing– review and editing.

## Supporting information

Material and Methods Resources Table

## Acknowledgement

All work was supported by Autolus Therapeutics.

## REFERENCES

1. Kershaw MH, Westwood JA, Darcy PK. Gene-engineered T cells for cancer therapy. Nat Rev Cancer. 2013;13:525–41.

2. Dwyer CJ, Knochelmann HM, Smith AS, Wyatt MM, Rangel Rivera GO, Arhontoulis DC, et al. Fueling Cancer Immunotherapy With Common Gamma Chain Cytokines. Front Immunol [Internet]. Frontiers; 2019 [cited 2020 Aug 2];10. Available from: https://www.frontiersin.org/articles/10.3389/fimmu.2019.00263/full

3. Whiteside TL. The tumor microenvironment and its role in promoting tumor growth. Oncogene. NIH Public Access; 2008;27:5904.

4. Rafiq S, Hackett CS, Brentjens RJ. Engineering strategies to overcome the current roadblocks in CAR T cell therapy. Nat Rev Clin Oncol. 2020;17:147–67.

5. Leonard JP, Sherman ML, Fisher GL, Buchanan LJ, Larsen G, Atkins MB, et al. Effects of Single-Dose Interleukin-12 Exposure on Interleukin-12–Associated Toxicity and Interferon-γ Production. Blood. American Society of Hematology; 1997;90:2541–8.

6. Rosenberg SA, Yannelli JR, Yang JC, Topalian SL, Schwartzentruber DJ, Weber JS, et al. Treatment of patients with metastatic melanoma with autologous tumor-infiltrating lymphocytes and interleukin 2. J Natl Cancer Inst. 1994;86:1159–66.

7. Chen Y, Sun C, Landoni E, Metelitsa L, Dotti G, Savoldo B. Eradication of Neuroblastoma by T Cells Redirected with an Optimized GD2-Specific Chimeric Antigen Receptor and Interleukin-15. Clin Cancer Res. 2019;25:2915–24.

8. Hoyos V, Savoldo B, Quintarelli C, Mahendravada A, Zhang M, Vera J, et al. Engineering CD19-specific T lymphocytes with interleukin-15 and a suicide gene to enhance their anti-lymphoma/leukemia effects and safety. Leukemia. 2010;24:1160–70.

9. Ogawa K, Kawahara M, Nagamune T. Construction of unnatural heterodimeric receptors based on IL-2 and IL-6 receptor subunits. Biotechnology Progress. 2013;29:1512–8.

10. Wang Y, Jiang H, Luo H, Sun Y, Shi B, Sun R, et al. An IL-4/21 Inverted Cytokine Receptor Improving CAR-T Cell Potency in Immunosuppressive Solid-Tumor Microenvironment. Front Immunol [Internet]. Frontiers; 2019 [cited 2020 Aug 19];10. Available from: https://www.frontiersin.org/articles/10.3389/fimmu.2019.01691/full

11. Wilkie S, Burbridge SE, Chiapero-Stanke L, Pereira ACP, Cleary S, van der Stegen SJC, et al. Selective expansion of chimeric antigen receptor-targeted T-cells with potent effector function using interleukin-4. J Biol Chem. 2010;285:25538–44.

12. Hurton LV, Singh H, Najjar AM, Switzer KC, Mi T, Maiti S, et al. Tethered IL-15 augments antitumor activity and promotes a stem-cell memory subset in tumor-specific T cells. Proc Natl Acad Sci U S A. 2016;113:E7788–97.

13. Shum T, Omer B, Tashiro H, Kruse RL, Wagner DL, Parikh K, et al. Constitutive Signaling from an Engineered IL7 Receptor Promotes Durable Tumor Elimination by Tumor-Redirected T Cells. Cancer Discov. 2017;7:1238–47.

14. Straathof K, Flutter B, Wallace R, Jain N, Loka T, Depani S, et al. Antitumor activity without on-target off-tumor toxicity of GD2–chimeric antigen receptor T cells in patients with neuroblastoma. Science Translational Medicine. American Association for the Advancement of Science; 2020;12:eabd6169.

15. Nicholson IC, Lenton KA, Little DJ, Decorso T, Lee FT, Scott AM, et al. Construction and characterisation of a functional CD19 specific single chain Fv fragment for immunotherapy of B lineage leukaemia and lymphoma. Molecular Immunology. 1997;34:1157–65.

16. Philip B, Kokalaki E, Mekkaoui L, Thomas S, Straathof K, Flutter B, et al. A highly compact epitope-based marker/suicide gene for easier and safer T-cell therapy. Blood. American Society of Hematology; 2014;124:1277–87.

17. Thomas S, Straathof K, Himoudi N, Anderson J, Pule M. An Optimized GD2-Targeting Retroviral Cassette for More Potent and Safer Cellular Therapy of Neuroblastoma and Other Cancers. PLOS ONE. 2016;11:e0152196.

18. Hu J, Ge H, Newman M, Liu K. OSA: a fast and accurate alignment tool for RNA-Seq. Bioinformatics. Oxford Academic; 2012;28:1933–4.

19. Love MI, Huber W, Anders S. Moderated estimation of fold change and dispersion for RNA-seq data with DESeq2. Genome Biol [Internet]. 2014 [cited 2020 Apr 24];15. Available from: https://www.ncbi.nlm.nih.gov/pmc/articles/PMC4302049/

20. Nelson BH, Lord JD, Greenberg PD. Cytoplasmic domains of the interleukin-2 receptor beta and gamma chains mediate the signal for T-cell proliferation. Nature. 1994;369:333–6.

21. Usacheva A, Sandoval R, Domanski P, Kotenko SV, Nelms K, Goldsmith MA, et al. Contribution of the Box 1 and Box 2 motifs of cytokine receptors to Jak1 association and activation. J Biol Chem. 2002;277:48220–6.

22. Kenderian SS, Ruella M, Shestova O, Kim MY, Klichinsky M, Chen F, et al. Ruxolitinib Prevents Cytokine Release Syndrome after CART Cell Therapy without Impairing the Anti-Tumor Effect in a Xenograft Model. Blood. 2016;128:652–652.

23. Dinarello CA. Overview of the IL-1 family in innate inflammation and acquired immunity. Immunol Rev. 2018;281:8–27.

24. Vignali DAA, Kuchroo VK. IL-12 Family Cytokines: Immunological Playmakers. Nat Immunol. 2012;13:722–8.

25. Bucks CM, Norton JA, Boesteanu AC, Mueller YM, Katsikis PD. Chronic antigen stimulation alone is sufficient to drive CD8+ T cell exhaustion. J Immunol. 2009;182:6697–708.

26. Lange S, Sand LG, Bell M, Patil SL, Langfitt D, Gottschalk S. A chimeric GM-CSF/IL18 receptor to sustain CAR T-cell function. Cancer Discov. 2021;

27. Klein Geltink RI, Kyle RL, Pearce EL. Unraveling the Complex Interplay Between T Cell Metabolism and Function. Annu Rev Immunol. 2018;36:461–88.

28. Feige MJ, Groscurth S, Marcinowski M, Shimizu Y, Kessler H, Hendershot LM, et al. An Unfolded CH1 Domain Controls the Assembly and Secretion of IgG Antibodies. Molecular Cell. 2009;34:569–79.

29. Röthlisberger D, Honegger A, Plückthun A. Domain interactions in the Fab fragment: a comparative evaluation of the single-chain Fv and Fab format engineered with variable domains of different stability. J Mol Biol. 2005;347:773–89.

30. Hust M, Jostock T, Menzel C, Voedisch B, Mohr A, Brenneis M, et al. Single chain Fab (scFab) fragment. BMC Biotechnology. 2007;7:14.

31. Klein C, Schaefer W, Regula JT. The use of CrossMAb technology for the generation of bi- and multispecific antibodies. MAbs. 2016;8:1010–20.

32. Cendrowski J, Mamińska A, Miaczynska M. Endocytic regulation of cytokine receptor signaling. Cytokine & Growth Factor Reviews. 2016;32:63–73.

33. Moraga I, Wernig G, Wilmes S, Gryshkova V, Richter CP, Hong W-J, et al. Tuning cytokine receptor signaling by re-orienting dimer geometry with surrogate ligands. Cell. 2015;160:1196– 208.

34. Shum T, Omer B, Tashiro H, Kruse RL, Wagner DL, Parikh K, et al. Constitutive Signaling from an Engineered IL7 Receptor Promotes Durable Tumor Elimination by Tumor-Redirected T Cells. Cancer Discov. 2017;7:1238–47.

35. Nakamura Y, Russell SM, Mess SA, Friedmann M, Erdos M, Francois C, et al. Heterodimerization of the IL-2 receptor beta- and gamma-chain cytoplasmic domains is required for signalling. Nature. 1994;369:330–3.

36. Floss DM, Schönberg M, Franke M, Horstmeier FC, Engelowski E, Schneider A, et al. IL-6/IL-12 Cytokine Receptor Shuffling of Extra- and Intracellular Domains Reveals Canonical STAT Activation via Synthetic IL-35 and IL-39 Signaling. Sci Rep [Internet]. 2017 [cited 2020 Apr 24];7. Available from: https://www.ncbi.nlm.nih.gov/pmc/articles/PMC5680241/

37. Dwyer CJ, Knochelmann HM, Smith AS, Wyatt MM, Rangel Rivera GO, Arhontoulis DC, et al. Fueling Cancer Immunotherapy With Common Gamma Chain Cytokines. Front Immunol [Internet]. 2019 [cited 2020 Sep 2];10. Available from: https://www.ncbi.nlm.nih.gov/pmc/articles/PMC6391336/

38. Locke FL, Ghobadi A, Jacobson CA, Miklos DB, Lekakis LJ, Oluwole OO, et al. Long-term safety and activity of axicabtagene ciloleucel in refractory large B-cell lymphoma (ZUMA-1): a single-arm, multicentre, phase 1–2 trial. The Lancet Oncology. 2019;20:31–42.

39. Maude SL, Laetsch TW, Buechner J, Rives S, Boyer M, Bittencourt H, et al. Tisagenlecleucel in Children and Young Adults with B-Cell Lymphoblastic Leukemia. New England Journal of Medicine. 2018;378:439–48.

40. Raje N, Berdeja J, Lin Y, Siegel D, Jagannath S, Madduri D, et al. Anti-BCMA CAR T-Cell Therapy bb2121 in Relapsed or Refractory Multiple Myeloma. New England Journal of Medicine. Massachusetts Medical Society; 2019;380:1726–37.

41. Nakanishi K. Unique Action of Interleukin-18 on T Cells and Other Immune Cells. Front Immunol [Internet]. Frontiers; 2018 [cited 2020 May 1];9. Available from: https://www.frontiersin.org/articles/10.3389/fimmu.2018.00763/full

42. Kunert A, Chmielewski M, Wijers R, Berrevoets C, Abken H, Debets R. Intra-tumoral production of IL18, but not IL12, by TCR-engineered T cells is non-toxic and counteracts immune evasion of solid tumors. OncoImmunology. 2018;7:e1378842.

43. Chmielewski M, Abken H. CAR T Cells Releasing IL-18 Convert to T-Bethigh FoxO1low Effectors that Exhibit Augmented Activity against Advanced Solid Tumors. Cell Reports. Elsevier; 2017;21:3205–19.

44. Hu B, Ren J, Luo Y, Keith B, Young RM, Scholler J, et al. Augmentation of Antitumor Immunity by Human and Mouse CAR T Cells Secreting IL-18. Cell Rep. 2017;20:3025–33.

45. Fields JK, Günther S, Sundberg EJ. Structural Basis of IL-1 Family Cytokine Signaling. Front Immunol [Internet]. 2019 [cited 2020 Apr 22];10. Available from: https://www.ncbi.nlm.nih.gov/pmc/articles/PMC6596353/

46. Warner N, Núñez G. MyD88: A Critical Adaptor Protein in Innate Immunity Signal Transduction. The Journal of Immunology. American Association of Immunologists; 2013;190:3–4.

47. Mata M, Gerken C, Nguyen P, Krenciute G, Spencer DM, Gottschalk S. Inducible Activation of MyD88 and CD40 in CAR T Cells Results in Controllable and Potent Antitumor Activity in Preclinical Solid Tumor Models. Cancer Discov. 2017;7:1306–19.

48. Prinzing B, Schreiner P, Bell M, Fan Y, Krenciute G, Gottschalk S. MyD88/CD40 signaling retains CAR T cells in a less differentiated state. JCI Insight. 2020;5:136093.

49. Glienke W, Dragon AC, Zimmermann K, Martyniszyn-Eiben A, Mertens M, Abken H, et al. GMP-Compliant Manufacturing of TRUCKs: CAR T Cells targeting GD2 and Releasing Inducible IL-18. Front Immunol. 2022;13:839783.

50. Blokon-Kogan D, Levi-Mann M, Malka-Levy L, Itzhaki O, Besser MJ, Shiftan Y, et al. Membrane anchored IL-18 linked to constitutively active TLR4 and CD40 improves human T cell antitumor capacities for adoptive cell therapy. J Immunother Cancer. BMJ Specialist Journals; 2022;10:e001544.

51. Zhou T, Damsky W, Weizman O-E, McGeary MK, Hartmann KP, Rosen CE, et al. IL-18BP is a secreted immune checkpoint and barrier to IL-18 immunotherapy. Nature. 2020;583:609–14.

52. Cavazzana-Calvo M, Hacein-Bey S, de Saint Basile G, Gross F, Yvon E, Nusbaum P, et al. Gene therapy of human severe combined immunodeficiency (SCID)-X1 disease. Science. 2000;288:669–72.

53. Hacein-Bey-Abina S, Von Kalle C, Schmidt M, McCormack MP, Wulffraat N, Leboulch P, et al. LMO2-associated clonal T cell proliferation in two patients after gene therapy for SCID-X1. Science. 2003;302:415–9.

54. Ruggero K, Al-Assar O, Chambers JS, Codrington R, Brend T, Rabbitts TH. LMO2 and IL2RG synergize in thymocytes to mimic the evolution of SCID-X1 gene therapy-associated T-cell leukaemia. Leukemia. 2016;30:1959–62.

55. Zhou S, Mody D, DeRavin SS, Hauer J, Lu T, Ma Z, et al. A self-inactivating lentiviral vector for SCID-X1 gene therapy that does not activate LMO2 expression in human T cells. Blood. 2010;116:900–8.

