## Supplementary material for "Enhancing CAR T cell therapy using Fab-Based Constitutively Heterodimeric Cytokine Receptors": Material and Methods Resources Table

**Key Resources Table**

| **Reagent** | **Source** | **Identifier** |
| --- | --- | --- |
| **Antibody** | | |
| Anti-GD2CAR idiotype | This Lab | N/A |
| Anti-CD19CAR idiotype | This Lab | N/A |
| hCCR7 PE (clone REA546) | Miltenyi | Cat# 130-119-583, RRID:AB_2655953 |
| hCD8 VioBlue (clone REA734) | Miltenyi | Cat# 130-110-683  RRID:AB_10828909 |
| hCD3 VioGreen (clone REA613) | Miltenyi | Cat# 130-113-142, RRID:AB_2657067 |
| hCD45RA APCVio770 (clone REA1047) | Miltenyi | Cat# 130-117-747, RRID:AB_2658333 |
| hLAG3 VioBright515 (clone REA351) | Miltenyi | Cat# 130-120-012, RRID:AB_2656417 |
| hPD1 PE (clone REA1165) | Miltenyi | Cat# 130-120-382, RRID:AB_2661364 |
| hTIM3 (CD366) PEVio770 | Miltenyi | Cat# 130-121-334, RRID:AB_2784166 |
| hKLRG1 APCVio770 (clone REA635) | Miltenyi | Cat# 130-120-423, RRID:AB_2727835 |
| hREA control (S) PE (clone REA293) | Miltenyi | Cat# 130-113-438, RRID:AB_2661690 |
| hREA control (S) VioBright515 (clone REA293) | Miltenyi | Cat# 130-113-445, RRID:AB_2661711 |
| hREA control (S) PEVio770 (clone REA293) | Miltenyi | Cat# 130-113-440, RRID:AB_2661707) |
| hREA control (S) APCVio770 (clone REA293) | Miltenyi | Cat# 130-113-435, RRID:AB_2661696 |
| hCD69 FITC (clone FN50) | Biolegend | Cat# 310904,  RRID: AB_314838 |
| hCD25 Pacific-Blue (clone BC96) | Biolegend | Cat# 302627,  RRID: AB_493649 |
| hCD95 APC-Cy7 (clone DX2) | Biolegend | Cat# 305636,  RRID: AB_2566108 |
| hCD3 PE-Cy7 (clone SK7) | Biolegend | Cat# 317334,  RRID: AB_2043994 |
| hCD2 APC (clone RPA-2.10) | Biolegend | Cat# 300214,  RRID: AB_10900259 |
| hGD2 PE (clone 14g2a) | Biolegend | Cat# 357304,  RRID: AB_2561884 |
| hCD34 FITC (Clone QBEnd10) | R&D system | Cat# [FAB7227G](https://www.rndsystems.com/products/human-cd34-alexa-fluor-488-conjugated-antibody-qbend10_fab7227g), RRID:AB_10973657 |
| rat CD90/mouse CD90.1 (Thy-1.1) Pacific-blue (Clone OX-7) | Biolegend | Cat# 202522, RRID:AB_1595477 |
| mCD3-APC (clone 17A2) | Biolegend | Cat# 100236, RRID:AB_2561456 |
| mCD4-FITC (clone RM4-4) | Biolegend | Cat# 116004, RRID:AB_313688 |
| mCD45-PerCp-Cy5.5 (clone 30-F11) | Biolegend | Cat# 103132, RIID: AB_893340 |
| mCD11b-PE/Cyanine7 (clone M1/70) | Biolegend | Cat# 101216, RRID, AB_312799 |
| FITC anti-mouse CD11c | Biolegend | 117306, RRID  AB_313775 |
| PE anti-mouse F4/80 | Biolegend | 157304, RRID  AB_2832547 |
| PE anti-mouse/human CD44 | Biolegend | 163609, RRID  AB_2924492 |
| Brilliant Violet 510™ anti-mouse CD62L | Biolegend | 104441, RRID  AB_2561537 |
| Human Kappa Light Chain Biotin (Clone L1C1) | Invitrogen | Cat# MA5-12114, RRID:AB_10984749 |
| Streptavidin PE | Biolegend | Cat# 405204 |
| DyLight™ 488 AffiniPure Mouse Anti-Human IgG, F(ab')₂ fragment specific | JacksonImmuno | Cat# 209-485-097, RRID:AB_2339126 |
| R-Phycoerythrin AffiniPure Fab Fragment Goat Anti-Rabbit IgG, Fc fragment specific | JacksonImmuno | Cat# 111-117-008, RRID:AB_2632461 |
| Goat anti-Rabbit IgG APC | ThermoFisher | Cat# A-10931, RRID:AB_141369 |
| SYTOX™ Blue Dead Cell Stain | ThermoFisher | Cat# S10274 |
| eBioscience™ Fixable Viability Dye eFluor™ 450 | ThermoFisher | Cat# 65-0863-18 |
| eBioscience™ Fixable Viability Dye eFluor™ 780 | ThermoFisher | Cat# 65-0865-14 |
| Human TruStain FcX™ (Fc Receptor Blocking Solution) | Biolegend | Cat# 422301  RRID: AB_2818986 |
| TruStain FcX™ PLUS (anti-mouse CD16/32) Antibody (clone S17011E) | Biolegend | Cat# 156604,  RRID: AB_2783138 |
| rabbit IgG, human GAPDH (clone D16H11) | Cell Signalling Technology | Cat# 5174S, RRID:AB_10622025 |
| rabbit IgG, HRP-linked Antibody | Cell Signalling Technology | Cat# 7074P2, RRID:AB_2099233 |
| rabbit IgG, human Phospho-STAT1 (Tyr701) | Cell Signalling Technology | Cat# 7649S, RRID:AB_10950970 |
| rabbit IgG, human Phospho-STAT3 (Tyr705) | Cell Signalling Technology | Cat# 9134S, RRID:AB_331589 |
| rabbit IgG, human Phospho-STAT4 (Ser721) Polyclonal Antibody | ThermoFisher | Cat# PA5-105861, RRID:AB_2817260 |
| rabbit IgG, human Phospho-STAT5 alpha (Tyr694) Polyclonal Antibody (Clone ZyAL) | ThermoFisher | Cat# 71-6900, RRID:AB_88412 |
| rabbit IgG, human, Phospho-p44/42 MAPK (Erk1/2) (Thr202/Tyr204) (clone D13.14.4E) | Cell Signalling Technology | Cat# 4370S, RRID:AB_2315112 |
| **Biological samples** | | |
| Buffy coats from healthy donors | NHS Blood and Transplant. NHSBT, London, UK | NA |
| **Chemicals, peptides, recombinant proteins and media** | | |
| Ruxolitinib | StemCell | Cat# 73402 |
| Retronectin | Takara/Clontech | Cat# T100B |
| Human IL2 | GenScript | Cat# Z00368 |
| GeneJuice | Merck Millipore | Cat# 70967 |
| Concavalin-A | Sigma Aldrich | Cat# C0412-5MG |
| Mouse IL7 | Miltenyi | Cat# 130-094-066 |
| β-mercaptoethanol | Bio-Rad | Cat# 1610710 |
| Fetal Bovine Serum (South America) | BIOSERA | Cat# FB-1101/100 |
| HEPES (1 M) | Gibco™ | Cat# 15630106 |
| GlutaMAX™ Supplement | Gibco™ | Cat# 35050061 |
| CD3 Antibody, anti-human, pure-functional grade | Miltenyi | Cat# 130-093-387 |
| CD28 Antibody, anti-human, pure-functional grade | Miltenyi | Cat# 130-093-375 |
| Ficoll-Paque™ PREMIUM | GE Healthcare | Cat# 17-5442-02 |
| IMDM | Gibco™ | Cat# 12440053 |
| RPMI 1640 | Gibco™ | Cat# 21875034 |
| EasySep™ Buffer | STEMCELL™ | Cat# 20144 |
| Bovine Serum Albumin | Sigma-Aldrich | Cat# A9418-500G |
| 10x Tris/Glycine/SDS | Bio-Rad | Cat# 1610772 |
| Clarity Western ECL Substrate | Bio-Rad | Cat# 1705061 |
| Pierce™ 20X TBS Tween™ 20 Buffer | Thermo Scientific™ | Cat# 28360 |
| Pierce™ 20X PBS Tween™ 20 Buffer | Thermo Scientific™ | Cat# 28352 |
| Gibco™ PBS, pH 7.4 | Gibco™ | Cat# 10010015 |
| Trypsin-EDTA (0.25%), phenol red | Gibco™ | Cat# 25200056 |
| ACK Lysing Buffer | Gibco™ | Cat# A1049201 |
| Hanks' Balanced Salt Solution (1X), with calcium, with magnesium, without phenol red | Thermo Scientific™ | Cat# J67763.K2 |
| Protease and Phosphatase Inhibitor Cocktail | Abcam | Cat# ab201119 |
| 100 mL RIPA Lysis Buffer, 10X | Millipore | Cat# 20-188 |
| VivoGlo™ Luciferin, In Vivo Grade | Promega | P1043 |
| **Critical commercial assays and kits** | | |
| LEGENDplex™ HU Proinflam. Chemokine Panel 1 (13-plex) w/VbP | Biolegend | Cat# 740985 |
| LEGENDplex™ HU Th Cytokine Panel (12-plex) w/ VbP V02 | Biolegend | Cat# 741028 |
| LEGENDplex™ Mouse Proinflammatory Chemokine Panel (13-plex) with V-bottom Plate | Biolegend | Cat# 740451 |
| LEGENDplex™ MU Th Cytokine Panel (12-plex) w/ VbP V03 | Biolegend | Cat# 741044 |
| ELISA MAX™ Deluxe Set Mouse IFN-γ | Biolegend | Cat# 430804 |
| ELISA MAX™ Deluxe Set Human IFN-γ | Biolegend | Cat# 430104 |
| ELISA MAX™ Deluxe Set Human IL-2 | Biolegend | Cat# 431804 |
| nCounter® CAR-T Characterization Panel | NanoString | NA |
| Tumor Dissociation Kit, murine | Miltenyi | Cat# 130-096-730 |
| CD34 MicroBead Kit, human | Miltenyi | Cat# 130-046-702 |
| PureLink RNA Mini Kit | Thermo Scientific™ | Cat# 12183018A |
| PureLink™ Genomic DNA Mini Kit | Thermo Scientific™ | Cat# K182002 |
| Seahorse XF Cell Mito Stress Test Kit RUO | Agilent | Cat# 103015-100 |
| BD Cytofix/Cytoperm™ Fixation/Permeabilization Solution Kit | BD Bioscience | Cat# BD 554714 |
| **Experimental models: Cell lines** | | |
| CT26 | ATCC | Cat# CRL-2638, RRID:CVCL_7256 |
| B16F10 | ATCC | Cat# CRL-6475, RRID:CVCL_0159 |
| HEK293T | ATCC | Cat# CRL-3216, RRID:CVCL_0063 |
| CHLA-255 | COG Cell Culture Core + Xenograft Repository, Texas Tech Uni HSC Cancer Center | RRID:CVCL_AQ27 |
| SupT1 | ATCC | Cat# CRL-1942, RRID:CVCL_1714 |
| SKOV3 | ATCC | Cat# HTB-77, RRID:CVCL_0532 |
| NALM6 | ATCC | RRID:CVCL_0092 |
| Phoenix eco | ATCC | Cat# SD-3444, RRID:CVCL_H717 |
| CT26-GD2 | This lab | N/A |
| B16F10-GD2 | This lab | N/A |
| SupT1-GD2 | This lab | N/A |
| SKOV3-GD2 | This lab | N/A |
| SKOV3-CD19 | This lab | N/A |
| NALM6-FFluc | This lab | N/A |
| NALM6-CD19KO | This lab | N/A |
| CHLA-FFluc | This lab | N/A |
| **Experimental models: Mouse** | | |
| NOD.Cg-Prkdcscid Il2rgtm1Wjl/SzJ | Jackson Lab | Strain #:005557  RRID:IMSR_JAX:005557 |
| BALB/cAnNCrl | Charles River | Strain Code 028 |
| C57BL/6NCrl | Charles River | Strain Code 027 |
| **Oligonucleotides** | | |
| Barcode Fwd primer  ACACTCTTTCCCTACACGACGCTCTTCCGATCTCCGCCAACGCGGTCGCAC | This Lab | NA |
| Barcode Rev primer  GACTGGAGTTCAGACGTGTGCTCTTCCGATCTGATGAGAACAGTATCGATTAGGGTTGACGGC | This Lab | NA |
| **Software and algorithms** | | |
| Flowjo v10 | FlowJo, LLC | RRID:SCR_008520 |
| GraphPad Prism | GraphPad Software Inc. | RRID:SCR_002798 |
| nSolver Analysis Software v4.0 | NanoString | RRID:SCR_003420 |
| nSolver Advanced Analysis v2.0 | NanoString | N/A |
| Seahorse Wave | Agilent | RRID:SCR_014526 |
| **Recombinant DNA** | | |
| **Human dFab_CCR vectors** | | |
| **Cytokine receptor** | **Constant light Kappa chain (P01834)** | **Constant Heavy 1 chain (P01857)** |
| dFab_CCR-IL2 | Common γ chain (P31785) | IL2 receptor β chain (P14784) |
| dFab_CCR-IL7 | Common γ chain (P31785) | IL7 receptor α chain (P16871) |
| dFab_CCR-IL4 | Common γ chain (P31785) | IL4 receptor α chain (P24394) |
| dFab_CCR-IL9 | Common γ chain (P31785) | IL9 receptor α chain (Q01113) |
| dFab_CCR-IL21 | Common γ chain (P31785) | IL21 receptor α chain (Q9HBE5) |
| dFab_CCR-TSLP | Cytokine receptor-like factor 2 (Q9HC73) | IL7 receptor α chain (P16871) |
| dFab_CCR-IL3 | Common β chain (P32927) | IL3 receptor α chain (P26951) |
| dFab_CCR-IL5 | Common β chain (P32927) | IL5 receptor α chain (Q01344) |
| dFab_CCR-GMCSF | Common β chain (P32927) | GMCSF receptor α chain (P15509) |
| dFab_CCR-IL12 | IL12 receptor β1 (P42701) | IL12 receptor β2 (Q99665) |
| dFab_CCR-IL23 | IL23 receptor α (Q9NPF7) | IL12 receptor β1 (P42701) |
| dFab_CCR-IL27 | IL27 receptor β (Q6UWB1) | GP130 (P40189) |
| dFab_CCR-IL10 | IL10 receptor β (Q08334) | IL10 receptor α (Q13651) |
| dFab_CCR-IL1 | IL1 receptor 1 (P14778) | IL1 receptor 2 (P27930) |
| dFab_CCR-IL18 | IL18 receptor 1 (Q13478) | IL18 receptor AP (O95256) |
| dFab_CCR-IL33 | IL-1 receptor-like 1 (P14778) | IL1 receptor AP (Q9NPH3) |
| dFab_CCR-IL17A | IL17 receptor A (Q96F46) | IL17 receptor B (Q9NRM6) |
| dFab_CCR-IL17E | IL17 receptor A (Q96F46) | IL17 receptor C (Q8NAC3) |
| Switched dFab_CCR-IL2 | IL2 receptor β chain (P14784) | Common γ chain (P31785) |
| **Murine dFab_CCR vectors** | | |
| **Cytokine receptor** | **Constant light Kappa chain (P01834)** | **Constant Heavy 1 chain (P01857)** |
| dFab_CCR-IL2mu | Common γ chain (P34902) | IL2 receptor β chain (P16297) |
| dFab_CCR-IL7mu | Common γ chain (P34902) | IL7 receptor α chain (P16872) |
| dFab_CCR-GMCSFmu | Common β chain (P26955) | GMCSF receptor α chain (Q00941) |
| dFab_CCR-IL12mu | IL12 receptor β1 (Q60837) | IL12 receptor β2 (P97378) |
| dFAb_CCR-IL10mu | IL10 receptor β (Q61190) | IL10 receptor α (Q61727) |
| dFAb_CCR-IL18mu | IL18 receptor 1 (Q61098) | IL18 receptor AP (Q9Z2B1) |
| **CARs** | | |
| CAR.GD2.huk666VLVH.8aHTM.BBζ | This manuscript | N/A |
| CAR.CD19.fmc63VLVH.8aHTM.BBζ | This manuscript | N/A |
| CAR.PSMA.10C11VLVH.8aHTM.BBζ | This manuscript | N/A |
| muCAR.GD2.muk666VLVH.mu8aHTM.muBBmuζ | This manuscript | N/A |
