## Supplementary Figure 1-25 for "Enhancing CAR T cell therapy using Fab-Based Constitutively Heterodimeric Cytokine Receptors"

### Supplementary Figures for Righi et al.

Supplementary Figure S1. dFab_CCR-IL2 receptor hetero-pairs orientation influence dFab_CCR-IL2 signal output. 3
Supplementary Figure S2. dFab_CCR-IL2 receptor endodomain signalling motifs influence dFab_CCR-IL2 signal potency. 4
Supplementary Figure S3. JAK kinase inhibitor Ruxolitinib tunes dFab_CCR-IL2 receptor signal. 6
Supplementary Figure S4. dFab_CCR-IL2 and native IL2 receptor have comparable signal and transcriptome quality. 8
Supplementary Figure S5. dFab_CCR expression profile in primary T cells. 10
Supplementary Figure S6. dFab_CCRs function primary T cells. 12
Supplementary Figure S7. dFab_CCR-IL2 maintains functionality when co-expressed with a second generation GD2 CAR in a bi-cistronic vector. 14
Supplementary Figure S8. dFab_CCRs co-expression protect GD2 CAR T cells cytokine secretion. 16
Supplementary Figure S9. Characterisation of selected dFab_CCRs co-expressed with a second generation 41bbζ-CD19 CAR. 18
Supplementary Figure S10. dFab_CCRs co-expression do not alter CAR specificity. 20
Supplementary Figure S11. Differential effect on GD2 CAR functionality of the remaining dFab_CCRs library receptor upon antigen chronic stimulation. 22
Supplementary Figure S12. Selected dFab_CCRs library receptors protect GD2 CAR T cells cytokine secretion upon antigen chronic stimulation. 24
Supplementary Figure S13. Chronic Antigen exposure effects on CD19 CAR T cells co-expressing selected dFab_CCRs. 25
Supplementary Figure S14. Differential effect of the remaining dFab_CCRs library receptors on exhaustion and memory phenotype after chronic antigen exposure. 27
Supplementary Figure S15. Differential effect of selected dFab_CCRs library receptors on exhaustion and memory phenotype after CD19 CAR T cellschronic antigen exposure. 29
Supplementary Figure S16. dFab_CCR DNA barcoded dFab_CCR library deconvolute individual dFab_CCR functionality. 31
Supplementary Figure S17. Transcriptomic profile of GD2 CAR T cells co-expressing a selection of dFab_CCRs. 33
Supplementary Figure S18. Chemokines and cytokines secretion profile of GD2 and CD19 CAR T cells co-expressing a selection of dFab_CCRs. 35
Supplementary Figure S19. Metabolomic profile of GD2 and CD19 CAR T cells co-expressing a selection of dFab_CCRs. 37
Supplementary Figure S20. dFab_CCR co-expression augment CD19 CAR T cells efficacy in an establishes ALL NALM6 xenograph model. 39
Supplementary Figure S21. Bioluminescence imaging of the established ALL NALM6 xenograph model. 41

#### SUPPLEMENTARY FIGURE S1

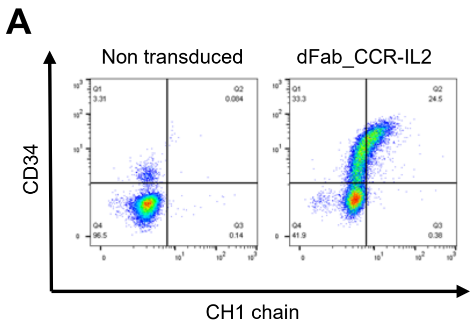

**Supplementary Figure S1. dFab_CCR-IL2 receptor hetero-pairs expression.**

**(A)** Flow-cytometric analysis of the constant CH1 chain expression in dFab_CCR-IL2 transduced T cell, after fixation/permeabilization.

#### SUPPLEMENTARY FIGURE S2

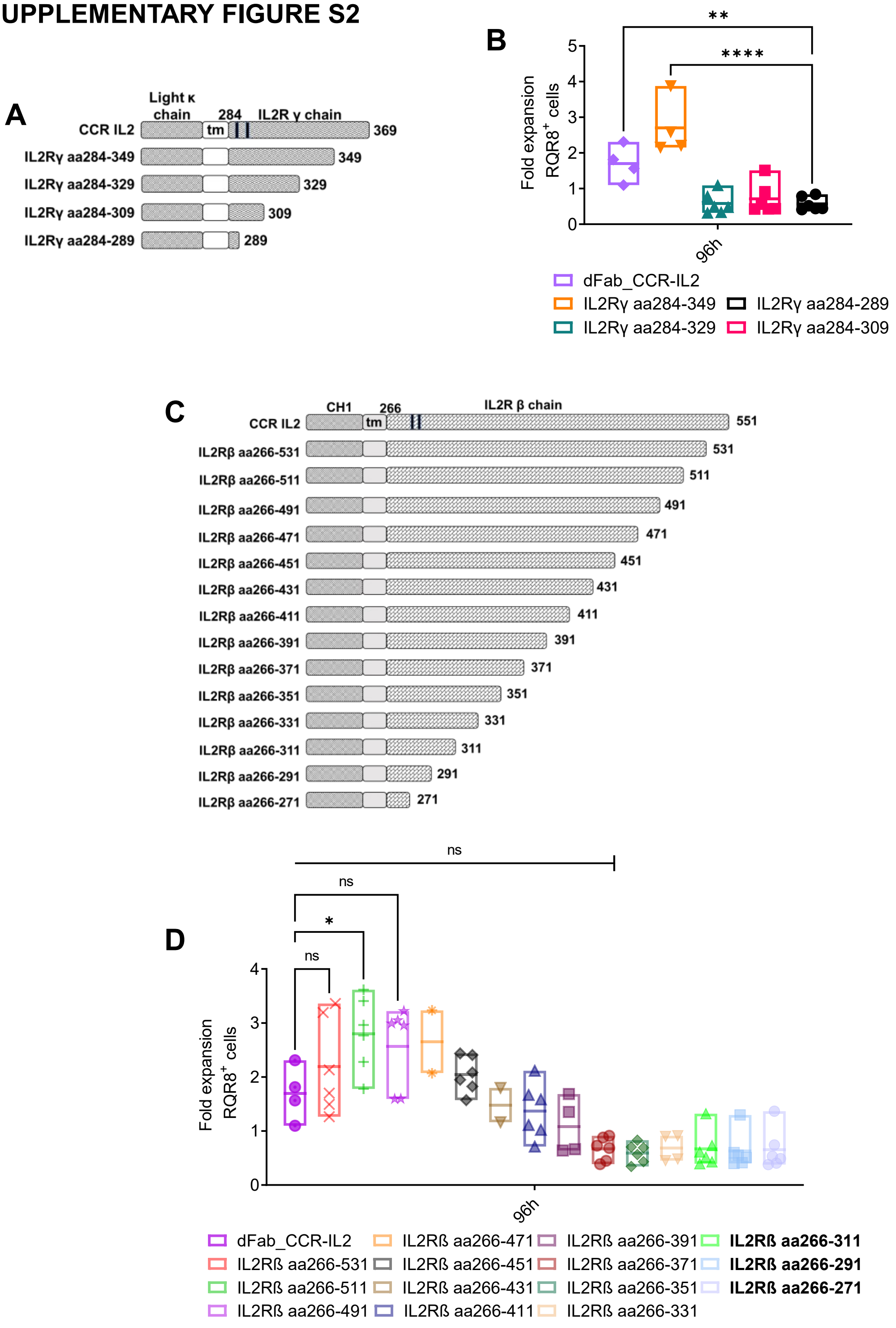

**Supplementary Figure S2. dFab_CCR-IL2 receptor endodomain signalling motifs influence dFab_CCR-IL2 signal potency.**

**(A)** Schematic of the polypeptide sequence of the dFab_CCR CγC truncations. Each truncation was expressed with the full length IL2Rβ/CH1 chain. **(B)** Quantitated *in vitro* proliferation of each dFab_CCR CγC truncations, expressed as $RQR8^+$ / $CD3^+$ T cells fold expansion (n=4, one-way ANOVA). **(C)** Schematic of the polypeptide sequence of the dFab_CCR IL2Rβ truncations. Each truncation was expressed with the full length CγC/CL chain. **(D)** Quantitated *in vitro* proliferation of the series of dFab_CCR IL2Rβ truncations, expressed as $RQR8^+$ / $CD3^+$ T cells fold expansion (n=4, one-way ANOVA). All data are presented as mean ± SEM.

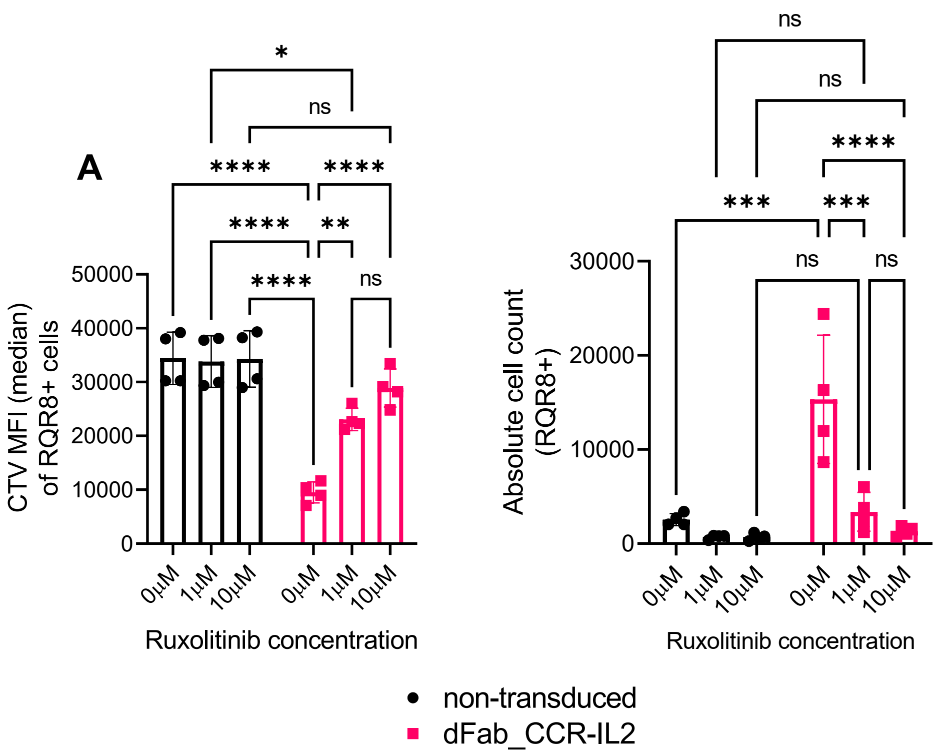

**Supplementary Figure S3. JAK kinase inhibitor Ruxolitinib tunes dFab_CCR-IL2 receptor signal.**

**A)** Sensitivity of dFab_CCR-IL2 engineer T cell to Ruxolitinib. T cell were pre-treated with increasing concentration of Ruxolitinib for 4h. T cells were cultured for 96 hours in cytokine starvation condition. Proliferation expressed as CTV MFI dilution (left), and absolute count (right) of $RQR8^{+}$ / $CD3^{+}$ T cells. (n=4, two-way ANOVA). All data are presented as mean ± SEM.

#### SUPPLEMENTARY FIGURE S4

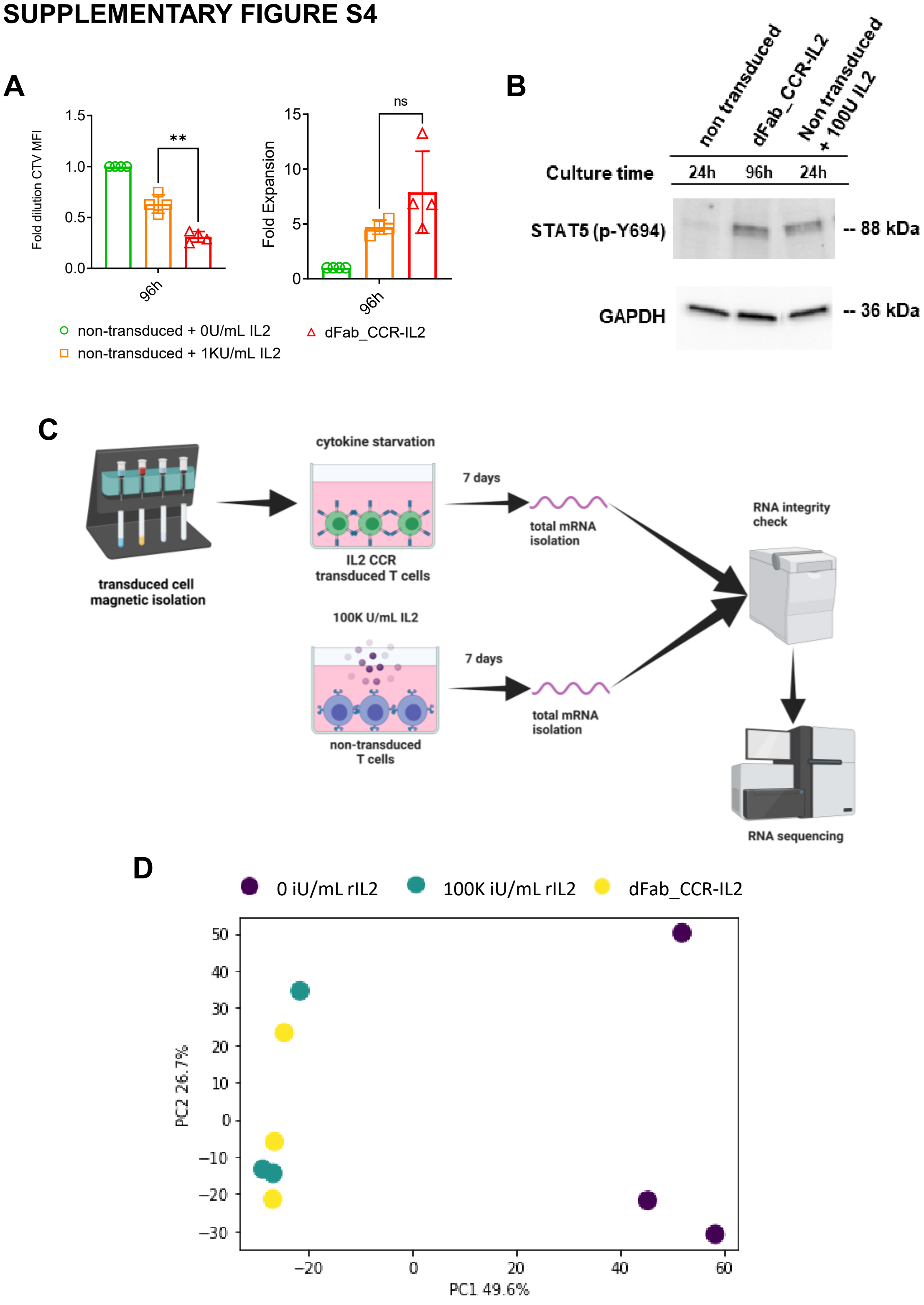

**Supplementary Figure S4. dFab_CCR-IL2 and native IL2 receptor have comparable signal and transcriptome quality.**

**(A)** Comparison of dFab_CCR-IL2 and natural IL2 receptor signal. dFab_CCR-IL2 T cells were cytokine starved for 96 hours. Non transduced T cells were cultured for 96 hours with or without 100000 iU/mL of IL2 for 96 hours. Fold CTV dilution and fold expansion were calculated normalizing the value with the non-transduced T cells in absence of IL2 (n=4, one-way ANOVA). (**B)** Western blot analysis of *in vitro* STAT5-Y694 phosphorylation. Non transduced T cells were culture in presence or absence of 100 IU/mL of IL2 for 24 hours. dFab_CCR-IL2 T cells were cultured in cytokine starvation for 96 hours. GAPDH was used as loading control. Data are representative of three independent experiments. **(C)** Schematic of the total mRNA sequencing experiments. dFab_CCR-IL2 transduced T cell were magnetically isolated using CD34 microbeads and subjected to 7 days of cytokine starvation. T cell were incubated for 7 days with 100000 IU/mL of IL2 or without IL2. IL2 was replenished after 4 days. Total mRNA was isolated and NGS sequenced. **(D)** PCA plot representing the differences between 7 days with 100000 IU/mL of IL2, without IL2 or dFab_CCR-IL2 expression.

#### SUPPLEMENTARY FIGURE S5

**A**

*Human CCR vectors*

| *Cytokine receptor* | **Constant light Kappa chain** | **Constant Heavy 1 chain** |
|---|---|---|
| *Fab_CCR-IL2* | Common γ chain | IL2 receptor βchain |
| *Fab_CCR-IL7* | Common γ chain | IL7 receptor α chain |
| *Fab_CCR-IL4* | Common γ chain | IL4 receptor α chain |
| *Fab_CCR-IL9* | Common γ chain | IL9 receptor α chain |
| *Fab_CCR-IL21* | Common γ chain | IL21 receptor α chain |
| *Fab_CCR-TSLP* | Cytokine receptor-like factor 2 | IL7 receptor α chain |
| *Fab_CCR-IL3* | Common β chain | IL3 receptor α chain |
| *Fab_CCR-IL5* | Common β chain | IL5 receptor α chain |
| *Fab_CCR-GMCSF* | Common β chain | GMCSF receptor α chain |
| *Fab_CCR-IL12* | IL12 receptor β 1 | IL12 receptor β 2 |
| *Fab_CCR-IL23* | IL23 receptor α | IL12 receptor β 1 |
| *Fab_CCR-IL27* | IL27 receptor β | GP130 |
| *Fab_CCR-IL10* | IL10 receptor β 2 | IL10 receptor α |
| *Fab_CCR-IL1* | IL1 receptor 1 | IL1 receptor AP |
| *Fab_CCR-IL18* | IL18 receptor 1 | IL18 receptor AP |
| *Fab_CCR-IL33* | Interleukin-1 receptor-like 1 | IL1 receptor AP |
| *Fab_CCR-IL17A* | IL17 receptor A | IL17 receptor B |
| *Fab_CCR-IL17E* | IL17 receptor A | IL17 receptor C |

**B**

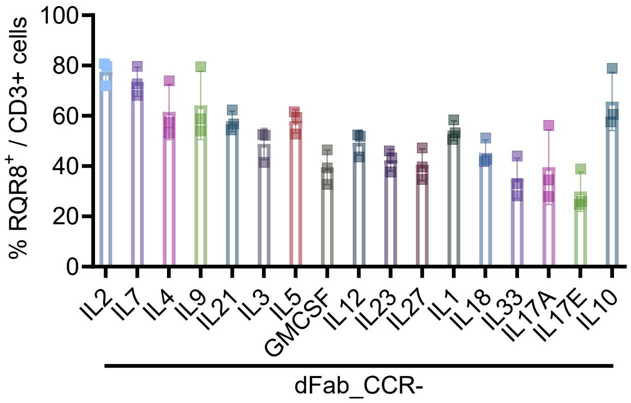

**C**

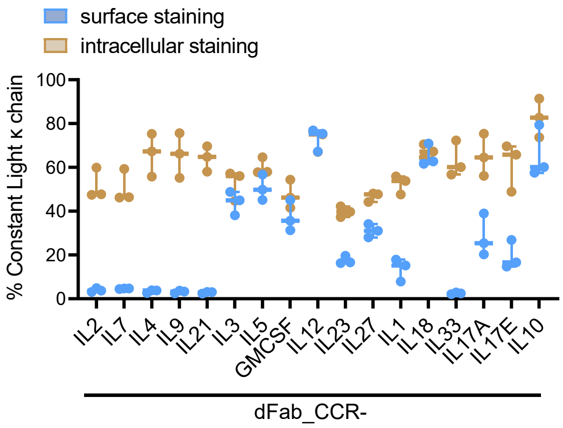

**D**

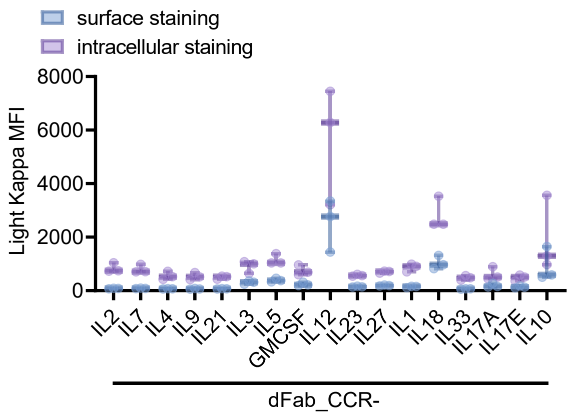

**Supplementary Figure S5. dFab_CCR expression profile in primary T cells.**

**(A)** List of dFab_CCRs cytokine receptor pairs utilized in Figure 2. **(B)** Flow cytometry analysis of dFab_CCRs T cells transduction efficiency as measured by CD34 staining of RQR8 marker gene. These cells were used for the experiments in figure 2. **(C)** Histograms representing the percentage of expression of light kappa chain based on surface staining (blue) and intracellular staining (yellow), (n = 4) **(D)** Histograms representing the dFab_CCRs density expressed of MFI of light kappa chain based on surface staining (blue) and intracellular staining (purple), (n = 4), All data are presented as mean ± SEM.

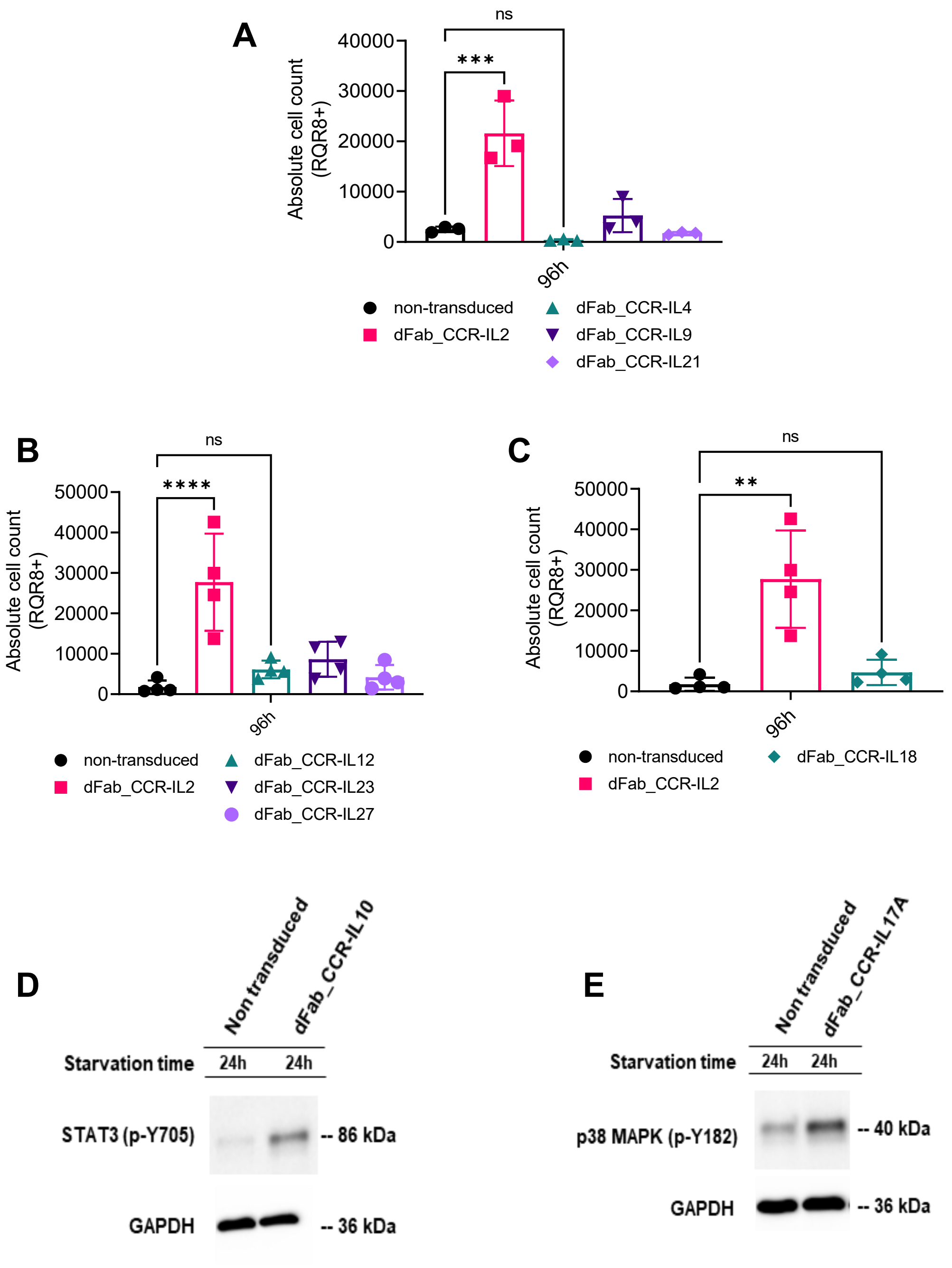

**Supplementary Figure S6. dFab_CCRs function primary T cells.**

**(A, B, C)** Quantification of *in vitro* proliferation after 96h of cytokine starvation of T cells engineered with IL4, IL9, IL21 (A), IL2, IL12, IL23, IL27 (B), IL18 (C) dFab_CCRs, expressed as absolute cell count of RQR8$^{+}$ / CD3$^{+}$ T cells (n=4, one-way ANOVA). **(D)** Immunoblot analysis of *in vitro* STAT3 phosphorylation, monitoring site-specific phosphorylation at Y705. Non-transduced T cells and dFab_CCR-IL10 T cells were cultured in cytokine starvation for 24 hours. GAPDH was used as loading control. Data are representative of three independent experiments. **(E)** Immunoblot analysis of in vitro P38/MAPK phosphorylation, monitoring site-specific phosphorylation at Y182. Non-transduced and dFab_CCR-IL17A transduced T cells were cultured in cytokine starvation for 24 hours. GAPDH was used as loading control. Data are representative of three independent experiments. All data are presented as mean ± SEM.

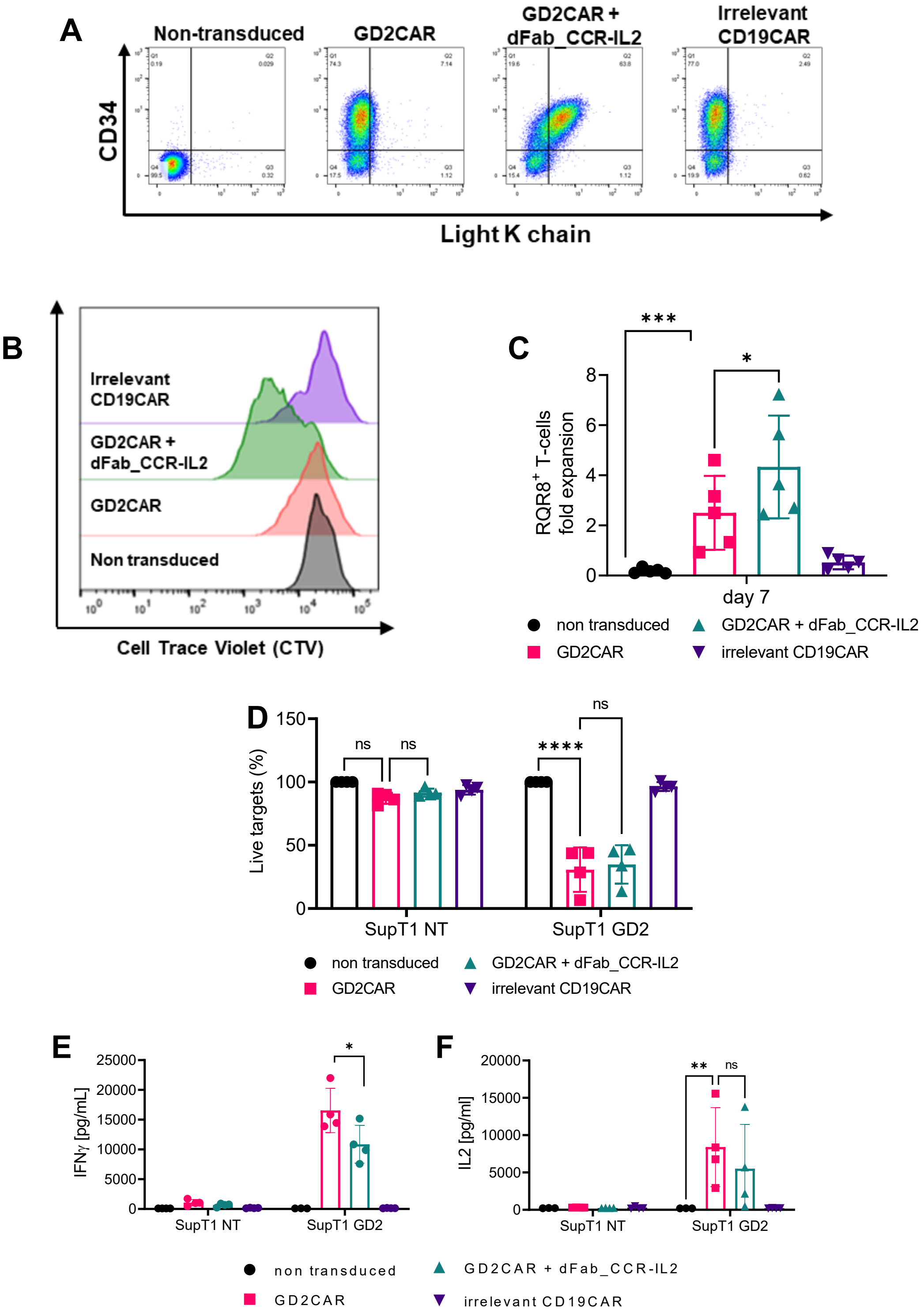

**Supplementary Figure S7. dFab_CCR-IL2 maintains functionality when co-expressed with a second generation GD2 CAR in a bi-cistronic vector.**

**(A)** Representative dot plot of the flow cytometry evaluation of T cells transduced with GD2 CAR, GD2 CAR co-expressing dFab_CCR-IL2 or irrelevant CD19 CAR. The graph represents

CD34 and light κ chain expression. **(B)**. Representative histogram of Cell Trace Violet (CTV) dilution of non-transduced, CAR only or CAR co-expressing dFab_CCR-IL2 after 96h of cytokine starvation. **(C)** Proliferation of either GD2 CAR alone or GD2CAR co-expressing dFab_CCR-IL2 T cells cultured for 7 days in cytokine starvation. Specific proliferation expressed as fold expansion (right) of $RQR8^{+}$ / $CD3^{+}$ T cells (n=4, one way ANOVA), **(D)** Killing of SupT1-NT or SupT1-$GD2^{+}$ after 48 hours co-culture with GD2 CAR alone or co-expressing dFab_CCR-IL2 T cells at a 1:1 effector:target ratio. Data shows mean percentage (± SD) of live cells compared to non-transduced (NT) control (n=4, two-way ANOVA). **(E, F)** Quantification of IFNγ (E) or IL2 (F) release from D. Data shows mean percentage (± SD) (n=4, two-way ANOVA). All data are presented as mean ± SEM.

#### SUPPLEMENTARY FIGURE S8

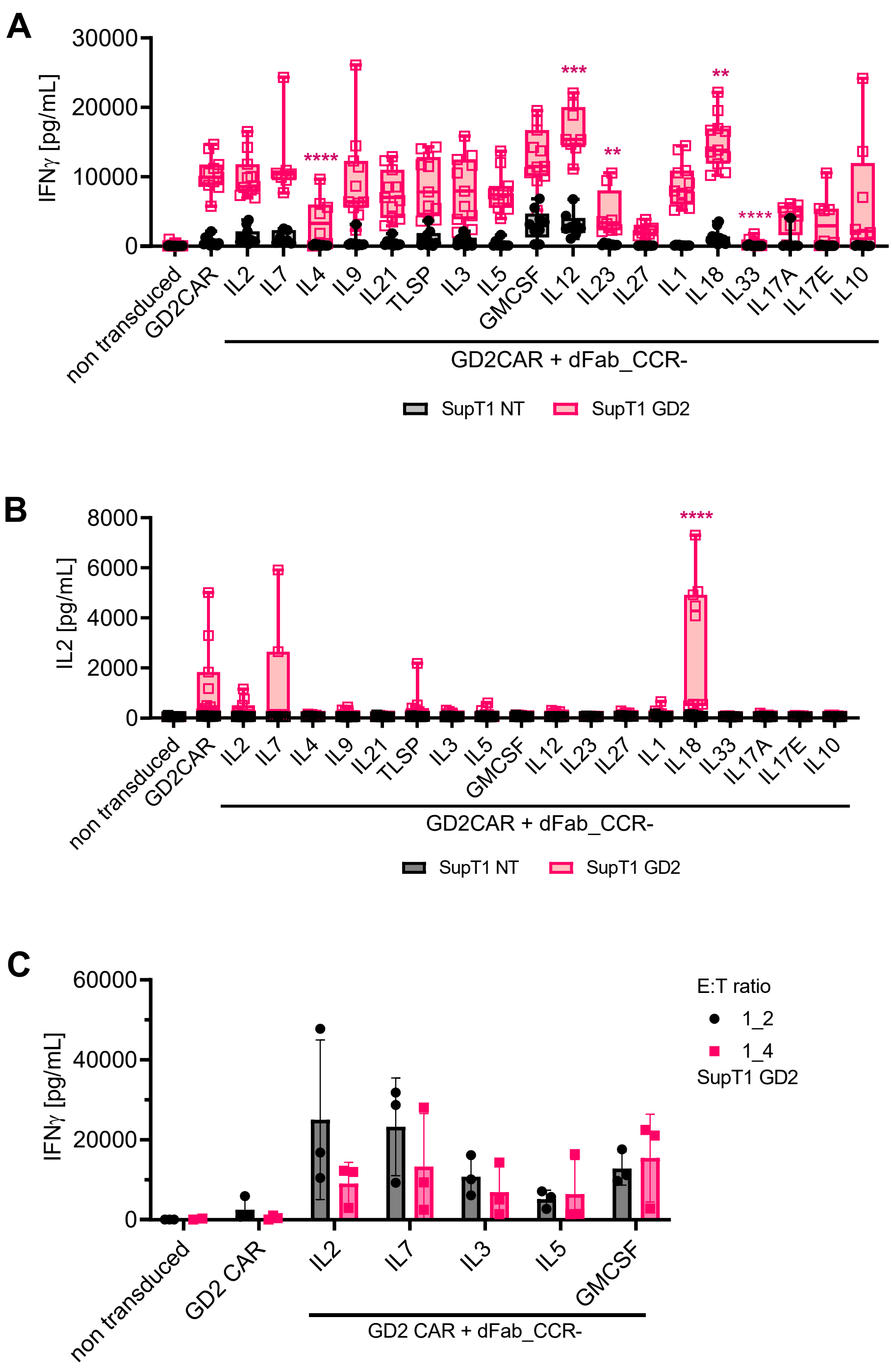

**Supplementary Figure S8. dFab_CCRs co-expression protect GD2 CAR T cells cytokine secretion.**

**(A, B)** IFNγ (A) and IL2 (B) secretion evaluated from co-culture experiment in Figure 3C. (n=4, two-way ANOVA). All data are presented as mean ± SEM. **(C)** IFNγ (A) and IL2 (B) secretion evaluated from co-culture experiment in Figure 3C Figure 3E (n=4, two-way ANOVA).

#### SUPPLEMENTARY FIGURE S9

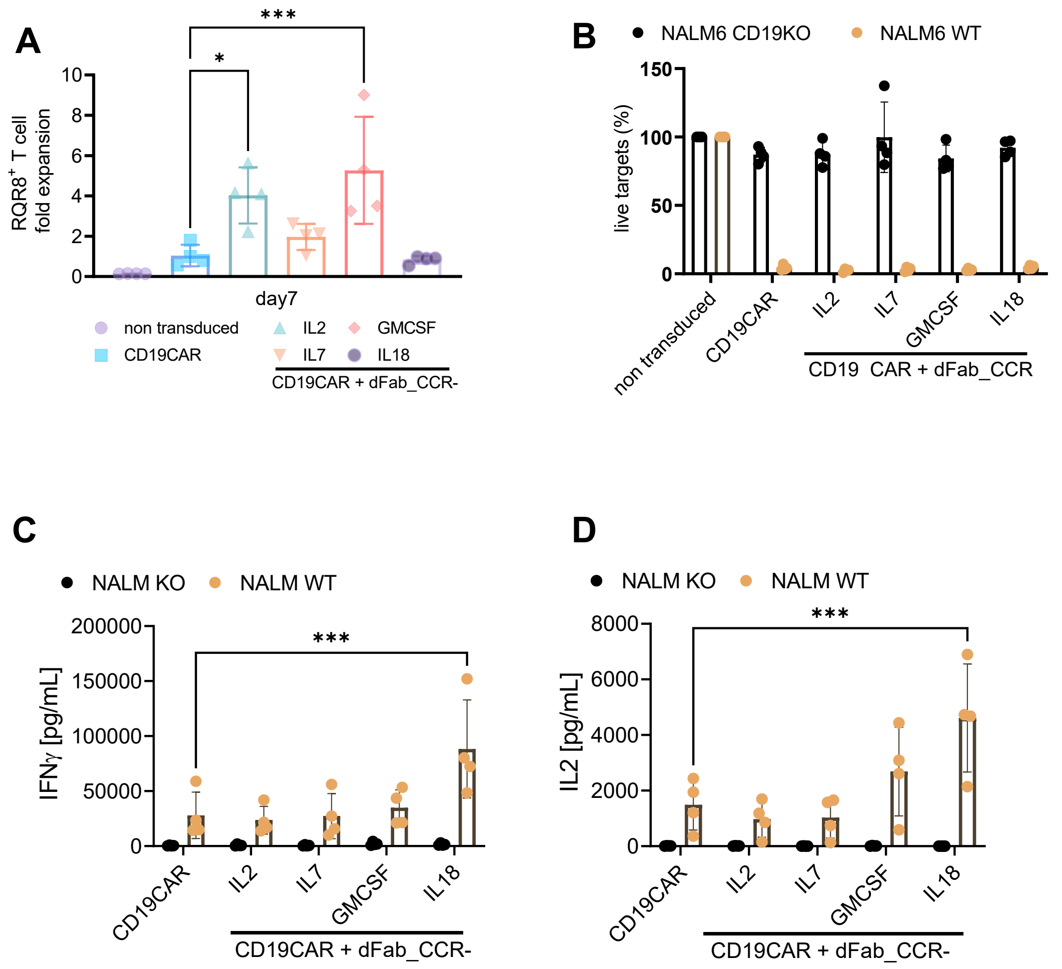

**Supplementary Figure S9. Characterisation of selected dFab_CCRs co-expressed with a second generation 41bbζ-CD19 CAR.**

**(A)** Proliferation of either CD19 CAR alone or CD19CAR co-expressing dFab_CCR-IL2, IL7, GMCSF or IL18 cultured for 7 days in cytokine starvation. Specific proliferation expressed as fold expansion (right) of RQR8$^+$/CD3$^+$ T cells (n=4, one way ANOVA). **(B)** Killing of NALM6 CD19KO (black) or NALM6 WT (orange) after 48 hours co-culture with GD2 CAR alone or co-expressing dFab_CCR-IL2 T cells at 1:4 effector:target ratio. Data shows mean percentage (± SD) of live cells compared to non-transduced (NT) control (n=4). **(C, D)** Quantification of IFNγ (C) or IL2 (D) release from (B). (n=4, two-way ANOVA). All data are presented as mean ± SEM.

**SUPPLEMENTARY FIGURE S10**

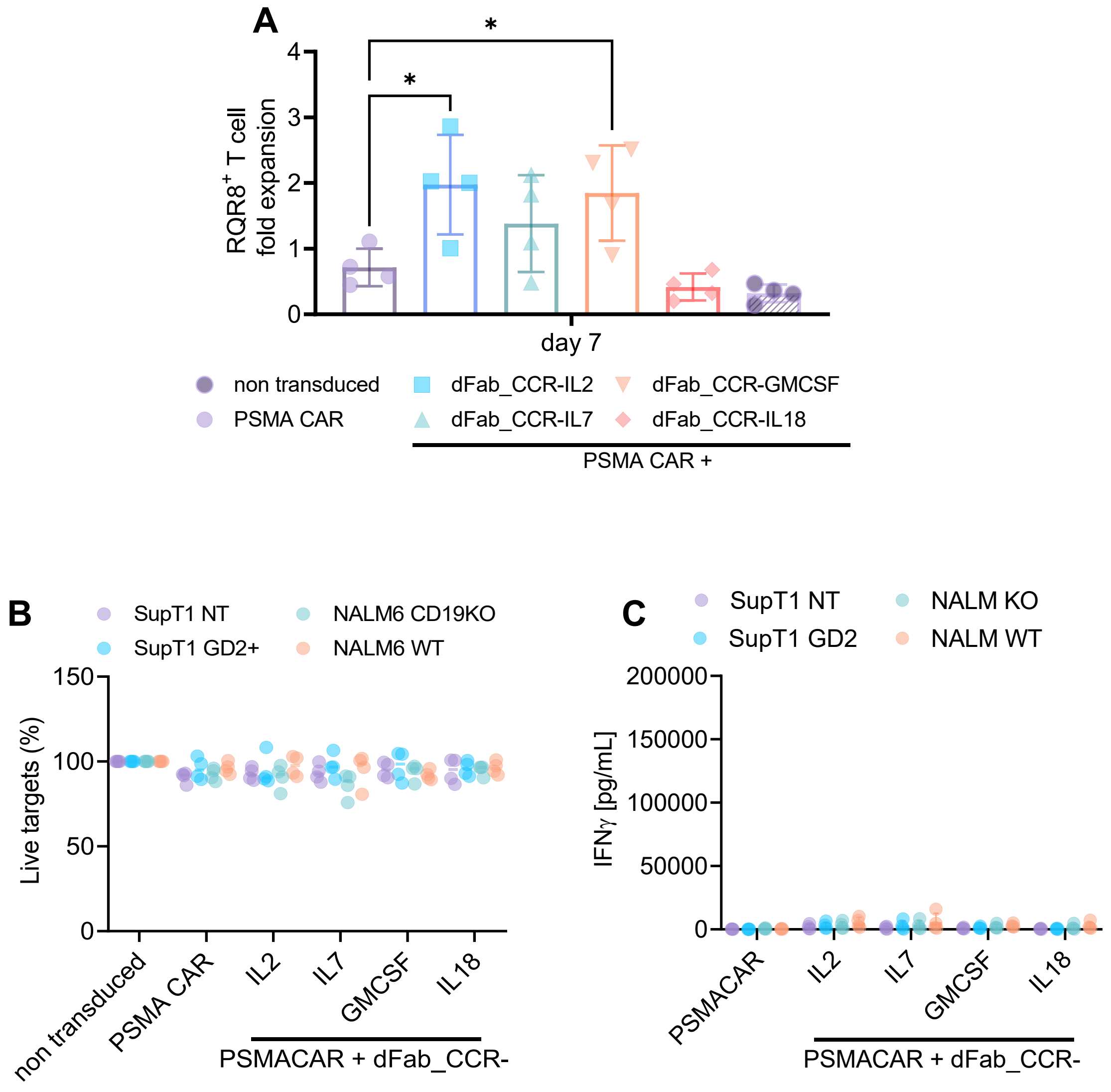

**Supplementary Figure S10. dFab_CCRs co-expression do not alter CAR specificity.**

**(A)** Proliferation of either PSMA CAR alone or PSMACAR co-expressing dFab_CCR-IL2, IL7, GMCSF or IL18 cultured for 7 days in cytokine starvation. Specific proliferation expressed as fold expansion of $RQR8^{+}$ / $CD3^{+}$ T cells (n=4, one way ANOVA), **(B)** Killing of SupT1 NT (purple), SupT1 GD2 (cyan), NALM6 CD19KO (green) or NALM6 WT (orange) after 48 hours co-culture with GD2 CAR alone or co-expressing dFab_CCR-IL2 T cells at 1:4 effector:target ratio. Data shows mean percentage (± SD) of live cells compared to non-transduced (NT) control (n=4). **(C)** Quantification of IFNγ release from B. (n=4). All data are presented as mean ± SEM.

#### SUPPLEMENTARY FIGURE S11

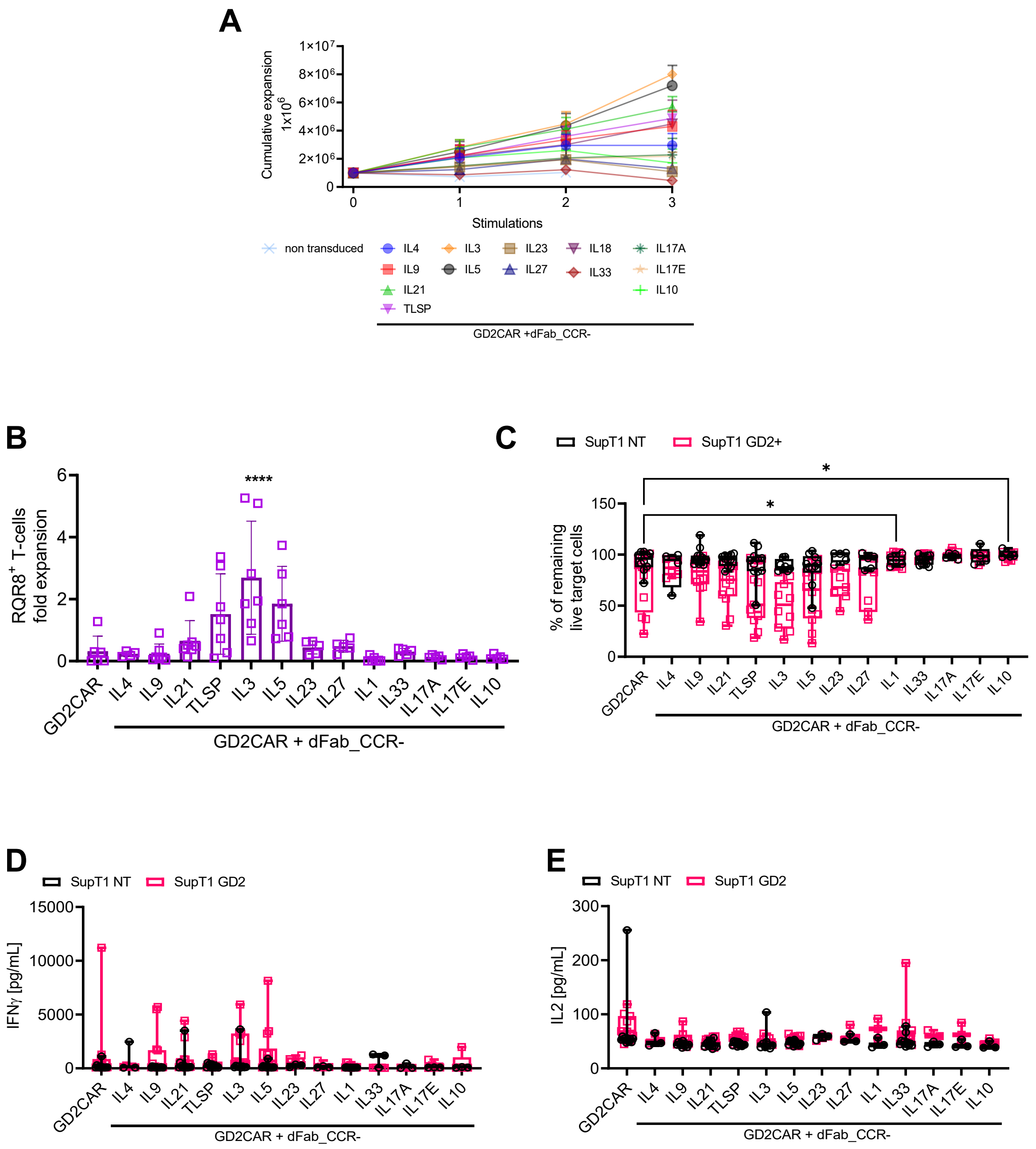

**Supplementary Figure S11. Differential effect on GD2 CAR functionality of the remaining dFab_CCRs library receptor upon antigen chronic stimulation.**

**(A)** Expansion of GD2 CAR T cells co-expressing dFab_CCRs. Data are expressed as $10^6$ cells cumulative expansion (n=6). **(B)** Quantitated *in vitro* proliferation of dFab_CCRs co-expressing GD2 CAR T cells, recovered after three rounds of stimulation (A) and cultured for 7 days in cytokine starvation condition. Proliferation expressed as fold expansion of RQR8$^{+}$ / CD3$^{+}$ GD2 CAR T cells (n=6, one-way ANOVA). **(C)** Killing of SupT1 NT (black) and GD2$^{+}$ (red) after 48 hours co-culture with GD2 CAR-T cells recovered after three rounds of co-culture (A) at 1:4 effector:target ratio. Data shows mean percentage (± SD) of live cells compared to non-transduced (NT) control (n=6, two-way ANOVA). **(D, E)** IFNγ (D), IL2 (E) secretion from (C) (n=6,). All data are presented as mean ± SEM.

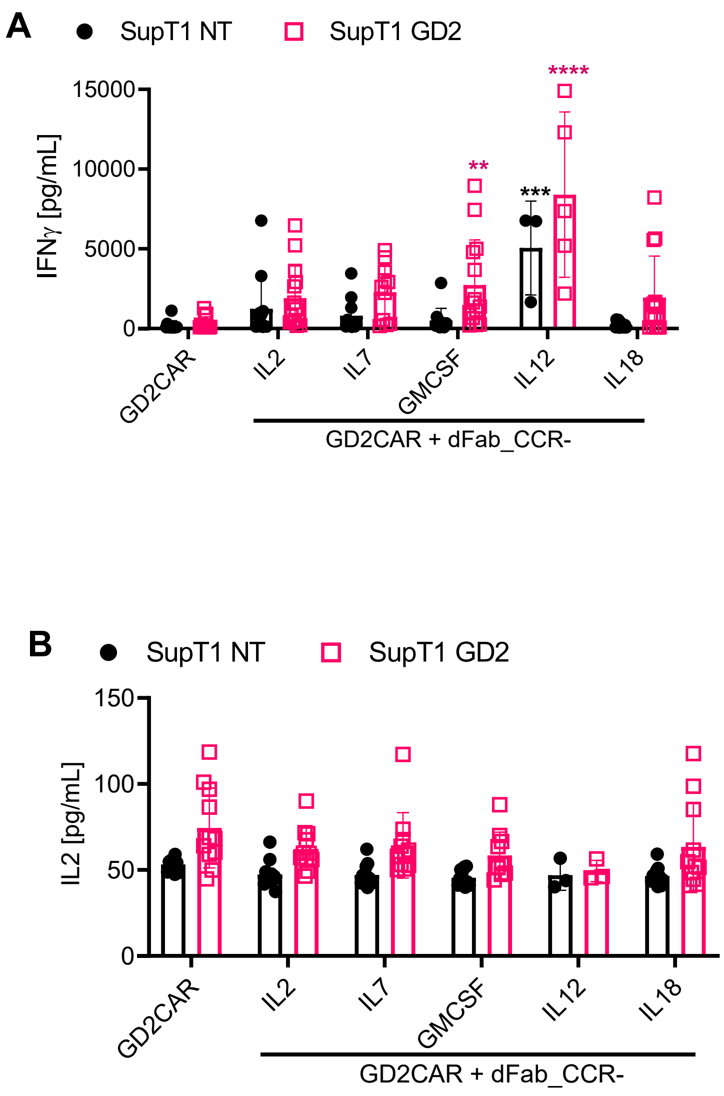

**Supplementary Figure S12. Selected dFab_CCRs library receptors protect GD2 CAR T cells cytokine secretion upon antigen chronic stimulation.**

**(A, B)** Quantification of IFNγ (A) or IL2 (B) release from co-culture in Figure 4D (n=4, two-way ANOVA). All data are presented as mean ± SEM.

#### SUPPLEMENTARY FIGURE S13

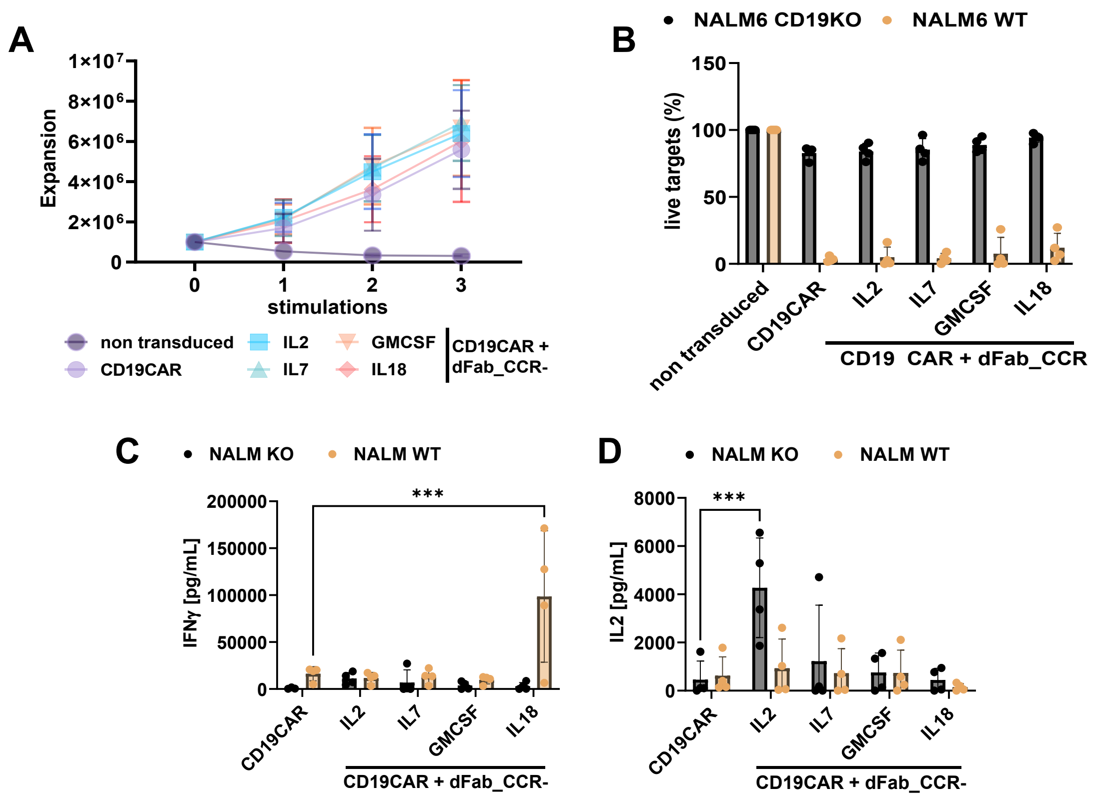

**Supplementary Figure S13. Chronic Antigen exposure effects on CD19 CAR T cells co-expressing selected dFab_CCRs.**

**A)** Expansion of GD2 CAR T cells co-expressing dFab_CCRs. Data are expressed as $10^6$ cells cumulative expansion (n=4). **(B)** Killing of NALM6-CD19KO (black) and NALM6 WT (orang) after 48 hours co-culture with cells recovered after three rounds of co-culture (A). at 1:4 effector:target ratio. Data shows mean percentage (± SD) of live cells compared to non-transduced (NT) control (n=4). **(C, D)** IFNγ (C), IL2 (D) secretion from (C) (n=4, two-way ANOVA).

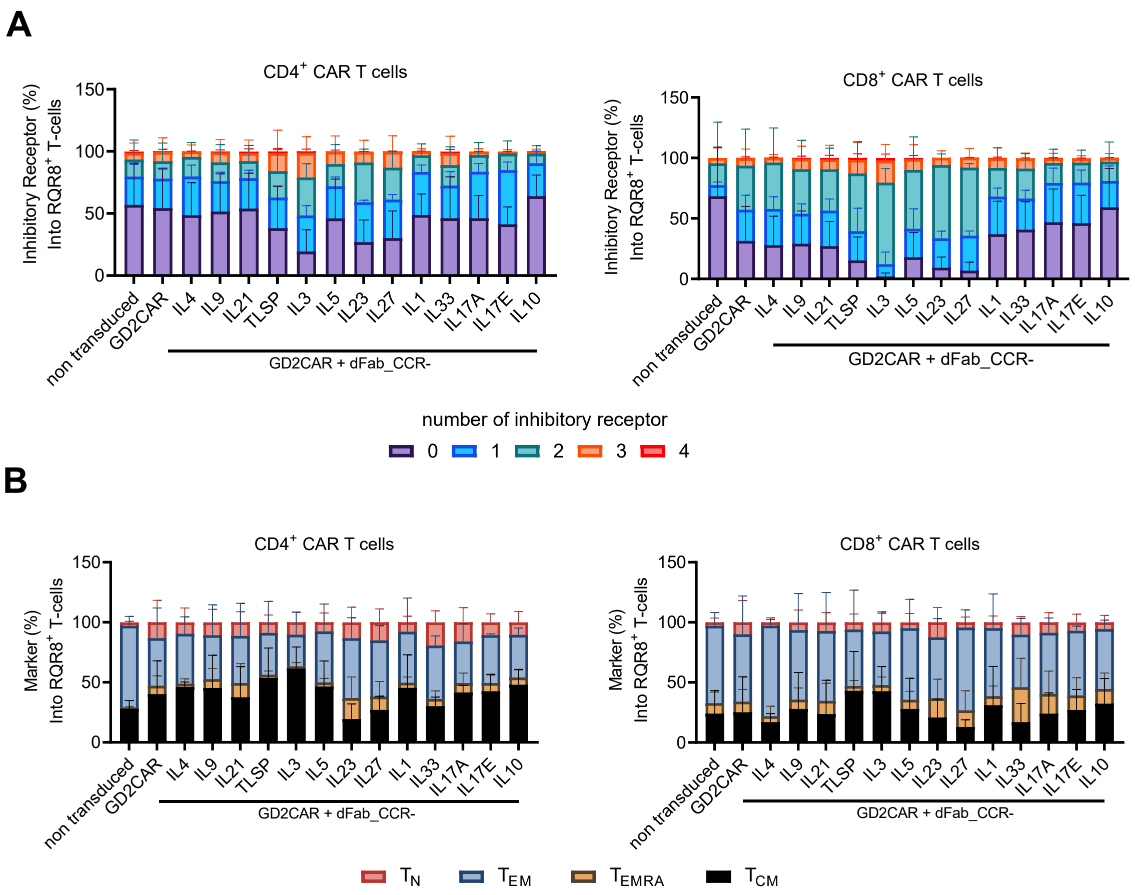

**Supplementary Figure S14. Differential effect of the remaining dFab_CCRs library receptors on exhaustion and memory phenotype after chronic antigen exposure.**

**(A)** Exhaustion phenotype of CAR T cells after three rounds of chronic antigen exposure, from Supp Fig S13A. Exhaustion evaluated by the expression of TIM3, LAG3, PD-1 or KLRG1 in either $CD4^+$ (left) or $CD8^+$ (right) T cells. Stacked bars show percentage of cells expressing 0, 1, 2, 3 or 4 markers per individual donors (n=11, one-way ANOVA). All data are presented as mean ± SEM. **(B)** Memory phenotype of CAR T cells after three rounds of chronic antigen exposure, from Supp Fig. S13A. Memory phenotype evaluated by the expression of CD45RA and/or CCR7 in either $CD4^+$ (left) or $CD8^+$ (right) T cells. Memory phenotype defined as: naïve T cells ($CD45RA^+$, $CCR7^+$), central memory (CM) T cells ($CD45RA^-$, $CCR7^+$), effector memory (EM) T cells ($CD45RA^-$, $CCR7^-$) and terminally differentiated (TEMRA) T cells ($CD45RA^+$, $CCR7^-$). Stacked bars show percentage of cells in each population markers per individual donor (n=11, one-way ANOVA). All data are presented as mean ± SEM.

#### SUPPLEMENTARY FIGURE S15

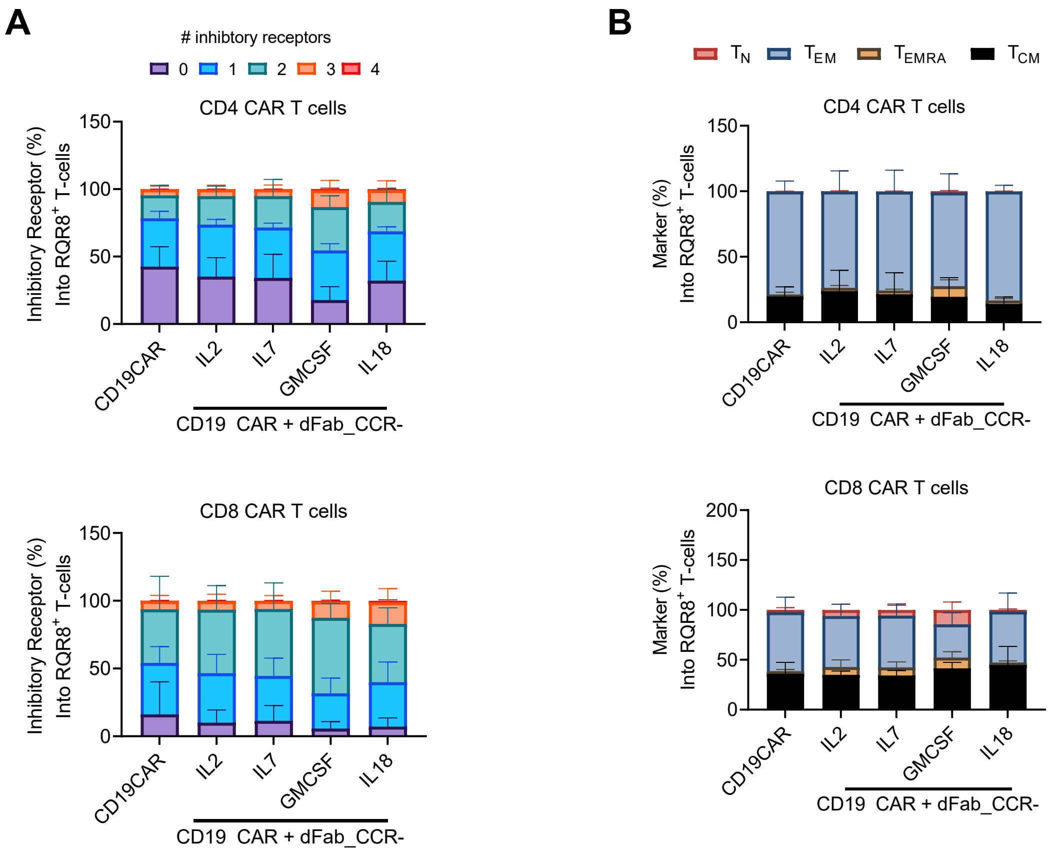

**Supplementary Figure S15. Differential effect of selected dFab_CCRs library receptors on exhaustion and memory phenotype after CD19 CAR T cell chronic antigen exposure.**

**(A)** Exhaustion phenotype of CAR T cells after three rounds of chronic antigen exposure, from A. Exhaustion evaluated by the expression of TIM3, LAG3, PD-1 or KLRG1 in either

$CD4^+$ (top) or $CD8^+$ (bottom) T cells. Stacked bars show percentage of cells expressing 0, 1, 2, 3 or 4 markers per individual donors (n=4, one-way ANOVA). All data are presented as mean ± SEM. **(B)** Memory phenotype of CAR T cells after three rounds of chronic antigen exposure, from A. Memory phenotype evaluated by the expression of CD45RA and/or CCR7 in either $CD4^+$ (top) or $CD8^+$ (bottom) T cells. Memory phenotype defined as: naïve T cells ($CD45RA^+$, $CCR7^+$), central memory (CM) T cells ($CD45RA^-$, $CCR7^+$), effector memory (EM) T cells ($CD45RA^-$, $CCR7^-$) and terminally differentiated (TEMRA) T cells ($CD45RA^+$, $CCR7^-$). Stacked bars show percentage of cells in each population markers per individual donors (n=4, one-way ANOVA). All data are presented as mean ± SEM.

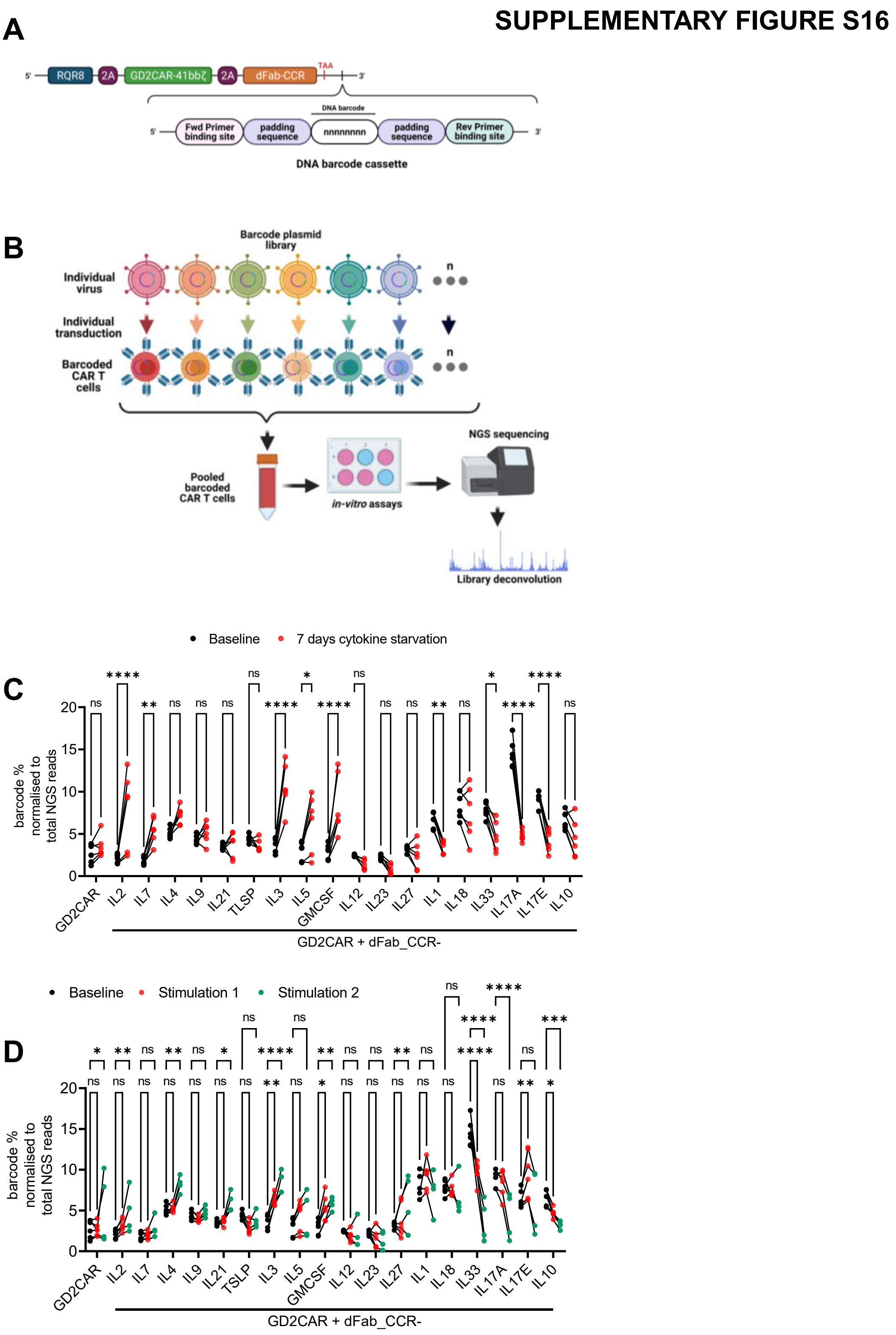

**Supplementary Figure S16. dFab_CCR DNA barcoded dFab_CCR library deconvolute individual dFab_CCR functionality.**

**(A)** Schematic of the Barcode DNA cassette. The individual DNA barcode is surrounded by two padding sequences and the primer binding sites**.** **(B)** Experimental set-up of *in vitro* and of the barcoded GD2CAR co-expressing dFab_CCRs pooled library. **(C)** Representation of individual barcoded dFab_CCR CAR T cells within the mix population. The graph represents the barcode frequency at day 0 (baseline, black) and after 7 days of cytokine starvation (red) (n=6, two-way ANOVA). **(D)** Representation of individual barcoded CCRs within the mix population after two rounds of serial co-culture. The graph represents the barcode frequency at day 0 (baseline, black), after one stimulation (red) and after the second stimulation (green) (n=6, two-way ANOVA).

### SUPPLEMENTARY FIGURE S17

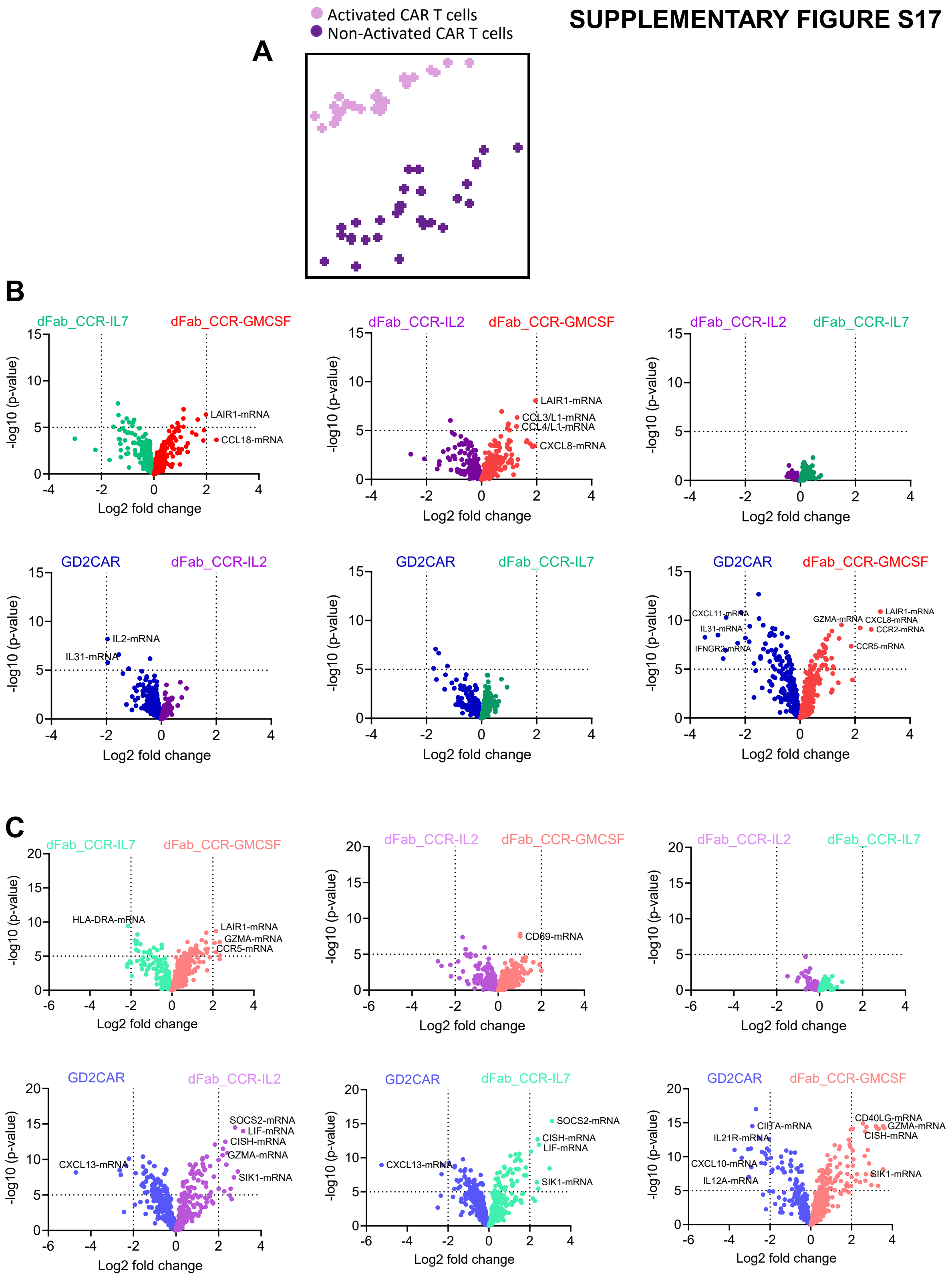

**Supplementary Figure S17. Transcriptomic profile of GD2 CAR T cells co-expressing a selection of dFab_CCRs.**

**(A)** PCA analysis of transcriptome differences in activated CAR T cells (pink) or non-activated CAR T cells (purple) (n=6). **(B, C)** Differential gene expression volcano plot comparisons in activated CAR T cells (B) or non-activated CAR T cells (C) (n=6)

#### SUPPLEMENTARY FIGURE S18

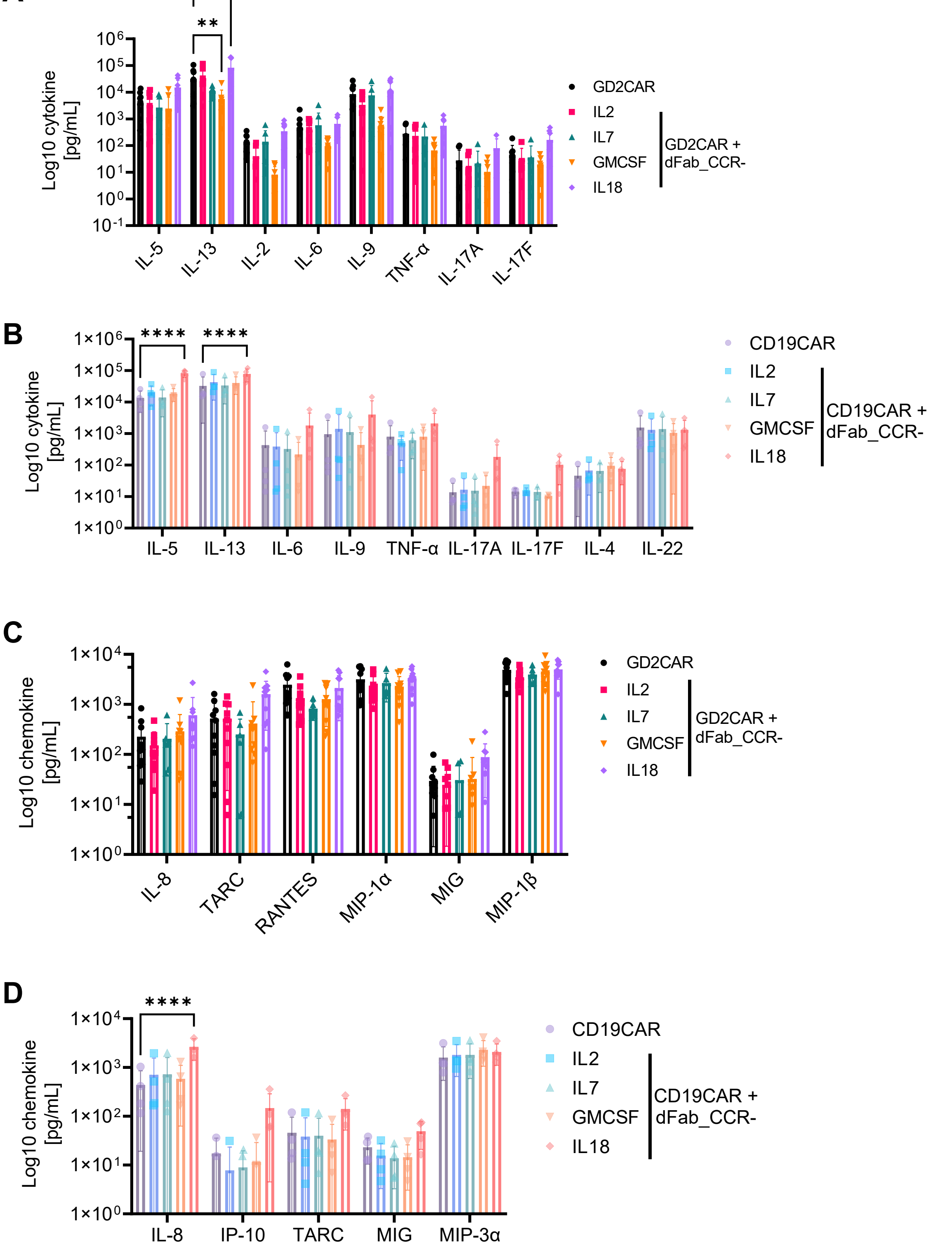

**Supplementary Figure S18. Chemokines and cytokines secretion profile of GD2 and CD19 CAR T cells co-expressing a selection of dFab_CCRs.**

**(A, B)** Quantitation of chemokines (A) or cytokines (B) of GD2CAR T cells co-expressing selected dFab_CCRs from co-culture with SupT1 GD2$^+$ target cells in Figure 3C, analyzed with cytokine beads array (n=9). **(D, E)** Quantitation of chemokines (D) or cytokines (E) of

CD19CAR T cells co-expressing selected dFab_CCRs from co-culture with NALM6 WT target cells in Supplementary Figure S10B analyzed with cytokine beads array (n=4)

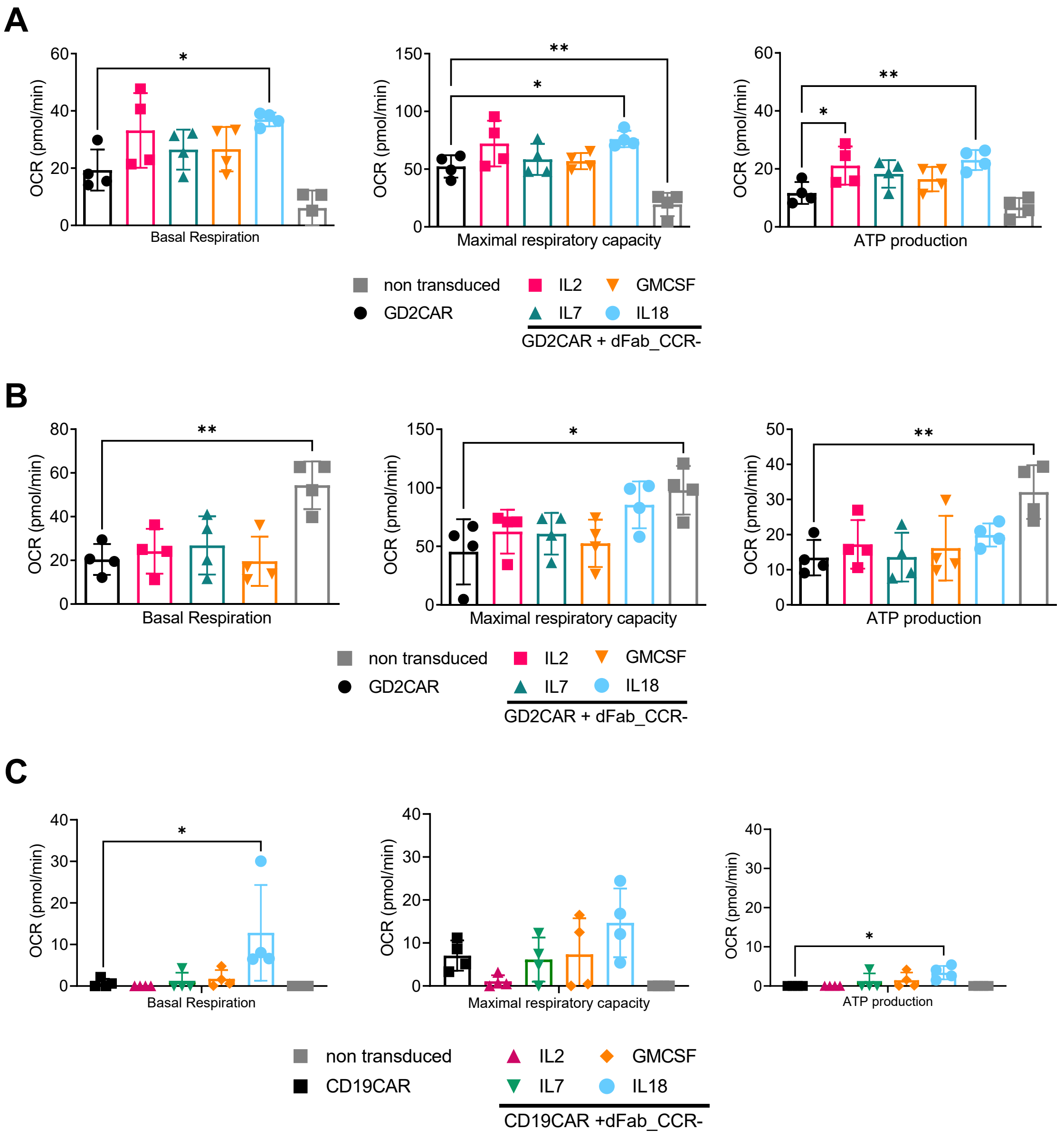

**Supplementary Figure S19. Metabolomic profile of GD2 and CD19 CAR T cells co-expressing a selection of dFab_CCRs.**

**(A, B)** Evaluation of basal respiration, maximal respiratory capacity, and ATP production in resting GD2 CAR T cells (A) and activated GD2 CAR T cells (B) (n=4). **(C)** Evaluation of basal respiration, maximal respiratory capacity, and ATP production in resting CD19 CAR T cells (n=4). All data are presented as mean ± SEM.

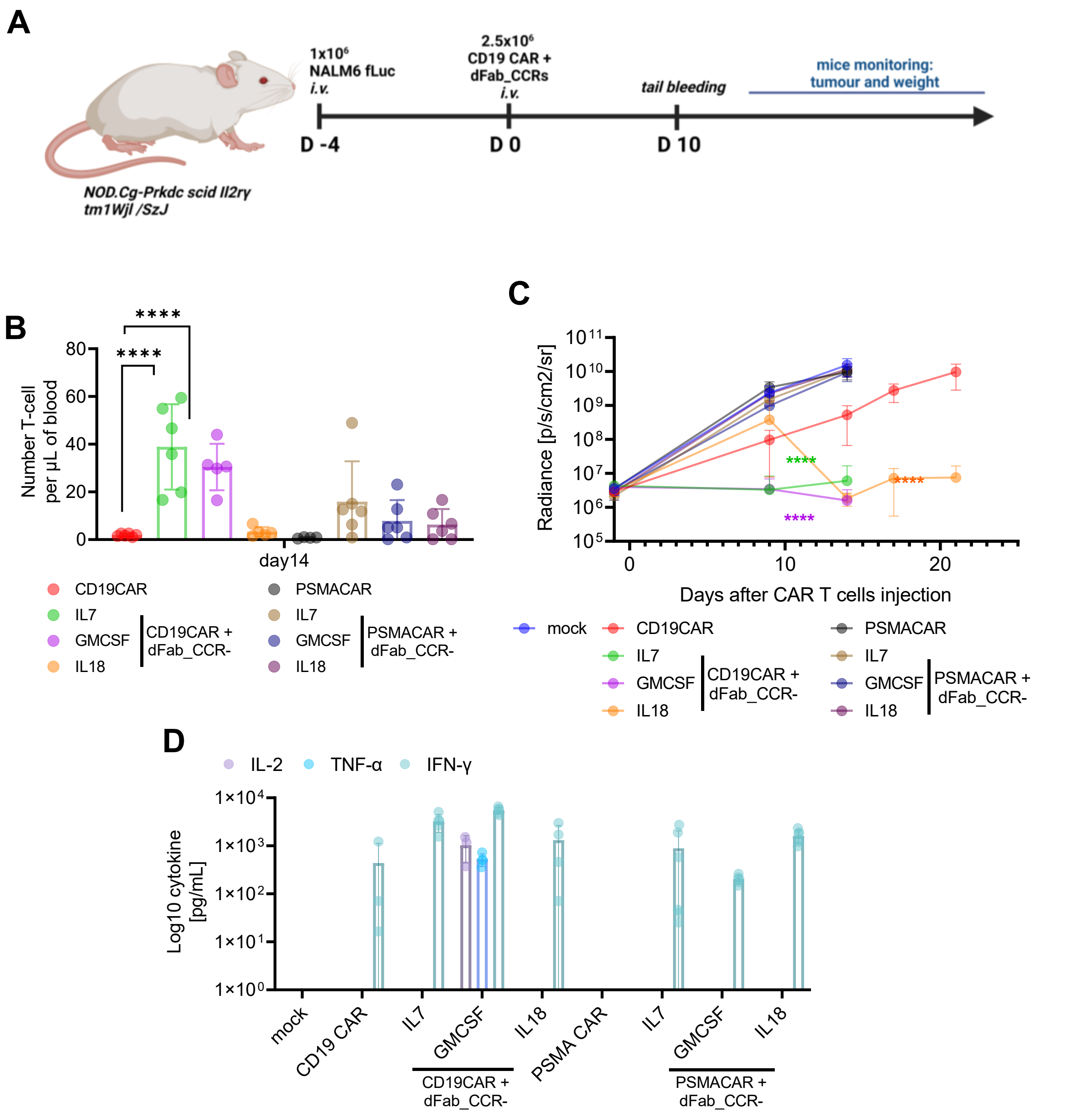

**Supplementary Figure S20. dFab_CCR co-expression augments CD19 CAR T cells efficacy in an establishes ALL NALM6 xenograph model.**

**(A)** Experimental *in vivo* setup of the established NALM-6 ALL xenograph model. **(B)** CD19 CAR T cells peripheral engraftment at day 14, quantitated via flow cytometry by the expression human CD3 (n=6, one-way ANOVA). **(C)** Bioluminescence analysis of total flux (n=6, day 14 student t-test, day 21 student t-test). **(D)** Quantitation of serum cytokines at day 10, analyzed with cytokine beads array (n=6). All data are presented as mean ± SEM.

#### SUPPLEMENTARY FIGURE S21

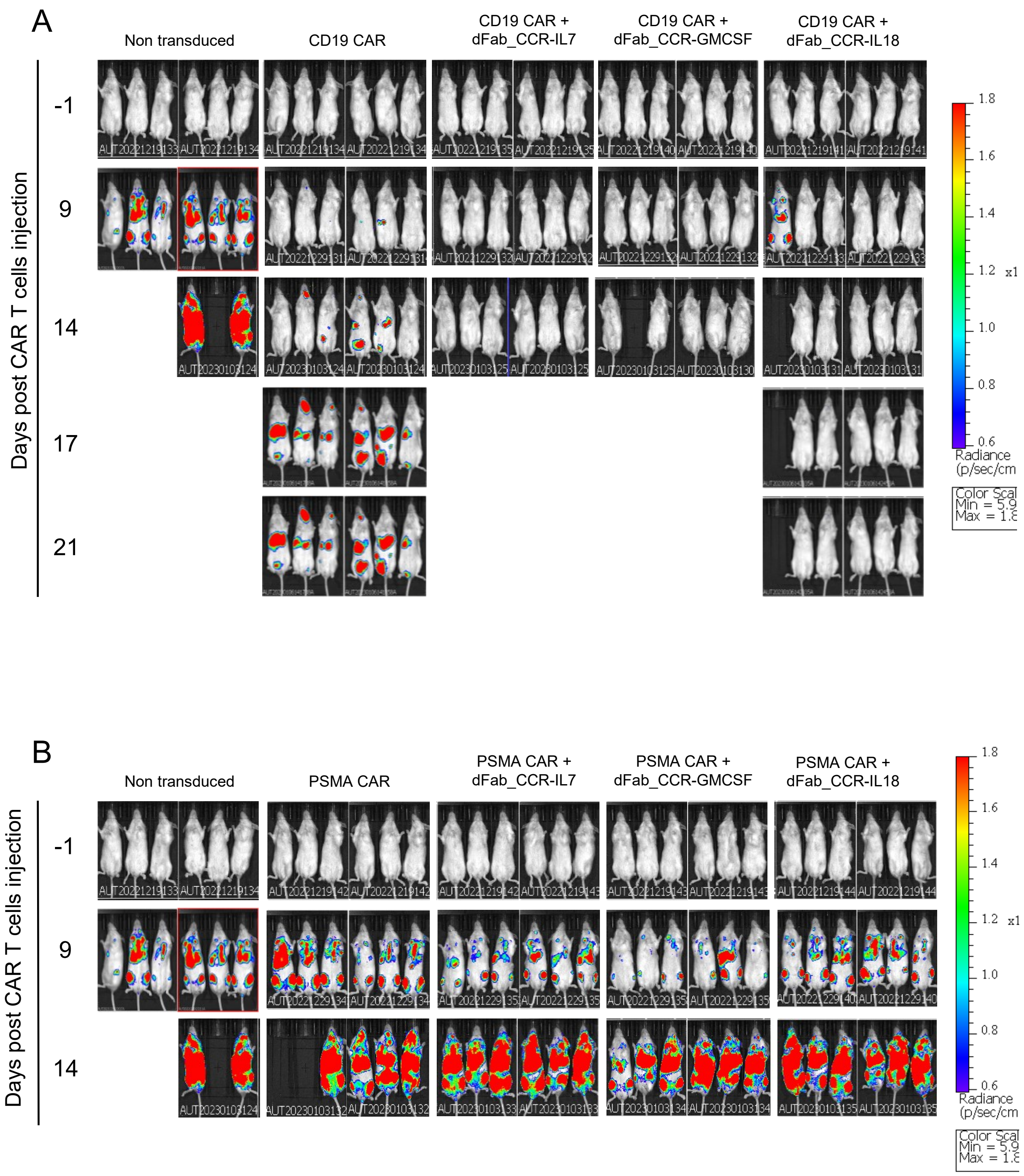

**Supplementary Figure S21. Bioluminescence imaging of the established ALL NALM6 xenograph model.**

**(A)** Bioluminescence imaging of CD19-CAR and CD19-CAR co-expressing dFab_CCR-IL7, GMCSF and IL18 in NALM6 xenograph model. **(B)** Bioluminescence imaging of PSMA-CAR and PSMA-CAR co-expressing dFab_CCR-IL7, GMCSF and IL18 in NALM6 xenograph model.

### SUPPLEMENTARY FIGURE S22

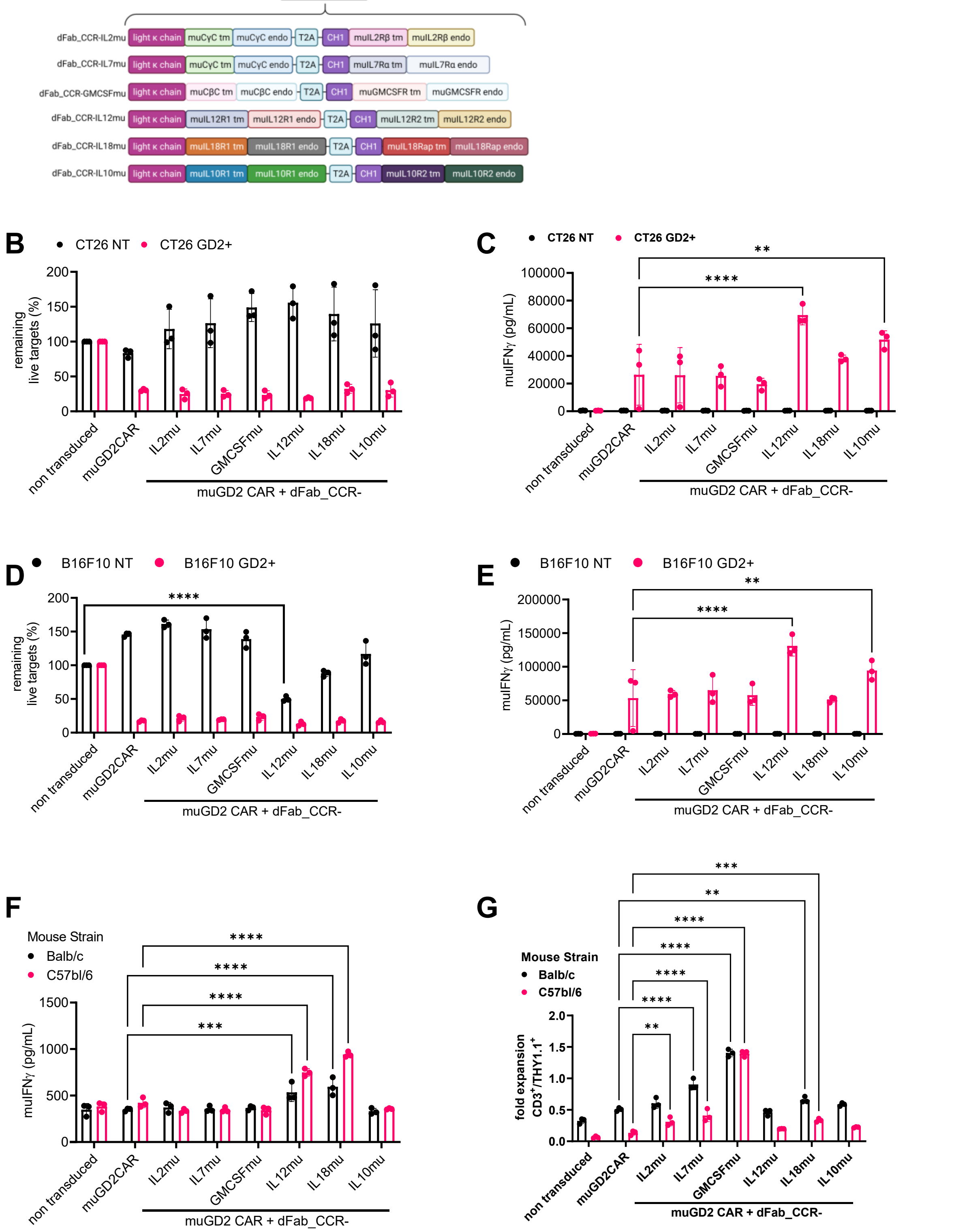

**Supplementary Figure S22.** ***In vitro*** **functionality of murinized dFab_CCR co-expressed with murine GD2CAR.**

**(A)** Schematic of the murine GD2 CAR plasmid DNA co-expressing selected murine optimized dFab_CCRs. **(B)** Killing of CT26-NT (black) and CT26-GD2$^{+}$ (red) after

48 hours co-culture with murine GD2 CAR T cells at 1:4 effector:target ratio. Data shows mean percentage (± SD) of live cells compared to non-transduced (NT) control (3 technical replicates, two-way ANOVA). **(C)** IFNγ secreted by co-culture from (A) (3 technical replicates, two-way ANOVA). **(D)** Killing of B16F10-NT (black) and B16F10-$GD2^+$ (red) after 48 hours co-culture with murine GD2 CAR T cells at 1:4 effector:target ratio. Data shows mean percentage (± SD) of live cells compared to non-transduced (NT) control (3 technical replicates, two-way ANOVA). **(E)** IFNγ secreted by co-culture from (C) (3 technical replicates, two-way ANOVA). **(F)** IFNγ secreted by murine GD2 CAR T cells culture for 48 hours in cytokine-free media (3 technical replicates, two-way ANOVA). **(G)** Quantitated *in vitro* proliferation of murine GD2 CAR T cells cultured for 96 hours in cytokine starvation condition. Proliferation expressed as fold expansion of $THY1.1^+$ / $CD3^+$ murine GD2 CAR T cells (3 technical replicates, two-way ANOVA). All data are presented as mean ± SEM.

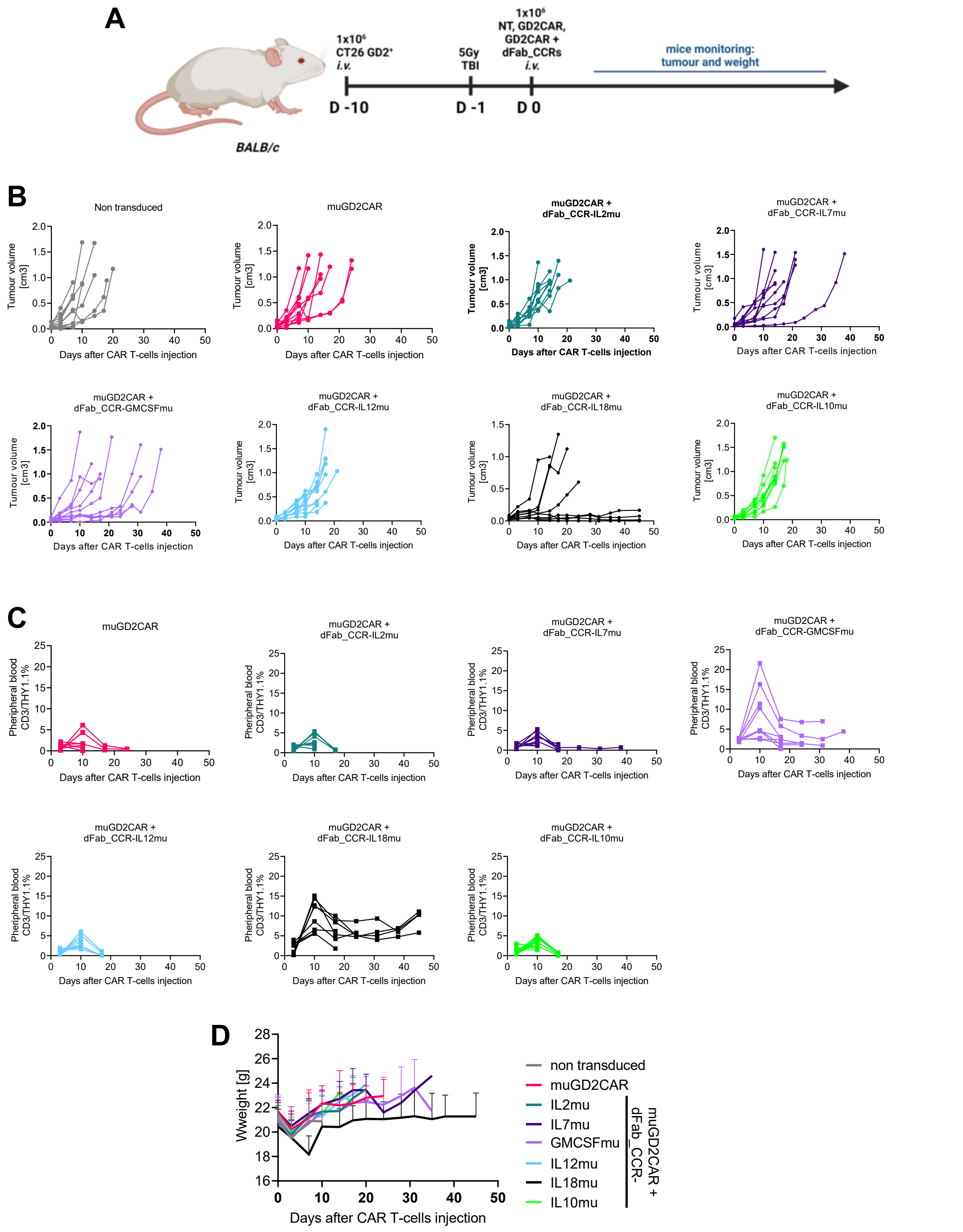

**Supplementary Figure S23. *In vivo* functionality of murinized dFab_CCR co-expressed with murine GD2CAR in syngeneic CT26-GD2 colon carcinoma model.**

(**A**) Experimental *in vivo* setup of the syngeneic CT26 GD2⁺ colon carcinoma model timeline, testing murine GD2 CAR T cells alone or co-expressing dFab_CCR-IL2mu, -IL7mu, -

GMCSFmu, -IL12mu, -IL18mu or -IL10mu. **(B)** Individual tumor growth traces ($mm^3$) (n=8) **(C)** Individual mice CAR T cell peripheral blood engraftment, evaluate as expression of transduction marker THY1.1 in $CD3^+/CD45^+/CD11b^-$ cells (n=8). All data are presented as mean ± SEM. **(D)** Mice weight measured bi-weekly and used as surrogate of CAR related toxicity (n-8).

#### SUPPLEMENTARY FIGURE S24

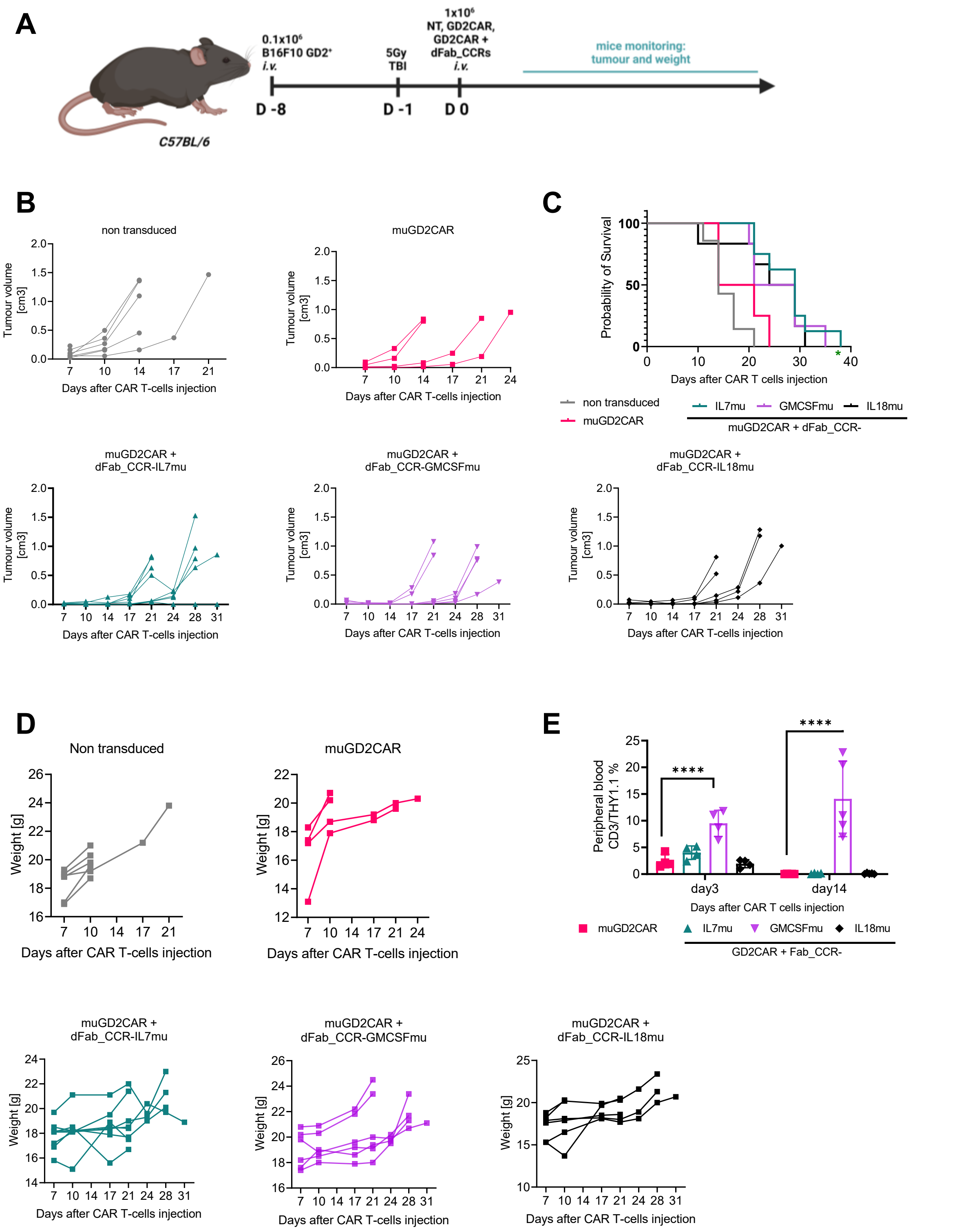

**Supplementary Figure S24.** ***In vivo*** **functionality of murinized dFab_CCR co-expressed with murine GD2CAR in syngeneic B16-GD2 melanoma model.**

**(A)** Experimental *in-vivo* setup of the syngeneic B16F10 GD2$^{+}$ melanoma model timeline, testing murine GD2 CAR T cells alone or co-expressing dFab_CCR-IL2mu, -IL7mu, -

GMCSFmu, -IL12mu, -IL18mu or -IL10mu. **(B)** Individual tumor growth traces (n=8). **(C)** Kaplan Mayer survival curve. **(D)** Individual mouse weight traces (n=8). **(E)** CAR T cell peripheral blood engraftment, evaluate as expression of transduction marker THY1.1 in $CD3^+$/$CD45^+$/$CD11b^-$ cells (n=8).

### SUPPLEMENTARY FIGURE S25

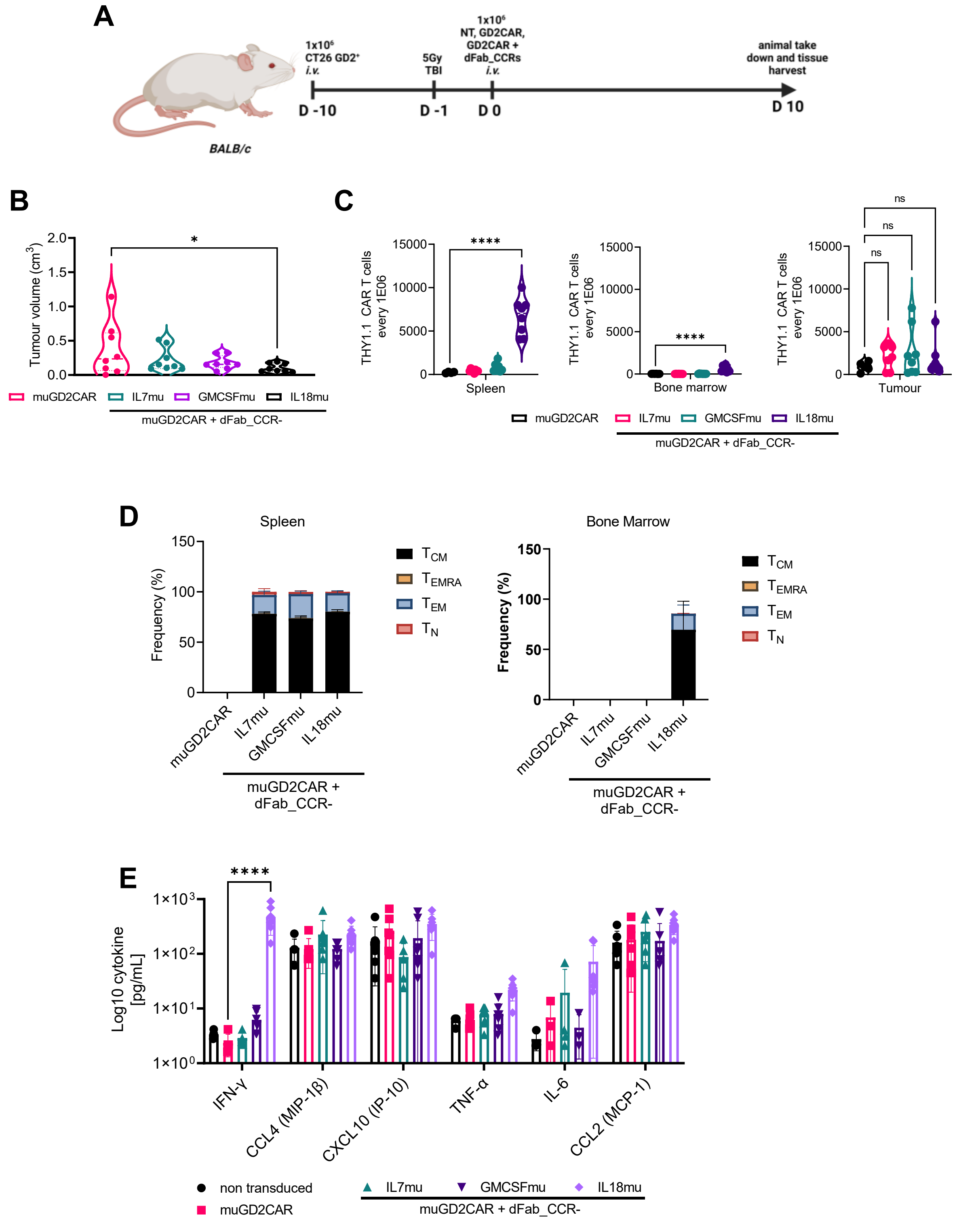

**Supplementary Figure S25. dFab_CCR-IL18 co-expression improved CAR T cells engraftment in lymphoid tissues.**

**(A)** Experimental timeline. **(B)** Tumour volume at day 10 (n=8, one-way ANOVA). **(C)** Murine GD2 CAR T cells engraftment in bone marrow, spleen and tumor expressed as THY1.1$^{+}$ CAR

T cell number every 1E06 $CD3^{+}$/$CD45^{+}$/$CD11b^{-}$ T cells (n=8, one-way ANOVA). **(D)** Memory phenotype of murine THY1.1 CAR T cells in the bone marrow and spleen. Memory phenotype evaluated by the expression of CD62L and CD44. Naïve T cells ($C44^{-}$, $CD62L^{+}$), effector memory (EM) T cells ($CD44^{+}$, CD62L+), Central Memory (CM) T cells (CD44+, $CD62L^{+}$) and terminally differentiated (TEMRA) T cells ($CD44^{-}$, $CD62L^{-}$). Stacked bars show the percentage of cells in each population markers per individual donor (n=8) All data are presented as mean ± SEM. **(E)** Quantification of serum chemokines and cytokines (n=8, two-way ANOVA). All data are presented as mean ± SEM.
