## Supplementary Table 1 for "Enhancing CAR T cell therapy using Fab-Based Constitutively Heterodimeric Cytokine Receptors"

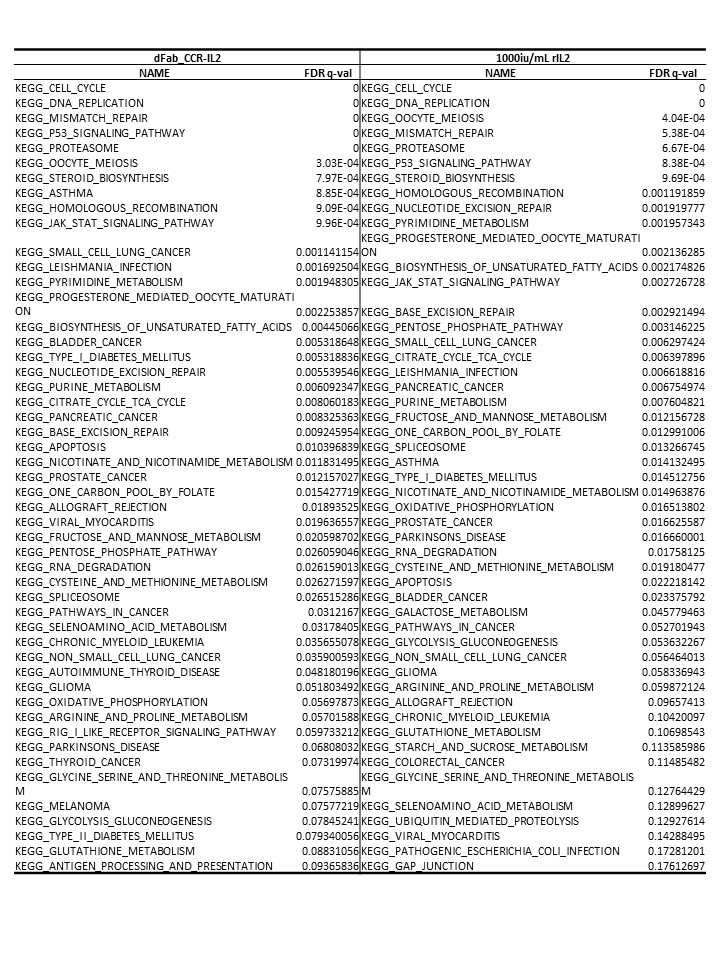
Supplementary Table S1. Top 50 up-regulated GSEA pathways.

Top 50 up-regulated GSEA pathways utilised for the generation of the Venn diagram in figure 1H. dFab_CCR-IL2 T cells or non-transduced T cells treated with 100000 iU/mL IL2, were compared to non-transduced T cells culture in cytokine starvation.
